# Integrative genetic and transcriptomic analysis identifies candidate genes and compounds for Fragile X syndrome and autism spectrum disorder

**DOI:** 10.64898/2026.10.03.756403

**Authors:** Wen Zhang, Jingqi Yan, Michael DeGiorgio

## Abstract

Fragile X syndrome (FXS) is the most commonly known monogenic cause of autism spectrum disorder (ASD), yet how FXS biology relates to polygenic architecture of idiopathic ASD remains poorly defined. We integrated publicly available genetic data relevant to ASD and FXS, including population case-control risk, symptom discordance within monozygotic twin pairs, contrasts between ASD and other psychiatric disorders, and genetic variation associated with circulating plasma fragile X messenger ribonucleoprotein (FMRP) abundance. These data were used to identify associated genes and pathways across tissues and prioritize small molecules predicted to reverse genetically associated transcriptional signatures. Cross-tissue analysis identified *XRN2* as the strongest association for population ASD risk and *PPP3CA* as the strongest association with FMRP abundance, providing distinct candidate links to ASD and FXS biology. We found no evidence that population ASD risk and symptom discordance within monozygotic twin pairs share common-variant architecture after accounting for linkage disequilibrium, suggesting that these phenotypes capture at least partially distinct sources of genetic signal. Connectivity mapping identified candidate compounds across eight genetic signatures, including several with biologically plausible mechanisms. Together, these findings provide a genetics-anchored framework for connecting FXS and ASD biology and for prioritizing candidate therapeutic mechanisms and compounds, while highlighting methodological safeguards needed for reliable integrative genomic drug repurposing.

## Introduction

Autism spectrum disorder (ASD) affects an estimated one in 36 children in the United States, which is characterized by impairments in social communication and restricted, repetitive patterns of behavior (Constantino & Charman, 2016; American Psychiatric Association, 2013). Twin and family studies place the heritability of ASD between 64% and 91%, and large-scale genome-wide association studies (GWAS) have established that common genetic variation contributes substantially to population-level ASD liability. Rare de novo and inherited variants of larger effect also contribute to ASD risk (Bourgeron, 2015), as identified through exome and genome sequencing (Grove et al., 2019; Satterstrom et al., 2020). The genetic architecture of ASD is therefore highly polygenic, with thousands of common variants of individually small effect combining with a smaller number of rare, higher penetrance variants (Grove et al., 2019; Satterstrom et al., 2020). Together, these forms of genetic variation contribute to the observed phenotypic spectrum, consistent with the broader omnigenic model proposed for complex traits (Boyle, Li, & Pritchard, 2017). Coexpression and single-cell transcriptomic studies have further implicated mid-fetal cortical projection neuron development as a point of convergence for genetically heterogeneous ASD risk genes (Willsey et al., 2013). These findings underscore the value of transcriptomic approaches for interpreting ASD genetic association signals.

Among the monogenic causes of ASD, Fragile X syndrome (FXS) is the most commonly known monogenic cause (Hagerman et al., 2017; Bagni & Zukin, 2019), with approximately 30% to 60% of individuals with FXS meeting diagnostic criteria for ASD (Hagerman et al., 2017; Bagni & Zukin, 2019). FXS results from a CGG repeat expansion in the 5’ untranslated region of the *FMR1* gene that leads to epigenetic silencing and loss of its encoded protein, fragile X messenger ribonucleoprotein (FMRP) (Darnell et al., 2011; Bagni & Zukin, 2019). FMRP is an RNA binding protein that regulates the translation of a defined set of synaptic mRNAs (Darnell et al., 2011). Its loss produces exaggerated group I metabotropic glutamate receptor (mGluR) dependent long-term depression (Contractor et al., 2015). This mechanistic link motivated the mGluR theory of FXS and led to several clinical trials targeting the mGluR5 subtype, which ultimately failed to demonstrate clinical efficacy (Contractor, Klyachko, & Portera-Cailliau, 2015; Darnell et al., 2011). The failure of mechanistically motivated mGluR5 directed compounds in FXS has prompted renewed interest in data driven approaches to therapeutic target and compound prioritization that are not constrained to a single a priori hypothesis (So et al., 2017; Corsello et al., 2017; Musa et al., 2018).

Despite the shared clinical phenotype of FXS and idiopathic ASD, the degree to which the monogenic biology of FXS and the polygenic architecture of idiopathic ASD converge at the level of gene expression, cellular pathway, or druggable targets remains poorly characterized. This gap is partly a consequence of data availability. No GWAS of FXS as a case-control trait exists because FXS is caused by a large effect CGG repeat expansion in *FMR1* rather than a spectrum of common variants that can be interrogated using standard GWAS designs. Population scale genetic information relevant to FXS biology is therefore only indirectly accessible through studies of circulating FMRP protein abundance. These protein quantitative trait locus (pQTL) studies capture common variant regulators of circulating FMRP abundance in the general population (Sun et al., 2018). Whether genetic variation associated with circulating FMRP abundance in the general population produces downstream transcriptomic signals that converge with genetic variation associated with population level ASD risk remains an open empirical question that the present study directly addresses.

Two complementary methodologies have emerged over the past decade to translate GWAS summary statistics into mechanistic and therapeutic hypotheses. First, transcriptome-wide association studies (TWAS), implemented through tools such as PrediXcan (Gamazon et al., 2015) and its summary statistic extension S-PrediXcan (Barbeira et al., 2018), impute the genetically regulated component of gene expression from GWAS summary statistics using tissue-specific expression prediction models trained on reference panels such as the Genotype-Tissue Expression (GTEx) project (GTEx Consortium, 2020). This approach converts SNP level association signals into gene and tissue level signals that are more directly interpretable and more readily linked to biological mechanisms. Because genetically regulated expression is inferred directly from genotype data using pretrained prediction models, TWAS does not require additional RNA sequencing or other transcriptomic profiling of the study cohort itself. Expression can instead be inferred for individuals with genome-wide genotype data or directly from GWAS summary statistics. This makes the approach applicable to large GWAS cohorts that would be prohibitively costly and logistically infeasible to profile transcriptomically at a comparable scale. The resulting increase in sample size can substantially improve power of gene level association testing relative to studies based on directly measured expression.

EpiXcan was subsequently developed to incorporate epigenetic information into transcriptome prediction models and improve prediction performance (Zhang et al., 2019). The study also incorporated a computational drug repurpose pipeline into the downstream analyses. Evaluating the genetically predicted expression of a single gene across dozens of tissues can help identify the tissue or cell context driving a genetic association. Association signals across tissues can then be integrated using a robust combination method that accounts for correlations among tissue predictors arising from shared regulatory variants and overlapping reference panels.

Second, connectivity mapping provides a complementary approach for translating disease associated transcriptional signatures into therapeutic hypotheses. Pioneered by the Connectivity Map (CMap) and subsequently scaled by the LINCS L1000 platform, connectivity mapping provides a reference library of transcriptional signatures induced by thousands of small molecule perturbagens across numerous cell lines (Lamb et al., 2006). This enables systematic identification of compounds whose transcriptional signatures are anti-correlated with a disease associated signature and therefore predicted to reverse the underlying molecular state (Subramanian et al., 2017; Zhang et al., 2019). Connectivity-based drug repurposing has been applied across oncology and inflammatory disease and, increasingly, to neuropsychiatric phenotypes. These analyses typically combine a disease associated differential expression signature derived from case-control transcriptomic profiling with the L1000 reference library (Subramanian et al., 2017; Duan et al., 2016; Musa et al., 2018). More recently, several studies have substituted TWAS imputed, genetically anchored expression signatures for directly measured differential expression (Peng & Zhang, 2025). This strategy avoids some of the confounding factors associated with measured brain transcriptomic data in psychiatric disease (So et al., 2017), while restricting the analysis to the genetically regulated component of expression (Musa et al., 2018; Duan et al., 2016; Zhang et al., 2019; Tian et al., 2025).

Together, TWAS and connectivity mapping provide a tractable computational pipeline from genetic association to candidate therapeutic hypothesis, and this general strategy has been applied across a range of complex human diseases (Peng & Zhang, 2025; Tian et al., 2025). However, these approaches have not yet been integrated within a unified framework for comparing the molecular consequences of a monogenic disorder with the polygenic architecture of a related common disease. In particular, existing work rarely evaluates the indirect molecular signature of a monogenic disorder obtained through pQTL data alongside the polygenic architecture of a related common disease (Tian et al., 2025). Few pipelines explicitly also address the linkage disequilibrium (LD) and study design issues that can compromise these integrated analyses.

In this study, we assembled eight publicly available GWAS-scale datasets relevant to ASD and FXS biology spanning four distinct study designs: (i) population GWAS comparing individuals with and without ASD; (ii) GWAS of ASD symptom severity based on differences within monozygotic twin pairs; (iii) case-case GWAS (CC-GWAS) comparing allele frequencies between ASD and each of three other psychiatric diagnoses; and (iv) a plasma pQTL study of FMRP itself. We use these data to (1) construct a properly powered ASD case-control meta-analysis that explicitly excludes case-case contrast statistics; (2) impute tissue-specific gene expression genome-wide across 49 GTEx v8 tissues for all eight traits using S-PrediXcan; (3) integrate association signals across tissues using the aggregated Cauchy association test (ACAT) and assess Hallmark pathway enrichment; (4) formally evaluate whether population ASD genetic risk and the monozygotic twin (MZ-twin) symptom discordance signal share common-variant architecture using an analysis explicitly corrected for LD-driven statistical inflation; and (5) query the CMap LINCS L1000 Level 5 dataset with a five metric connectivity scoring framework to nominate small molecules predicted to reverse the disease associated transcriptional signatures, cross referenced against mechanism of action (MOA) annotations from the Broad Institute Drug Repurposing Hub (Corsello et al., 2017). We report the resulting prioritized genes, tissues, pathways, and drug candidates and discuss both the biological implications of our findings and the methodological safeguards used throughout the analysis. Many of these safeguards address analytical issues that we identified and corrected during the study and consider important for rigorous integrative genomic drug repurposing analyses.

## Results

### Corrected ASD case-control meta-analysis

Meta-analysis of the population case-control ASD GWAS (iPSYCH-PGC 2017, *N* = 46,350; FinnGen) across 6,159,907 harmonized non-palindromic SNPs on the autosomes and chromosome X (harmonization and meta-analysis procedure described in Methods) yielded a genomic control inflation factor λ = 1.136. This level of inflation is compatible with a highly polygenic trait at this sample size, though genomic control inflation alone cannot distinguish polygenicity from residual confounding. Thirty SNPs reached genome-wide significance (*p* < 5 × 10⁻^8^), with most clustering within a single region on chromosome 20 (20:21.2-21.3 megabase [Mb], **Figure S1**; **Figure 1**). The chromosome 6 signal observed in an initial analysis arose from treating an ASD-versus-Tourette syndrome CC-GWAS as a population case-control GWAS. Reanalysis under the correct case-case interpretation showed that the signal reflects the ASD-versus-Tourette-syndrome contrast rather than population ASD case-control risk (**Figure S2**). To assess heterogeneity between the two contributing studies, we computed Cochran’s *Q* and the corresponding *I*² heterogeneity statistic (see Methods) for every SNP in the corrected meta-analysis. The vast majority of SNPs showed no detectable heterogeneity (*I*² = 0%), and only a small minority reached nominal significance on the heterogeneity test (*p* < 0.05; **Figure S3**). These results indicate no widespread evidence of heterogeneity between iPSYCH-PGC and FinnGen and support the use of a fixed effects framework for the primary meta-analysis. The full list of 30 genome-wide significant SNPs and the 67 independent lead loci are provided in **Tables S1** and **S2**, respectively.

**Figure 1.**
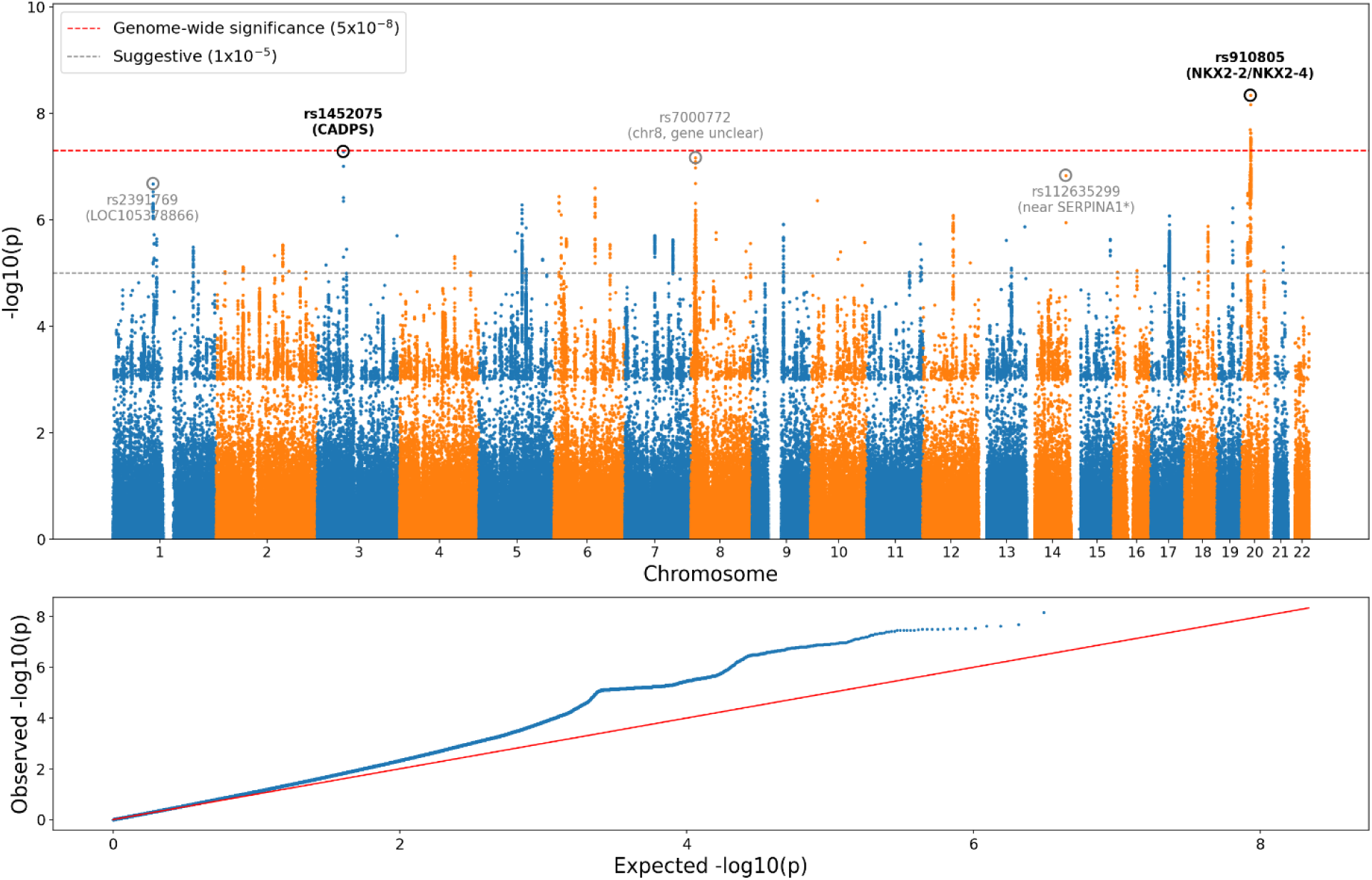
Manhattan and quantile-quantile plots for the corrected ASD case-control meta-analysis. The top panel shows the fixed-effects meta-analysis of the population case-control ASD GWAS datasets from iPSYCH-PGC 2017 and FinnGen across 6,159,907 SNPs. The bottom panel shows the corresponding quantile-quantile plot. The red dashed line indicates genome-wide significance (*p* = 5×10^-8^). *P*-values are unadjusted for genomic inflation. The genomic control inflation factor (λ = 1.136) is reported for diagnostic purposes only and was not applied as a correction to the test statistics or *p*-values.

### Genetic associations between population ASD risk and MZ-twin symptom discordance

Using genome-wide LD-pruned variant sets (pruning procedure described in Methods) containing 2,682 to 2,698 approximately independent variant sites per comparison, we found no evidence of *z*-score concordance between the meta-analyzed ASD case-control signal and either MZ-twin symptom-discordance GWAS. This was true for the all-age analysis (Pearson *r* = 0.008, *p* = 0.68, sign concordance 50.3%, binomial *p* = 0.77) and the childhood-restricted analysis (Pearson *r* = 0.003, *p* = 0.89, sign concordance 50.2%, binomial *p* = 0.86). Definitions of sign concordance and the binomial test are provided in the Methods. We next examined concordance among loci showing suggestive association with ASD (*p* < 10^-5^) in the meta-analysis. An initial analysis without pruning included 160 suggestive SNPs, of which 148 and 149 were also present in the two MZ-twin datasets, respectively, after allele harmonization. These SNPs showed strong apparent concordance, with Pearson *r* = 0.54 and 0.55 and sign concordance of 81% and 82% (both *p* < 10^-12^). However, examination of the underlying regions showed that this apparent signal was driven almost entirely by a single LD block: 113 of the 160 SNPs originated from a single region spanning approximately one Mb. The nominal significance was therefore substantially inflated by non-independence among the tested variants.

Restricting the analysis to independent lead variant sites (distance-based clumping procedure described in Methods) eliminated the apparent concordance. As a robustness check, we varied the physical pruning window from 500 kb to two Mb in 500 kb increments. This yielded 14 independent loci for the all-ages analysis and 15 for the restricted analysis at each of the four window sizes tested. Results were consistent across window sizes. For the all-ages comparison, Pearson correlation ranged from *r* = 0.08 to 0.25, with *p* > 0.05 for three of the four window sizes. For the childhood comparison, correlations ranged from *r* = 0.19 to 0.22, with *p* > 0.1 for all four window sizes. Sign concordance ranged from 51% to 57% in both comparisons, with all binomial *p* > 0.4. The consistency of these results across pruning thresholds indicates that the absence of detectable concordance is not driven by a particular choice of physical window. Together with the genome-wide analysis, these results provide no evidence for shared common-variant genetic architecture between population liability for ASD diagnosis and the genetic determinants of symptom severity discordance within twin pairs. The lack of concordance is consistent with the possibility that the twin discordance phenotype captures a genetically distinct component of ASD symptom variation, including effects specific to symptom discordance or involving interactions between gene-by-environment interactions. Genome-wide concordance after LD pruning is shown in **Figure S4**, and concordance restricted to independent lead variant sites is shown in **Figure S5**. At the genome-wide level, *z*-scores from the corrected ASD meta-analysis showed no detectable correlation with either MZ-twin symptom-discordance GWAS (MZtwin_ASD_all: *r* = 0.008, *p* = 0.68; MZtwin_ASD_children: *r* = 0.003, *p* = 0.89). Restricting the analysis to independent lead variant sites yielded weak concordance for MZtwin_ASD_all (*r* = 0.253, *p* = 0.046) but not MZtwin_ASD_children (*r* = 0.186, *p* = 0.15), with sign concordance near chance at (51% and 56 56%, respectively). This weak and inconsistent pattern contrasts with the much stronger apparent concordance observed when all nominally suggestive SNPs were analyzed without pruning, which was driven by correlated variant sites within a single LD block.

### Cross-tissue transcriptome-wide association identifies *XRN2*, *PPP3CA*, and *TNRC18*

Cross-tissue ACAT integration of S-PrediXcan results (see Methods) across 49 GTEx tissues identified two genes reaching Bonferroni-corrected significance for the corrected ASD case-control trait: *XRN2* (5’-3’ exoribonuclease 2, a transcription-termination factor; ACAT *p* = 2.3 × 10^-7^), supported across 40 tissues, and the uncharacterized long intergenic non-coding RNA *RP11-6N13.1* (ACAT *p* = 4.0 × 10^-7^). Neither gene reached significance in the twin-discordance or CC-GWAS trait analyses, consistent with these associations being specific to the population ASD liability signal (**Table 1**, **Figure 2**). For the FMRP plasma pQTL trait, *PPP3CA* (protein phosphatase 3 catalytic subunit alpha, the catalytic subunit of calcineurin) was the top cross-tissue integrated association (ACAT *p* = 1.4 × 10^-5^), with support across 33 of 49 tissues. *TNRC18* (trinucleotide-repeat-containing protein 18) was the second strongest association (ACAT *p* = 2.1 × 10^-5^), with its strongest single-tissue association in Brain Hypothalamus. Both genes were also among the top nominal gene associations in the individual tissue analyses conducted before integration, indicating that their signals were not produced solely by the ACAT combination procedure. The top 20 gene associations after integration across tissues for each trait are shown in **Table S3**. The numbers of genes tested and reaching Bonferroni significance for each trait are shown in **Figure S6**, and the inflation statistic (λ) calculated from the pooled S-PrediXcan *z*-scores for each trait is shown in **Figure S7**. The number of genes entering cross-tissue ACAT integration was broadly similar across traits (20,807–21,846), whereas the number reaching trait-specific Bonferroni significance differed sharply, with only ASD_caseControl_iPSYCH2017 yielding significant genes (*n* = 2). Given the similar testing burden across traits, this difference is primarily driven by differences in the underlying association signals rather than by differences in the number of genes tested. Inflation statistics were close to λ = 1 for most traits, with the three CC-GWAS-derived traits showing the highest values. This pattern indicates modest test-statistic inflation in these traits but does not establish its source.

**Figure 2.**
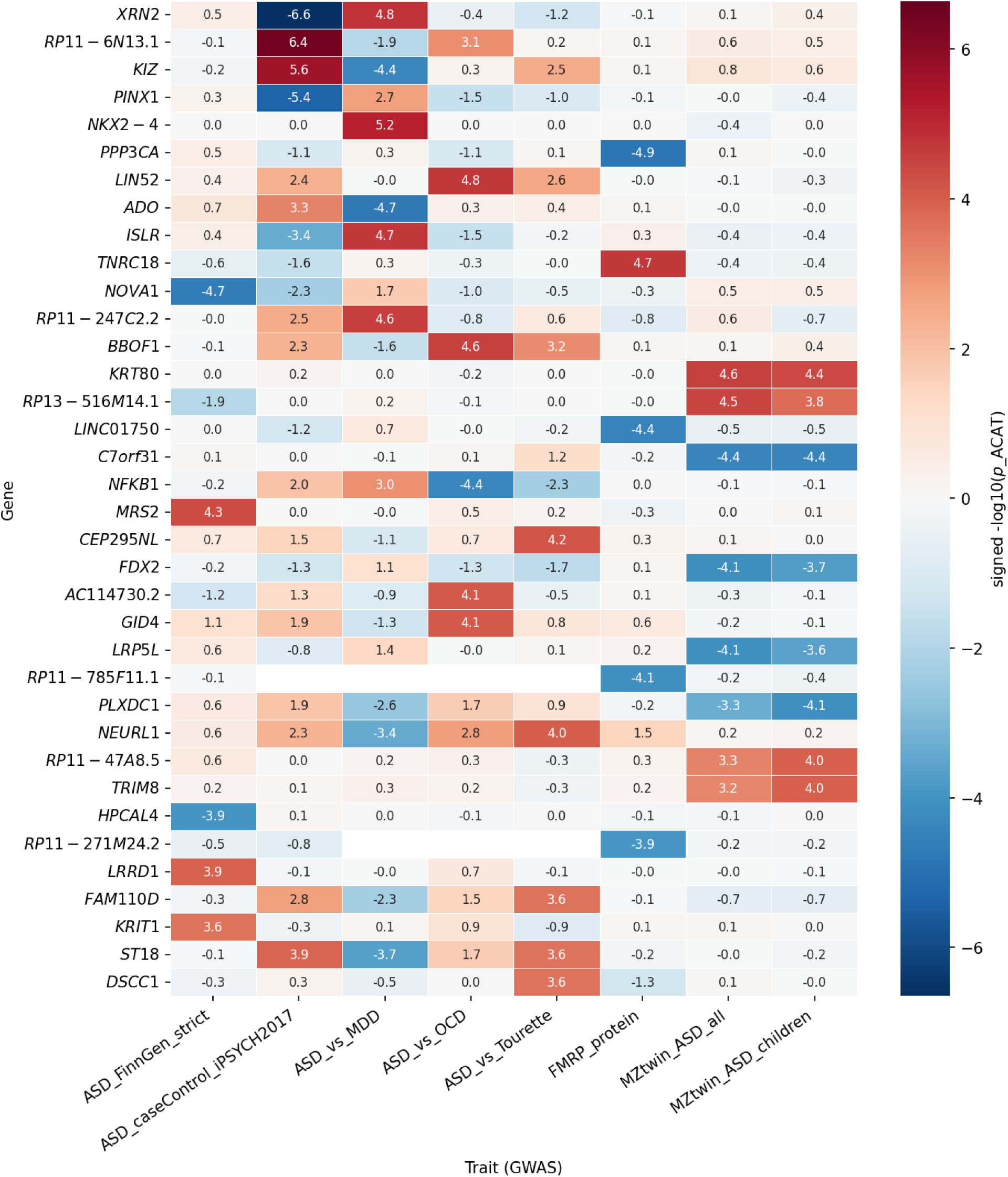
Heatmap of signed cross-tissue association significance for the top five integrated S-PrediXcan genes for each trait. The heatmap includes all eight traits. Each cell shows −log₁₀(*p*-value) from the cross-tissue ACAT analysis multiplied by the sign of the mean S-PrediXcan *z*-score across tissues. The magnitude indicates the strength of statistical significance, while the sign indicates the direction of association. Positive values indicate that higher genetically predicted expression is associated with higher trait values or disease risk, whereas negative values indicate the opposite. Values near zero indicate weak statistical association.

**Table 1.** Top two cross-tissue S-PrediXcan gene associations each trait.

| **Trait** | **Gene** | **ACAT $p$ value** | **Mean $z$** | **Tissues** | **Best tissue** |
| --- | --- | --- | --- | --- | --- |
| ASD_caseControl_iPSYCH2017 | *XRN2* | $2.26 \times 10^{-7}$ | -1.61 | 40 | Colon, Sigmoid |
| ASD_caseControl_iPSYCH2017 | *RP11-6N13.1* | $3.96 \times 10^{-7}$ | 5.07 | 1 | Testis |
| ASD_vs_MDD | *NKX2-4* | $6.80 \times 10^{-6}$ | 0.00 | 2 | Testis |
| FMRP_protein | *PPP3CA* | $1.36 \times 10^{-5}$ | -0.66 | 33 | Lung |
| ASD_vs_MDD | *XRN2* | $1.59 \times 10^{-5}$ | 1.21 | 37 | Skin (not sun exposed) |
| ASD_vs_OCD | *LIN52* | $1.70 \times 10^{-5}$ | 2.54 | 34 | Testis |
| FMRP_protein | *TNRC18* | $2.10 \times 10^{-5}$ | 2.25 | 11 | Brain Hypothalamus |
| ASD_FinnGen_strict | *NOVA1* | $2.22 \times 10^{-5}$ | -3.37 | 15 | Spleen |
| ASD_vs_OCD | *BBOF1* | $2.73 \times 10^{-5}$ | 2.70 | 45 | Colon, Transverse |
| MZtwin_ASF_all | *KRT80* | $2.82 \times 10^{-5}$ | 0.40 | 39 | Minor Salivary Gland |
| MZtwin_ASF_all | *RP13-516M14.1* | $3.16 \times 10^{-5}$ | 3.85 | 47 | Uterus |
| MZtwin_ASF_children | *KRT80* | $3.95 \times 10^{-5}$ | 0.36 | 39 | Minor Salivary Gland |
| MZtwin_ASF_children | *C7orf31* | $4.09 \times 10^{-5}$ | -2.78 | 42 | Prostate |
| ASD_FinnGen_strict | *MRS2* | $4.55 \times 10^{-5}$ | 3.17 | 46 | Adipose Visceral Omentum |
| ASD_vs_Tourette | *CEP295NL* | $6.68 \times 10^{-5}$ | 2.91 | 23 | Testis |
| ASD_vs_Tourette | *NEURL1* | $9.70 \times 10^{-5}$ | 0.60 | 24 | Spleen |

Calcineurin signaling mediated by *PPP3CA* intersects mechanistically with mGluR-dependent synaptic plasticity pathways implicated in FXS neurobiology (Contractor et al., 2015). This relationship provides a biologically plausible, hypothesis-generating connection between the FMRP pQTL signal and established FXS pathophysiology that is independent of any ASD case-control comparison. Gene-by-trait association patterns for the top integrated associations across all eight traits illustrate that most top genes are trait-specific rather than broadly shared (**Figure 2**). The notable exception is the substantial overlap between the two MZ-twin symptom-discordance traits, which share genetic samples and therefore should not be interpreted as independent replications. An extended heatmap showing the top 10 rather than top 5 genes for each trait is provided in **Figure S8**. This expanded view includes 73 unique genes, including the three highlighted genes (*XRN2*, *PPP3CA*, and *TNRC18*). Several genes show concordant signals specifically across MZtwin_ASD_all and MZtwin_ASD_children, consistent with the substantial sample overlap between these two related twin-discordance traits. Per-tissue S-PrediXcan *z*-scores for *XRN2*, *PPP3CA*, and *TNRC18* across all tested GTEx tissues are shown in **Figure S9**. *XRN2* and *PPP3CA* show both positive and negative *z*-scores across tissues, whereas *TNRC18* shows consistently positive *z*-scores.

### Pathway enrichment

Hallmark Gene set enrichment analysis (GSEA; see Methods) identified five trait pathway associations reaching false discovery rate (FDR) < 0.25 in at least one analysis. This threshold follows the convention established in the original GSEA methodology and is substantially more permissive than conventional genome-wide significance thresholds (Subramanian et al., 2005). The significant pathways were interferon alpha response for the contrast between ASD and Tourette syndrome; protein secretion and reactive oxygen species pathway in the FinnGen ASD case-control trait; oxidative phosphorylation for the childhood MZ-twin discordance trait; and cholesterol homeostasis in both MZ-twin discordance traits (**Figure 3**). A network linking traits, pathways, and leading-edge genes (see leading-edge gene definition given in Methods) is provided in **Figure 4**. Five of the eight traits connect to five distinct Hallmark pathways via seven trait-pathway edges. Oxidative phosphorylation shows opposite enrichment directions across its two connected traits (down in ASD_vs_MDD, up in MZtwin_ASD_children), illustrating trait-dependent rather than disease-general directionality, whereas cholesterol homeostasis is enriched in the same (negative) direction in both independent MZ-twin symptom-discordance traits, consistent with the cholesterol-pathway signal highlighted elsewhere in this study as being of particular biological interest.

**Figure 3.**
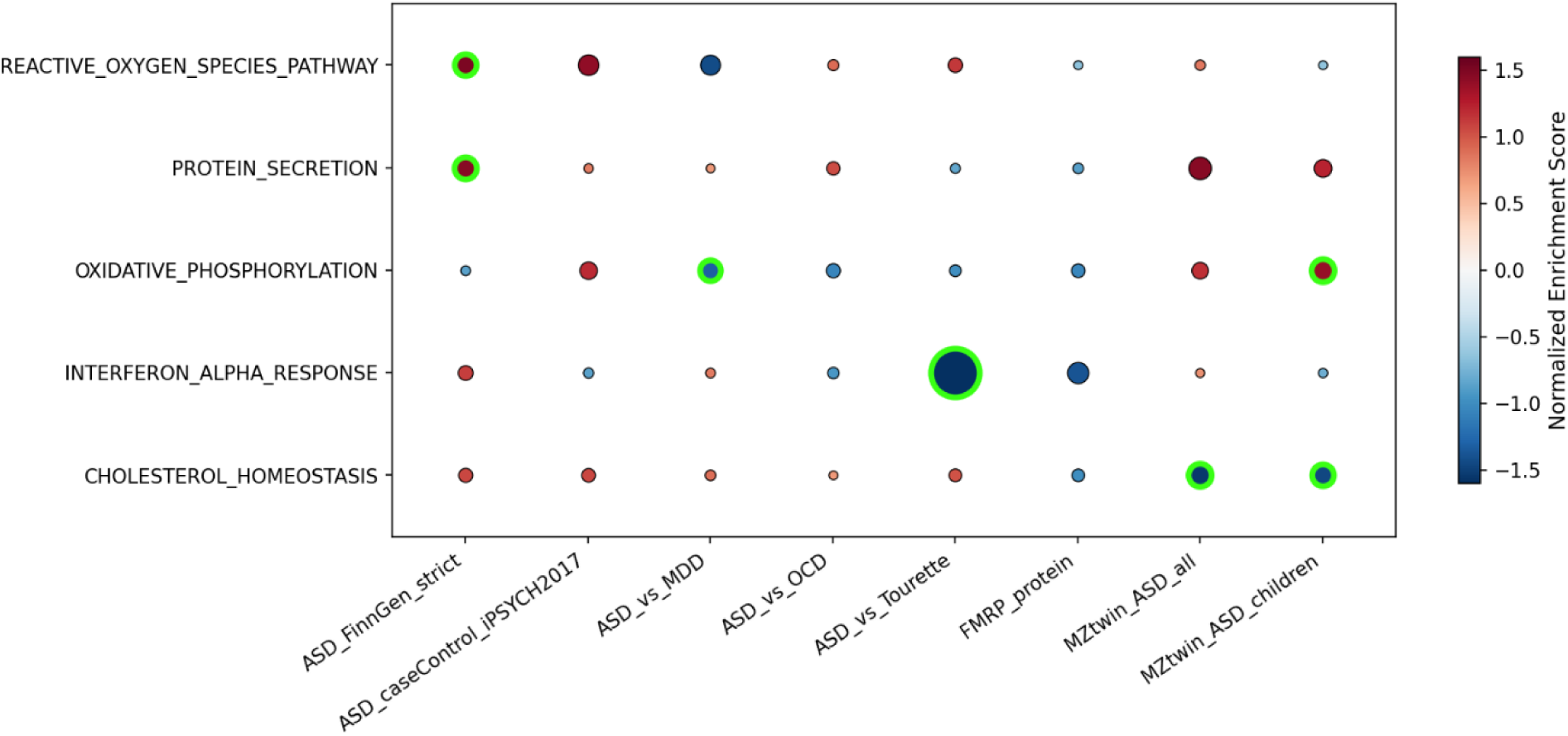
Hallmark GSEA normalized enrichment scores across traits. Pathways reaching FDR < 0.25 in at least one trait are shown across all eight traits. Dot size represents nominal significance, and green outlines indicate trait-pathway associations reaching FDR < 0.25.

**Figure 4.**
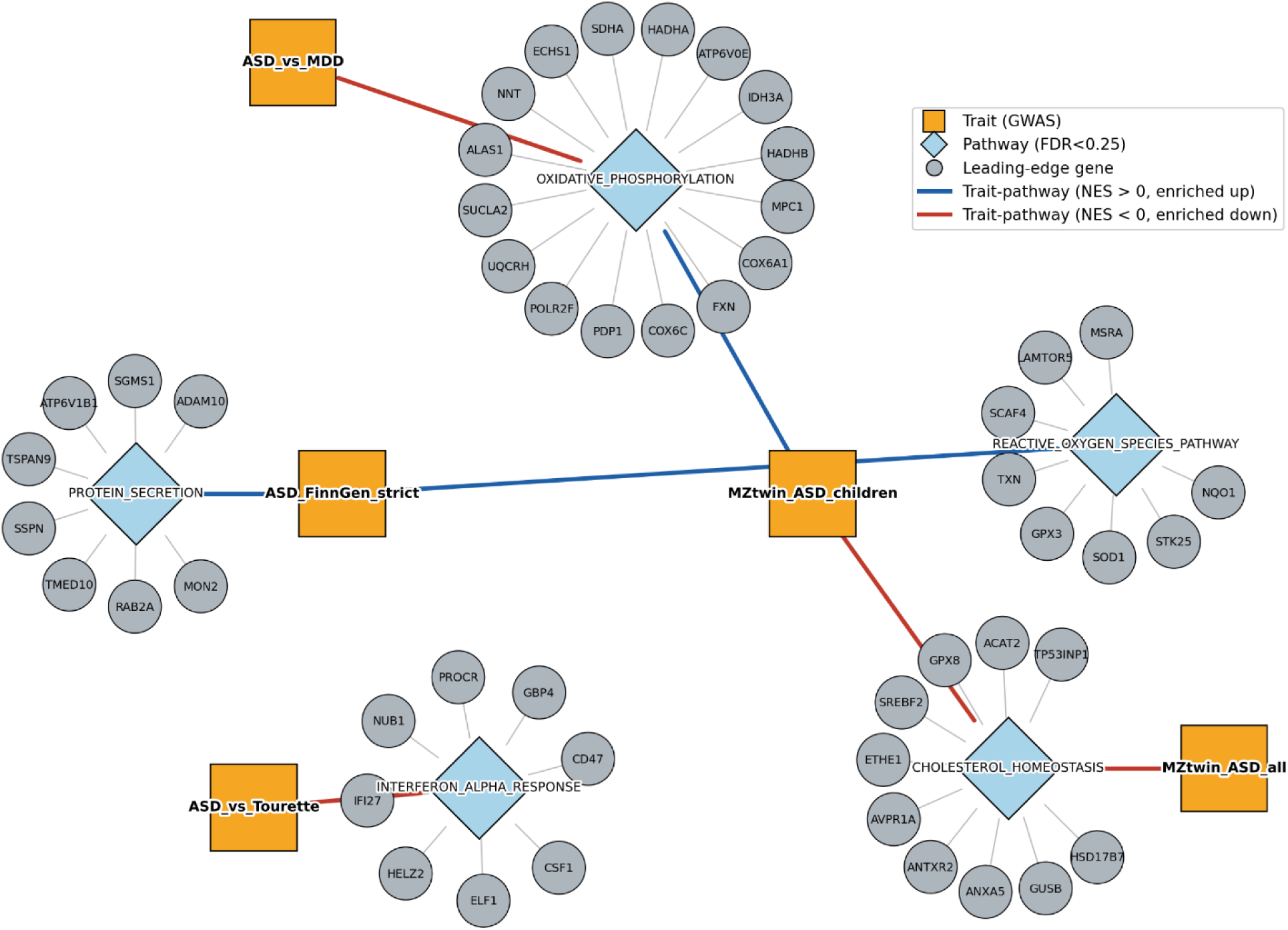
Trait, pathway, and gene network for significant Hallmark GSEA results. Significant pathway enrichments (FDR < 0.25) and the top eight leading-edge genes contributing to each pathway are shown. Five of the eight traits are represented, connecting to five distinct Hallmark pathways via seven trait-pathway edges, with edge color indicating the direction of enrichment (blue, NES > 0; red, NES < 0) and leading-edge genes shown as satellite nodes around each pathway. Oxidative phosphorylation is enriched in opposite directions across its two connected traits (down in ASD_vs_MDD, up in MZtwin_ASD_children), illustrating that the same Hallmark pathway can show trait-dependent directionality rather than a single, disease-general pattern. In contrast, cholesterol homeostasis is enriched in the same (negative) direction in both MZtwin_ASD_all and MZtwin_ASD_children, the two independent MZ-twin symptom-discordance traits, consistent with the concordant cholesterol-pathway signal highlighted as being of particular biological interest.

The concordant enrichment of cholesterol homeostasis in both the all-ages and childhood-restricted MZ-twin symptom-discordance analyses is of particular interest because disrupted cholesterol and lipid metabolism has been implicated repeatedly in ASD pathophysiology (Tierney et al., 2006; Lin et al., 2023). The MZ-twin design also reduces the potential influence of confounding factors shared by twins, including genetic background and aspects of rearing environment. This may provide some protection against confounding that can affect comparisons of unrelated cases and controls, though it does not eliminate all potential sources of bias (Myers et al., 2021). In particular, the childhood-restricted analysis is drawn from a subset of the same underlying cohort as the all-ages analysis. The concordant enrichment should therefore be regarded as internal consistency within a single study population rather than independent replication. Full Hallmark GSEA results for all 50 pathways across all eight traits are provided in **Table S4**. An unrestricted view of the top 25 pathways ranked by maximum absolute normalized enrichment score across all traits is shown in **Figure S10**, and classical GSEA running enrichment score plots for two representative trait-pathway pairs are shown in **Figure S11**. Most trait-pathway pairs in this extended view do not reach FDR < 0.25. The two representative running enrichment plots in **Figure S11** illustrate the corresponding enrichment patterns. For both cholesterol homeostasis (NES = −1.541, FDR = 9.96×10⁻³) and protein secretion (NES = 1.452, FDR = 1.55×10⁻²), pathway member genes are visibly concentrated toward one end of the ranked gene list rather than being uniformly distributed.

### Connectivity mapping nominates candidate reversal compounds

Across the eight trait signatures and 19,730 compounds, we evaluated 157,840 compound-trait combinations (see Methods). Of these, 2,788 met the prespecified candidate criteria of an empirical *z*-score < -2 relative to the tested compound library and concordant reversal direction across all five connectivity metrics. The number of candidates ranged from 265 (MZtwin_ASD_all) to 431 (ASD_vs_Tourette) per trait (**Figure 5**). Seventeen compounds were identified as significant reversal candidates in at least three of the eight traits, representing the compounds with the greatest cross-trait consistency. However, 14 of these 17 compounds lacked public compound names and therefore could not be readily interpreted or evaluated as therapeutic candidates without further chemical characterization. The empirical Kolmogorov-Smirnov (KS) *z*-score distribution for each trait is shown in **Figure S12**, the number of significant candidates per trait is shown in **Figure S13**, and the distribution of the number of traits in which each significant compound was identified is shown in **Figure S14**. Candidate counts per trait ranged from 265 (MZtwin_ASD_all) to 431 (ASD_vs_Tourette), summing to the 2,788 significant compound-trait pairs reported. Candidate counts did not scale simply with GWAS sample size. For example, ASD_caseControl_iPSYCH2017 (*N* = 46,350) yielded fewer candidates than ASD_vs_OCD (*N* = 21,069), indicating that the number of candidates depends on trait-specific signal strength rather than sample size alone. The large majority of significant compounds (2,202 of 2,486, 88.6%) were significant in only one trait, while 267 were significant in two traits, 16 in three traits, and one in four traits. The 17 compounds identified in three or more traits therefore represent the small right tail of the recurrence distribution. Full results for the 17 cross-trait candidates are provided in **Table S5**.

**Figure 5.**
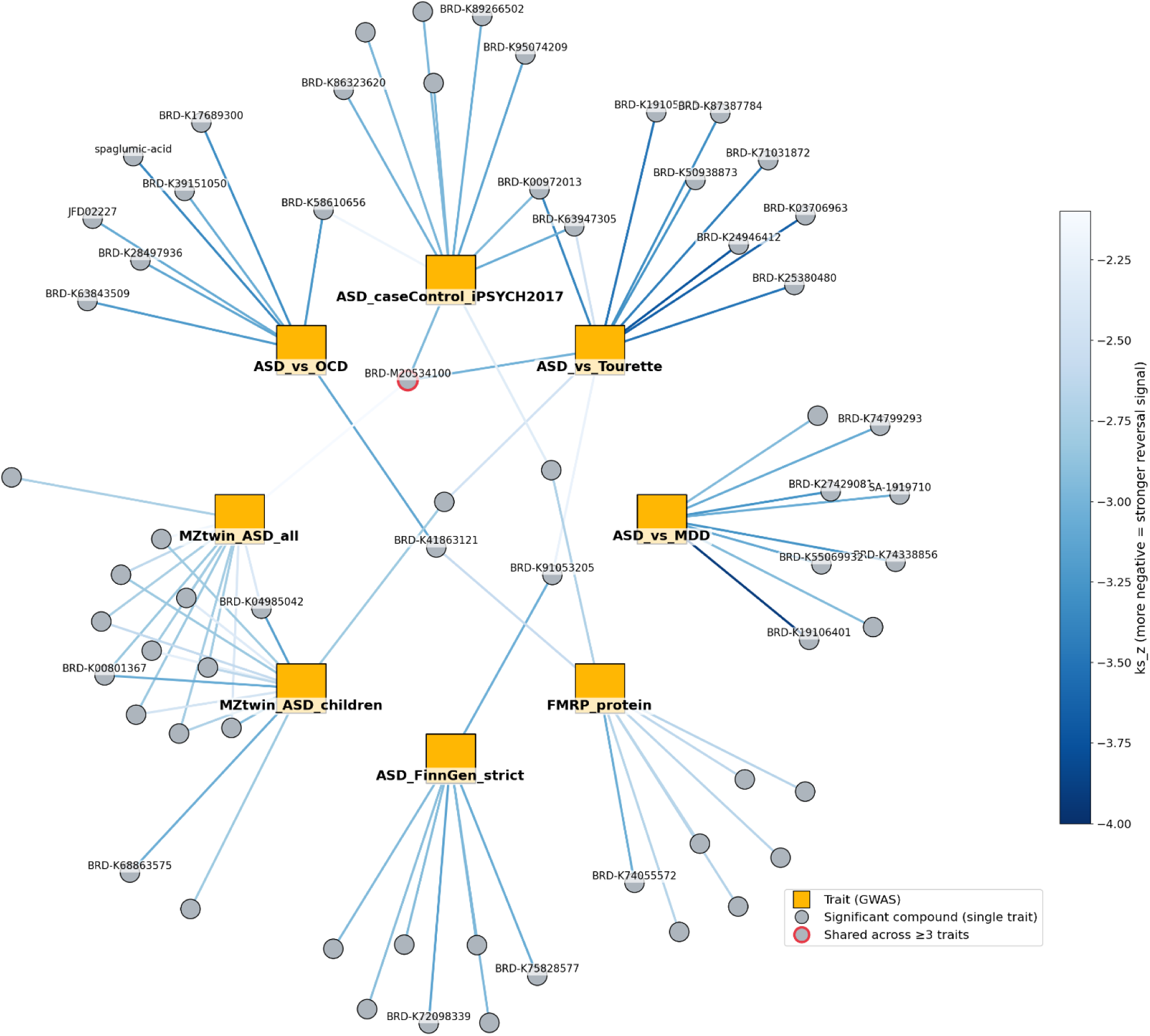
Trait and drug candidate network. For each trait, the top eight reversal candidates ranked by empirical *z*-score are shown as circular nodes connected to their corresponding trait (orange square). Compounds significant for more than one trait are shown as a single node with an edge to each associated trait. Node border color distinguishes compounds significant in a single trait (black) from those significant in three or more traits (red). One compound, BRD-M20534100, met the three-trait threshold, with edges to ASD_caseControl_iPSYCH2017, ASD_vs_Tourette, and MZtwin_ASD_all. Two additional compounds, BRD-K41863121 and BRD-K91053205, were each significant for exactly two traits and are therefore shown with two edges and a black border. Edge color represents the empirical z-score (ks_z) on a continuous blue scale, with the darker blue indicating a stronger reversal signal. The color scale spans the range of ks_z values shown in the network (−4.0 to −2.1). Compound names are shown only for compounds significant in multiple traits and for compounds with ks_z ≤ −3.0 to preserve legibility.

Among candidates with resolvable identities and MOA annotation (see compound identity resolution and MOA annotation source described in Methods), several biologically plausible compounds emerged (**Table 2**; **Table S6**). Spaglumic acid, annotated as a glutamate receptor antagonist, was the top reversal candidate for the contrast between ASD and obsessive compulsive disorder (empirical *z* = -3.35; **Table 2**). This finding is notable given the mechanistic importance of glutamatergic signaling in FXS pathophysiology (Contractor et al., 2015).

**Table 2.**
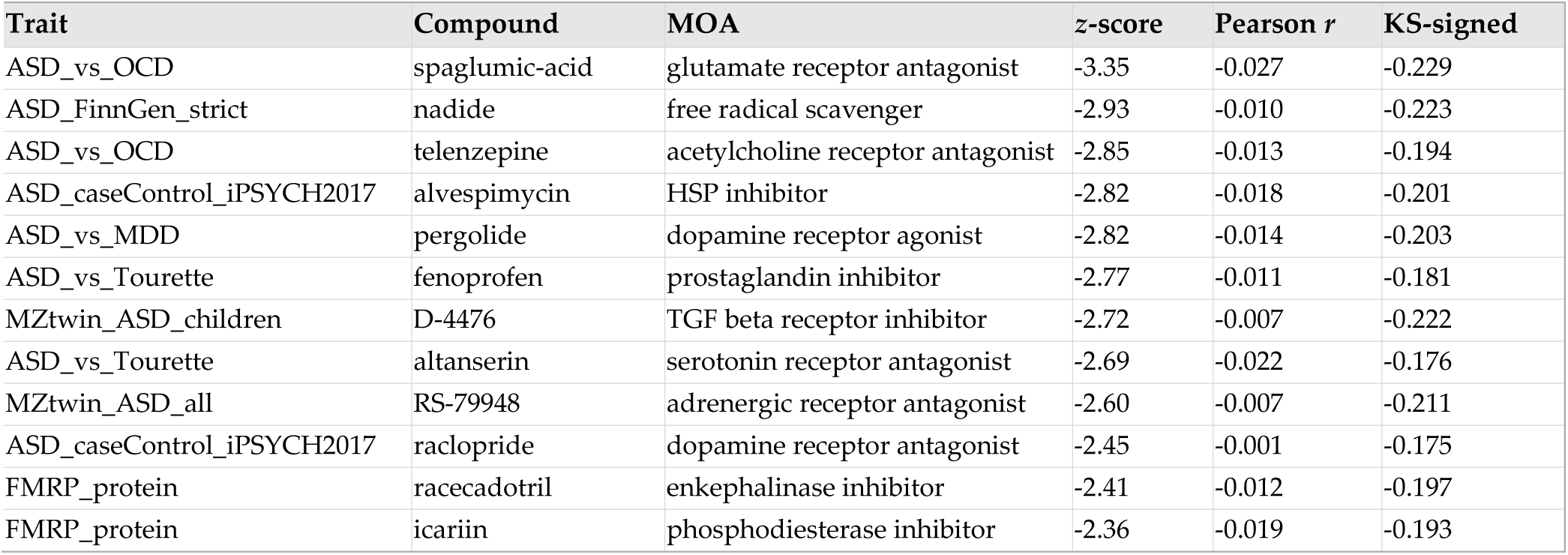
Top MOA-annotated reversal candidate for each trait.

| **Trait** | **Compound** | **MOA** | **z-score** | **Pearson *r*** | **KS-signed** |
| --- | --- | --- | --- | --- | --- |
| ASD_vs_OCD | spaglumic-acid | glutamate receptor antagonist | -3.35 | -0.027 | -0.229 |
| ASD_FinnGen_strict | nadide | free radical scavenger | -2.93 | -0.010 | -0.223 |
| ASD_vs_OCD | telenzepine | acetylcholine receptor antagonist | -2.85 | -0.013 | -0.194 |
| ASD_caseControl_iPSYCH2017 | alvespimycin | HSP inhibitor | -2.82 | -0.018 | -0.201 |
| ASD_vs_MDD | pergolide | dopamine receptor agonist | -2.82 | -0.014 | -0.203 |
| ASD_vs_Tourette | fenoprofen | prostaglandin inhibitor | -2.77 | -0.011 | -0.181 |
| MZtwin_ASF_children | D-4476 | TGF beta receptor inhibitor | -2.72 | -0.007 | -0.222 |
| ASD_vs_Tourette | altanserine | serotonin receptor antagonist | -2.69 | -0.022 | -0.176 |
| MZtwin_ASF_all | RS-79948 | adrenergic receptor antagonist | -2.60 | -0.007 | -0.211 |
| ASD_caseControl_iPSYCH2017 | raclopride | dopamine receptor antagonist | -2.45 | -0.001 | -0.175 |
| FMRP_protein | racecadotril | enkephalinase inhibitor | -2.41 | -0.012 | -0.197 |
| FMRP_protein | icariin | phosphodiesterase inhibitor | -2.36 | -0.019 | -0.193 |

Oleoylethanolamide was independently identified as a significant candidate in both MZ-twin symptom-discordance traits (**Table S6**). Valsartan, an angiotensin II receptor antagonist, was among the candidates shared across multiple traits (**Table S6**).

MOA enrichment analysis (Fisher’s exact test procedure described in Methods) identified T-type calcium channel blockade and angiotensin receptor antagonism as the two mechanistic classes with the strongest nominal enrichment among terms with a sufficiently large annotated compound set (*n* ≥ 32). Several terms ranked above these two by nominal *p*-value had identical *p*- values across multiple mechanistically unrelated MOA classes. These identical *p*-values arise because Fisher’s exact test yields only a limited number of distinct attainable *p*-values when the annotated compound group is very small (*n* = 8). The Fisher’s exact test *p*-values for T-type calcium channel blockade and angiotensin receptor antagonism were 0.0096 and 0.025, respectively (**Figure 6**; **Table S7**). These values are uncorrected nominal *p*-values rather than FDR-adjusted significance estimates. Unlike the Hallmark GSEA analysis, for which FDR < 0.25 follows an established convention for this type of gene-set analysis (Subramanian et al., 2005), no comparable threshold was prespecified for the MOA enrichment analysis. Moreover, Benjamini-Hochberg correction across all 473 tested MOA terms resulted in no significant term, even at *q* < 0.25. These results reflect the limited statistical power resulting from MOA annotation being available for only 1,773 of the 19,730 tested compounds. Thus, the observed enrichments for T-type calcium channel blockade and angiotensin receptor antagonism should be regarded as exploratory, hypothesis-generating observations rather than statistically confirmed enrichments. Glutamate receptor antagonism did not show class-level enrichment (nominal *p* = 0.89). Reference to spaglumic acid (**Table 2**) reflects a single-compound finding rather than enrichment of its annotated mechanistic class. Full results for all 473 MOA terms are provided in **Table S7**, and the top 30 terms are shown in **Figure S15**. The five connectivity metrics used to call reversal candidates were not fully independent. Pairwise correlation among them (**Figure S16**) revealed two distinct clusters, with the two full-signature correlation metrics (Pearson, Spearman) forming one cluster and the KS-signed statistic and the two extreme-gene correlation metrics forming the other. The second cluster is more tightly correlated, while correlation between the two clusters was comparatively weaker. This structure indicates that requiring concordance across all five metrics provides only modest additional stringency beyond the ks_z < −2 threshold alone. A subset of the top candidates was additionally evaluated against the published FXS and ASD pharmacology literature (evaluation criteria described in Methods), with the results of this manual validation provided in **Table S6** and **Figure S17**.

We linked the top significant MOA-annotated compound for each trait to the genes contributing most strongly to its connectivity signal and to the GTEx tissues in which those genes showed their strongest associations (gene-contribution and best-tissue assignment procedures are described in Methods) (**Figure 7**). Several of the strongest reversal signals were associated with genes whose best-supporting tissues include amygdala, hippocampus, cerebellum, and basal ganglia structures. These tissue-level associations provide additional context for a subset of candidate compounds. However, they should not be interpreted as evidence that the corresponding compounds act specifically through neural issues because the compound connectivity signatures were predominantly derived from non-neural cell lines.

**Figure 6.**
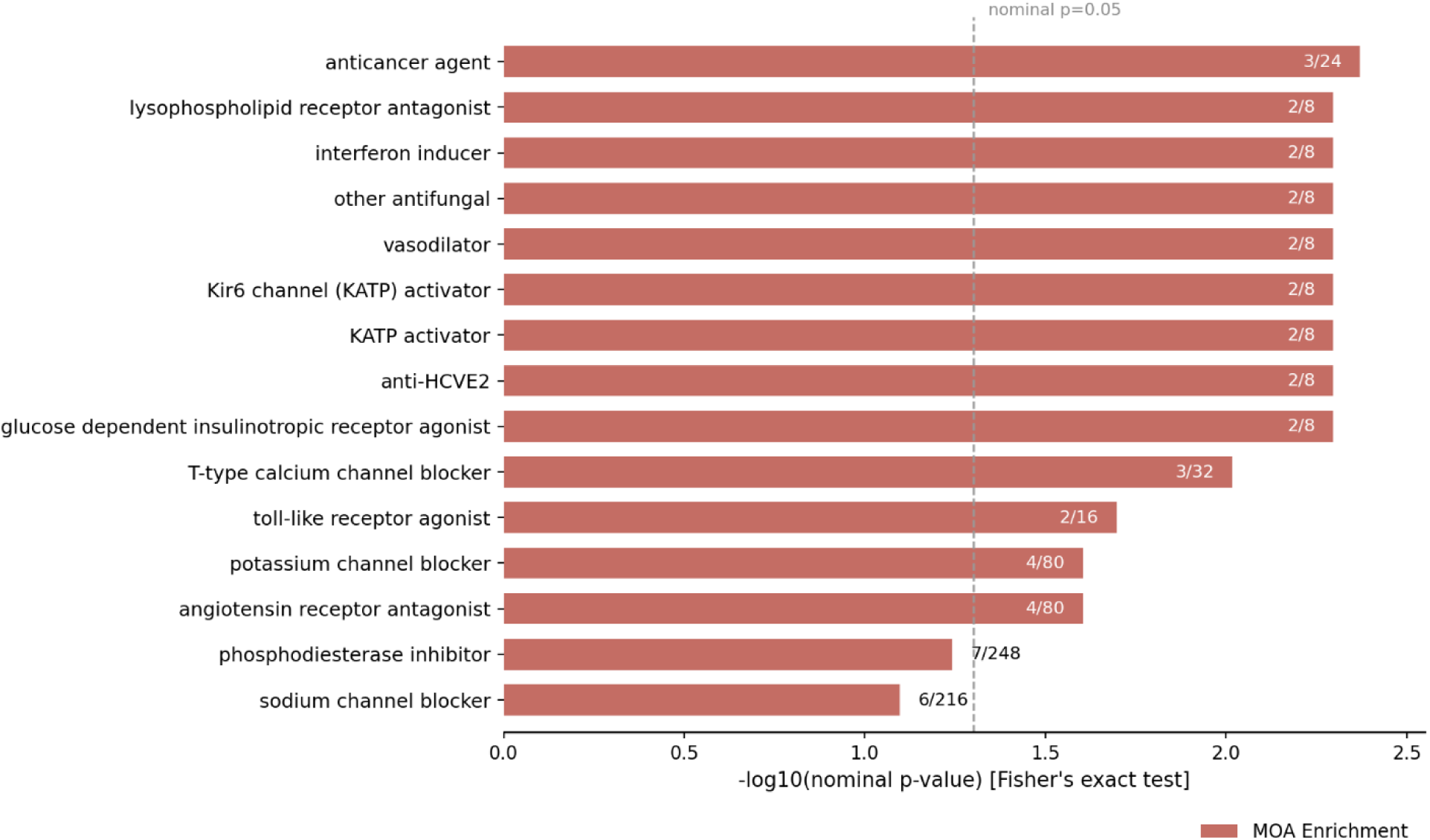
MOA enrichment among significant reversal candidates. The 15 elementary MOA terms with the smallest nominal Fisher’s exact test *p*-values are shown.

**Figure 7.**
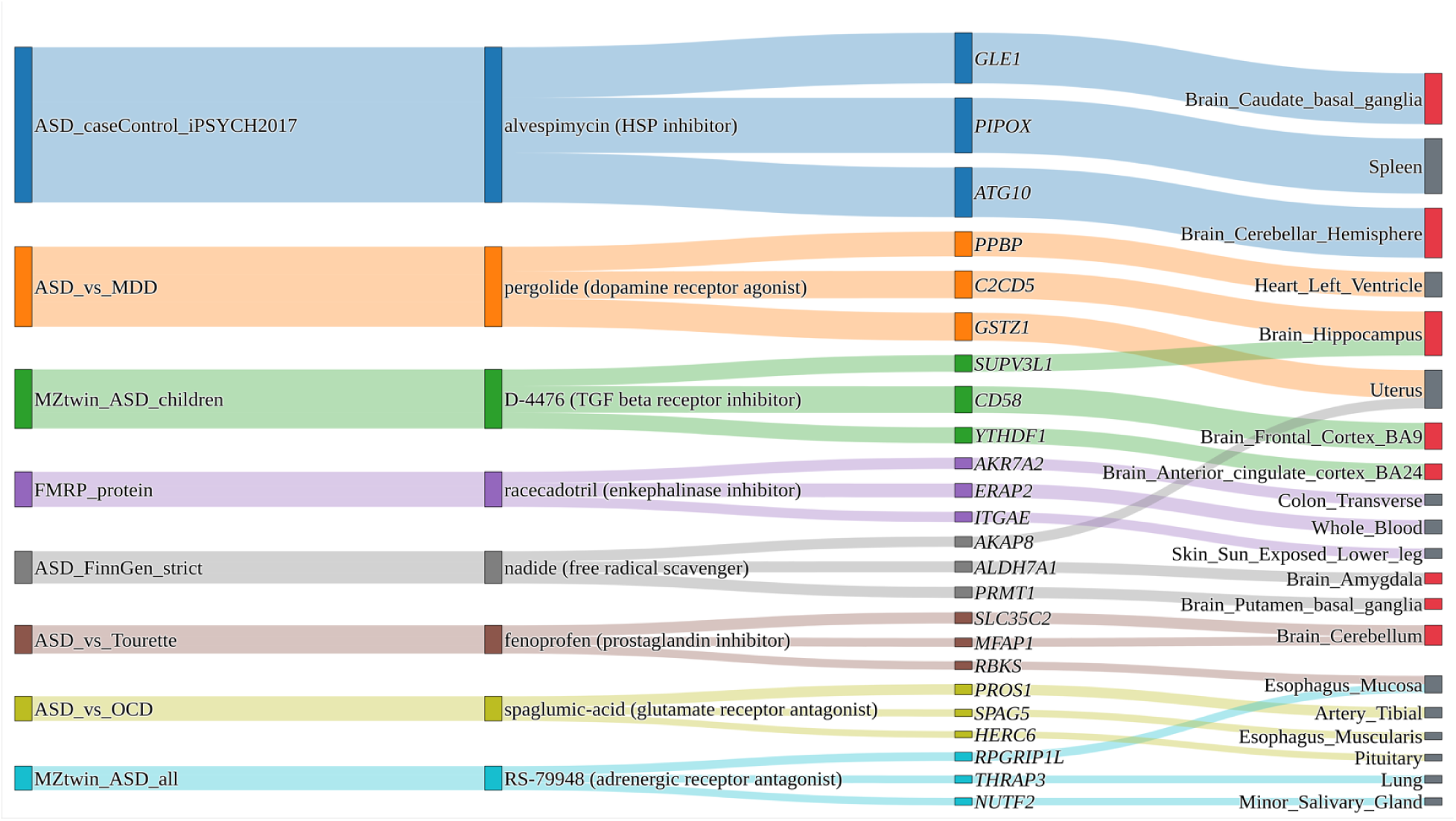
Sankey diagram linking traits, compounds, genes, and tissues. For the top MOA-annotated reversal candidates for each trait, the corresponding trait, compound and MOA annotation, top three contributing genes, and best-supporting GTEx tissue for each gene are shown.

Non-neural cell lines dominate the LINCS L1000 exemplar signature library, whereas neural lineages are substantially underrepresented. Prostate (VCAP), lung (A549), and breast (MCF7) cancer cell lines accounted for 34.1% of the library, comprising 38,320 of the 112,495 qualifying *trt_cp* exemplar signatures. Here, *trt_cp* denotes small-molecule compound treatment signatures, as opposed to signatures generated by genetic perturbations such as shRNA knockdown or cDNA overexpression. In contrast, neural progenitor cells (NPC; 3,496 signatures) and neuronal cells (NEU; 2,538 signatures) together accounted for only 5.4%. Across the 19,730 tested compounds, 3,374 (17.1%) had at least one qualifying *trt_cp* exemplar signature in either the NPC or NEU context. This imbalance limits the extent to which the connectivity results can be interpreted as neural-specific drug responses. The cell-line composition of the CMap L1000 exemplar signature library is shown in **Figure S18**. Neural-lineage cell lines were markedly under-represented relative to the most heavily profiled lines. NPC (3,496 signatures) and NEU (2,538 signatures) respectively ranked ninth and eleventh among the top 15 cell lines, together contributing fewer signatures than VCAP alone (15,390 signatures), the most represented cell line. The distribution of the number of GTEx tissues showing significant S-PrediXcan associations among the top contributing genes for the top compound-trait pairs is shown in **Figure S19**. Most candidate compound-trait pairs showed broad cross-tissue signal, with 26 of 40 pairs (65%) showing significant associations in at least 45 of the 49 tissues. This pattern is consistent with substantial sharing of genetically regulated expression signals across tissues among many of the contributing genes. **Figures S20** shows the same links restricted to cases in which the best- supporting tissue was a brain region. All eight traits retained at least one linked to a brain region, spanning both cortical and subcortical or cerebellar structures.

To further characterize the mechanistic structure of the candidate pool, we performed hierarchical clustering using Ward linkage (see Methods) of the top 30 MOA-annotated reversal candidates. Clustering was based on the Spearman correlation of the connectivity KS *z*-score profile for each compound across the eight traits was cross-referenced against Broad Repurposing Hub MOA annotations (**Figure 8**). Candidates with shared or related mechanisms, including serotonergic, dopaminergic, and adenosine or purinergic compounds, tended to cluster together despite being nominated independently across different traits. Several clusters also contained compounds acting on the same receptor system in opposite pharmacological directions, such as agonists and antagonists. This pattern is consistent with the possibility that modulation of a shared pathway in opposite directions can produce trait-specific reversal signatures. The clustering structure therefore provides a complementary descriptive view of the mechanistic organization of the candidate pool, though it should not be interpreted as independent statistical validation of the drug repurposing results.

**Figure 8.**
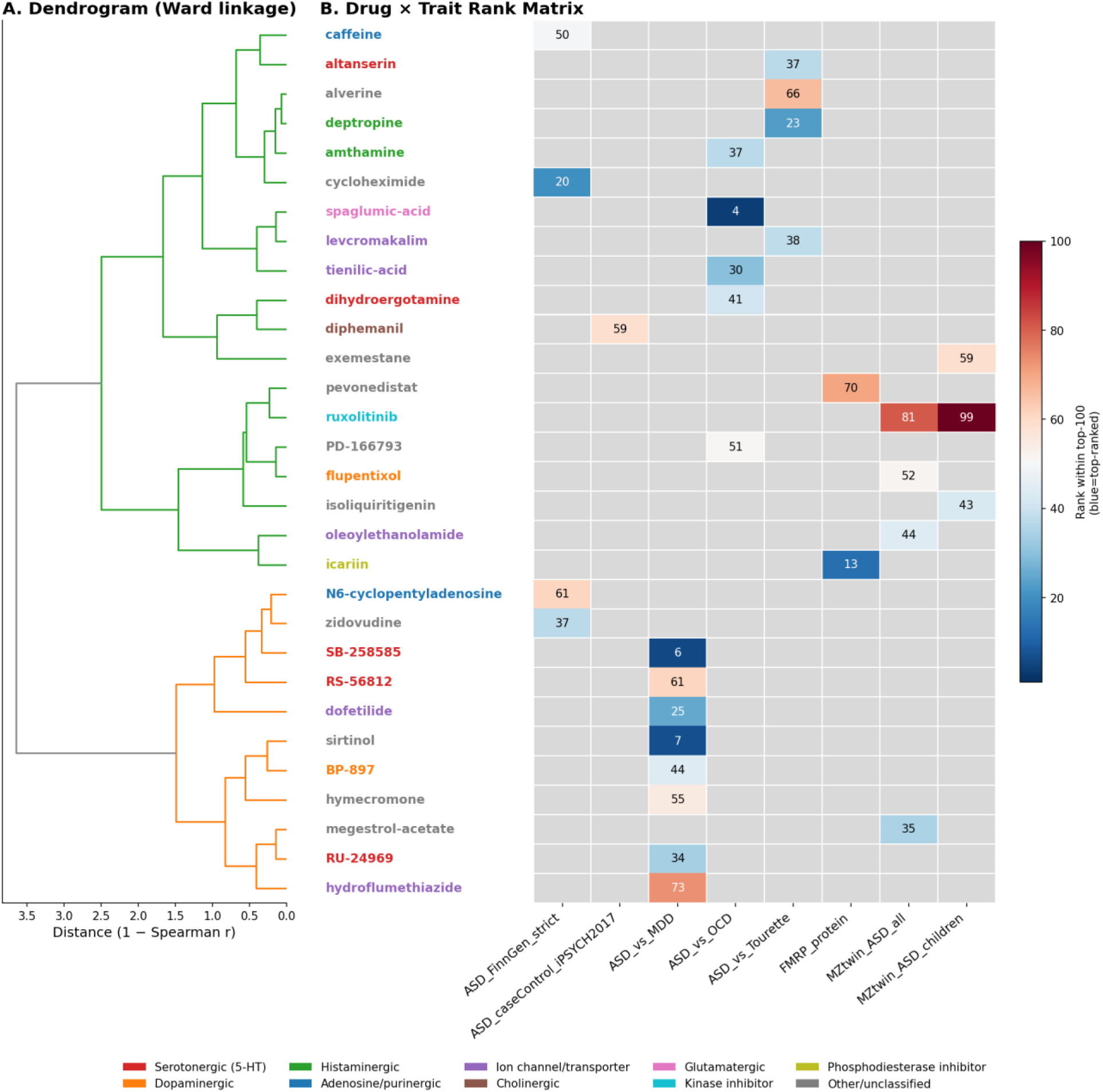
Functional similarity clustering of top computational drug-repurposing candidates. (A) Hierarchical clustering using Ward linkage of the top 30 MOA-annotated candidate compounds based on Spearman correlation of their connectivity (KS *z*-score) profiles across all eight traits. Drug-name colors indicate MOA classes based on Broad Repurposing Hub annotation. (B) Corresponding compound-by-trait rank matrix. Each cell shows the rank of the compound among the top 100 reversal candidates for that trait, with lower ranks indicating stronger reversal signals. Gray cells indicate that the compound was not among the top 100 reversal candidates for that trait.

## Discussion

By integrating population case-control, within twin discordance, case-case contrast, and plasma protein pQTL data with tissue-resolved transcriptome imputation and small molecule connectivity mapping, we identified candidate genes and compounds that provide several testable hypotheses concerning the relationships between FXS biology and ASD. *XRN2* was the strongest cross-tissue gene association for population ASD risk, while *PPP3CA* was the top cross-tissue association for the FMRP plasma pQTL. We also identified candidate compounds whose transcriptional signatures were predicted to reverse the genetically associated expression signatures, including compounds with biologically coherent MOA. At the same time, several of the strongest findings require cautious interpretation because cross-tissue consistency does not establish causal involvement. The connectivity analysis was based largely on non-neural cell lines, and the genetic overlap analysis used an LD pruned proxy rather than formal genetic correlation analysis.

The absence of detectable shared common-variant architecture between population ASD risk and MZ-twin symptom discordance is itself informative. After accounting for the substantial inflation produced by correlated variant sites within associated regions, we found no evidence that the population ASD signal and the signal associated with symptom discordance within MZ-twin pairs reflect the same common-variant architecture. This finding is consistent with the possibility that factors associated with whether an individual develops ASD and factors associated with differences in symptom severity between genetically identical individuals are at least partly distinct. The latter could include effects specific to symptom discordance or interactions between genes and the environment. However, the present analysis does not establish the biological mechanism underlying the MZ-twin associations, and the absence of detectable genetic correlation should not be interpreted as evidence that symptom severity has no genetic contribution.

This distinction has implications for the design of future genetic studies of ASD symptom severity. Case-control designs are well suited to identifying genetic factors associated with ASD diagnosis but are less directly suited to identifying factors associated with variation in symptom severity among affected individuals. Within family designs, including MZ-twin discordance studies, provide a complementary strategy because they control for genetic background and other actors shared within twin pairs. The corrected chromosome 20q13.13 locus identified in our meta-analysis, together with the *XRN2* transcriptome-wide association, should therefore be regarded as leads for follow-up rather than as mechanistically established findings. We did not perform fine mapping or functional annotation beyond the gene level TWAS association, and the causal gene or regulatory element underlying the chromosome 20 locus remains to be established.

The cross-tissue results for *PPP3CA* and *TNRC18* provide additional evidence that their associations are not confined to a single tissue prediction model, but this consistency should not be interpreted as tissue level replication. *PPP3CA* and *TNRC18* were supported across 33 and 11 GTEx tissues, respectively, and both were among the strongest nominal associations in individual tissue analyses. However, cross-tissue TWAS associations can arise from shared eQTL architecture and LD structure across prediction models trained on overlapping reference genotypes. Thus, observing an association across multiple tissues does not by itself establish a tissue-specific causal mechanism. As an exploratory analysis, we examined whether the single lead SNP comprising the MASHR expression prediction model for each of *XNR2*, *PPP3CA*, and *TNRC128* also occurred within a broader regional association signal in the corresponding GWAS. *XRN2* showed strong regional support, with 354 of 1,036 SNPs (34.2%) within 250 kb of the lead SNP reaching *p* < 10^-5^, consistent with a broader association signal at the same chromosome 20 locus identified in the ASD case-control meta-analysis. *PPP3CA* showed more modest regional support, with 10 of 1,671 SNPs (0.6%) reaching *p* < 10^-5^, though the strongest regional SNP reached *p* = 3.5×10^-8^). *TNRC18* showed the weakest regional support, with two of 2,705 SNPs (0.07%) reaching *p* < 10^-5^, with the signal concentrated in one or two highly correlated SNPs. This gradient in regional support suggests that the three genes differ in the strength of evidence connecting their expression prediction signals to broader GWAS associations patterns. These results are exploratory and do not establish colocalization, but they provide a basis for prioritizing candidates for formal fine mapping and functional follow-up.

*PPP3CA* encodes the catalytic subunit of calcineurin, a calcium and calmodulin dependent serine/threonine phosphatase that has an established role in activity dependent synaptic plasticity and is a downstream effector of mGluR signaling, a pathway implicated in FXS molecular pathophysiology (Contractor et al., 2015; Darnell et al., 2011). That identification of *PPP3CA* as the top cross-tissue association for a plasma pQTL of FMRP therefore provides a genetics first, hypothesis generating connection between circulating FMRP regulation and calcineurin mediated synaptic plasticity. This association is particularly useful because it does not depend on an ASD diagnosis phenotype and therefore provides an independent route for identifying biological processes related to FMRP regulation. Calcineurin pathway modulation in FXS model systems may therefore warrant further investigation, though the present results do not establish that *PPP3CA* itself mediates pQTL association or that calcineurin modulation would have therapeutic benefit (Bagni & Zukin, 2019).

*XRN2* emerged as the strongest cross-tissue gene association for population ASD risk. *XNR2* is a broadly expressed transcription termination and RNA surveillance factor with no established direct link to ASD neurobiology (West et al., 2004; Miki et al., 2013). Its role in RNA processing is nevertheless compatible with broader evidence implicating post-transcriptional regulation in ASD genetics (Darnell et al., 2011). Given the absence of fine mapping, formal colocalization, or functional validation, we regard the *XNR2* association as a lead for independent replication and mechanistic investigation rather than as a resolved biological mechanism.

The correction of the ASD GWAS analysis also illustrates an important methodological issue. CC-GWAS contrast statistics estimate a fundamentally different quantity from population case-control GWAS. They identify loci with divergent allele frequencies between the case populations of two disorders rather than differences between affected individuals and unaffected controls. Treating such a contrast as a population case-control GWAS therefore does more than reduce statistical power. It changes the estimand and undermines the construct validity of the resulting meta-analysis. In our analysis, the incorrectly included contrast produced a different signal from the corrected population case-control analysis, consistent with its representing the ASD versus Tourette syndrome contrast rather than population ASD risk. Dataset accession numbers and superficial trait name matching are therefore insufficient for establishing study design equivalence. GWAS pipelines should verify the underlying study design against the original source publications and confirm that all input datasets estimate comparable quantities before pooling their summary statistics.

A second methodological issue concerns the genetic overlap analyses in settings where formal LD score regression (LDSC; Bulik-Sullivan et al., 2015) is unavailable or unsuitable. Correlation of *z*-scores at nominally associated loci can produce dramatically inflated apparent concordance when multiple variant sites from the same LD block are treated as independent observations. In our analysis, an initially strong concordance signal among ASD suggestive loci was driven almost entirely by a single LD block. Restricting the analysis to independent lead variant sites reduced this apparent concordance to a null result. This does not make LD-pruned analysis equivalent to LDSC, and the resulting estimates should therefore be regarded as provisional. Nevertheless, genetic overlap analysis that cannot employ LDSC or an equivalent formal method should at minimum report results from LD-pruned variant set or a set of independent lead variant sites alongside any unpruned analysis. This allows the extent to which apparent concordance depends on correlated variant sites within individual loci to be assessed directly.

Several limitations of the connectivity analysis also warrant emphasis. First, CMap signatures were aggregated to one representative profile per compound based on transcriptional activity score without regard to the cell line from which the signature originated. Because the LINCS L1000 reference panel is dominated by cell lines with limited direct relevance to central nervous system biology, the selected representative signature will often reflect a peripheral or cancer cell context rather than a neural context. This may dilute or obscure reversal signals that are specific to neuronal biology. A cell line stratified reanalysis restricted to neural progenitor and neuron derived signatures would directly address this limitation. This present dataset provides a feasible basis for such analysis because 3,374 compounds had at least one qualifying neural lineage exemplar signature.

Second, our empirical connectivity significance framework prioritizes compounds according to their relative position within the tested compound library and does not provide a formal multiple testing correction for all compound trait combinations. The threshold used here was therefore intentionally exploratory. In addition, the connectivity metrics are correlated with one another and do not account for every compound specific technical feature of the underlying signatures. The requirement for concordance across all five metrics provides an additional filtering criterion, but it does not transform the resulting candidates into statistically confirmed therapeutic effects.

Third, MOA annotations were available for only 9.0% of the tested compound library. This limited annotation coverage substantially reduces the power of MOA enrichment analysis and likely contributes to the absence of FDR significant mechanistic classes despite several biologically coherent nominal associations. The glutamate receptor antagonist, angiotensin receptor antagonist, and T type calcium channel blockade categories should therefore be interpreted as exploratory observations rather than statistically established enrichments.

Fourth, our LD aware genetic overlap analysis is a pragmatic alternative to formal LDSC rather than a replacement for it. The null result between population ASD risk and MZ-twin symptom discordance should be interpreted as an absence of detectable concordance under the specific LD pruning framework used here. A formal genome-wide genetic correlation analysis could provide a more direct estimate of genetic covariance if appropriate summary statistics and reference data become available.

Despite these limitations, several compound candidates warrant further experimental evaluation. Glutamate receptor antagonism was nominally enriched among the reversal candidates and was specifically represented by spaglumic acid, which was the top candidate for the ASD versus OCD contrast. This finding is notable given the central role of glutamatergic signaling in FXS pathophysiology and provides an independent, transcriptome based indication that modulation of glutamatergic signaling may warrant continued investigation despite the disappointing clinical results of mGluR5 directed approaches (Contractor et al., 2015). The result does not imply that glutamate receptor antagonism as a class is therapeutically effective in FXS or ASD, but it identifies alternative pharmacological approaches within this pathway for experimental testing.

Oleoylethanolamide (cannabinoid receptor agonist and PPAR receptor agonist, among other annotated mechanisms; **Table S6**) was also identified in both MZ-twin symptom discordance analyses. The two analyses are not independent because the childhood-restricted dataset is derived from a subset of the broader twin cohort, so this observation should be interpreted as internal consistency rather than independent replication. Nevertheless, the association is of interest given evidence implicating endocannabinoid signaling in ASD (Aran et al., 2019; Chakrabarti et al., 2015; Siniscalco, et al. 2017). As with the other compounds identified here, oleoylethanolamide should be regarded as a computational candidate rather than a therapeutic lead established by the present study.

Several extensions follow naturally from these findings. A cell line stratified analysis of the connectivity mapping results, restricted to neural progenitor and neuron derived LINCS L1000 signatures, would determine whether the candidates identified here show stronger or more consistent reversal in a cellular context more directly relevant to ASD and FXS neurobiology. A formal bivariate LDSC analysis (Bulik-Sullivan et al., 2015) would provide a more rigorous assessment of genetic overlap between population ASD risk and MZ-twin symptom discordance and could potentially be extended to partitioned heritability analyses across functional genomic annotations (Finucane et al., 2015). Finally, experimental testing of the prioritized compounds in relevant cellular and animal models is necessary before any therapeutic inference can be made. In particular, cell based validation in FXS relevant neuronal models could determine whether the glutamatergic and cannabinoid/PPAR related candidates identified here produce the predicted molecular and phenotypic effects.

Taken together, these analyses demonstrate both the potential and the limitations of integrating genetically anchored expression signatures with pharmacological connectivity data to generate hypotheses about relationships between monogenic and polygenic disease biology. *XRN2* provides a candidate gene for population ASD risk that warrants independent replication and functional investigation, while *PP3CA* provides a biologically plausible connection between FMRP regulation and calcineurin mediated synaptic plasticity that can be investigated independently of ASD case status. The connectivity analysis similarly identifies compounds and mechanisms that merit experimental testing, but these candidates remain computational hypotheses until they are validated in biologically relevant models. More broadly, the study illustrates two safeguards that are important for integrative genomic drug repurposing analyses: verifying that GWAS datasets estimate comparable quantities before meta-analysis and accounting for LD before interpreting apparent genetic overlap. These considerations are particularly important when integrating datasets generated under different study designs, where superficially similar trait labels can conceal fundamentally different statistical estimands.

## Methods

An overview of the complete analytical pipeline is provided in **Figure S21**. Pipeline implementation details can be found in the GitHub repository https://github.com/zwerhu/ASD_FXS.

### GWAS summary statistics acquisition and quality control

Eight GWAS-scale summary statistics datasets were obtained in GWAS-VCF format from the IEU OpenGWAS project and the NHGRI-EBI GWAS Catalog (**Table S8**). These datasets comprised: (1) a population case-control ASD GWAS (18,381 cases, 27,969 controls; iPSYCH-PGC 2017 data freeze; Grove et al., 2019); (2) a FinnGen register-based case-control ASD phenotype using a stricter case-exclusion definition ("KRA_PSY_AUTISM_EXMORE"); (3, 4) two within MZ-twin pair difference GWAS of ASD symptom scores, one spanning all ages (*N* = 12,354 pairs) and one restricted to childhood assessments (*N* = 10,494 pairs), from the Within Families Consortium; (5) a plasma pQTL GWAS of FMRP (*N* = 3,301, INTERVAL study; Sun et al., 2018); and (6-8) three CC-GWAS case-case contrast statistics comparing ASD with Tourette syndrome, obsessive-compulsive disorder, and major depressive disorder (MDD). These contrasts identify loci with divergent allele frequencies between the case populations of two disorders rather than between cases and population controls (Peyrot & Robinson, 2021). Chromosome X coverage for each dataset, which is specifically relevant to the interpretation of *FMR1*, is shown in **Figure S22**. Sample sizes and study designs for all eight datasets are summarized in **Figure S23** and **Table S1**.

To ensure that each dataset was assigned to the appropriate analytical stratum, we verified its study design against the original source publication rather than relying on abbreviated catalog trait labels or prior aggregated dataset listings. This step is important because case-case (CC-GWAS) contrast statistics can carry case/control-labeled sample-size metadata that appears numerically consistent with a standard case-control design despite representing a comparison between two affected populations (Nudel et al., 2021; Peyrot & Price, 2021). For example, GWAS Catalog accession GCST90016603 compares 18,381 individuals with ASD against 4,819 individuals with Tourette syndrome rather than against unaffected population controls. Furthermore, the ASD individuals in this study are the same patients used in the primary ASD GWAS. Because of this patient overlap and the use of a disease-control group, this dataset cannot be treated as an independent population case-control study. Accordingly, we meta-analyzed two population case-control ASD summary statistic datasets: the iPSYCH-PGC 2017 freeze (Grove et al., 2019; OpenGWAS ID ieu-a-1185) and FinnGen. The three CC-GWAS contrast datasets (ASD-versus-Tourette syndrome, ASD-versus-OCD, and ASD-versus-MDD) were designated as a distinct analytic stratum. These contrasts were excluded from the case-control meta-analysis and genetic correlation testing but were retained for transcriptome imputation and connectivity mapping to serve as biologically distinct comparators.

Raw GWAS-VCF files were parsed using a custom ‘cyvcf2’-based converter that extracted variant identifier, chromosome, position, effect and non-effect allele, *p*-value (from the -log_10_(*p*) scaled LP field), *z*-score, effect size (ES), and standard error (SE) for each variant and exported to tab-separated tables (https://github.com/zwerhu/ASD_FXS). The genome build was GRCh37/hg19 for all input datasets.

### Fixed-effects meta-analysis of ASD case-control GWAS

The two population case-control ASD datasets (iPSYCH-PGC 2017 and FinnGen) were combined using inverse-variance-weighted (IVW) fixed-effects meta-analysis (Willer et al., 2010). Variants were merged by reference SNP ID (rsID), and allele alignment was harmonized between studies to ensure consistent effect and non-effect alleles. Same-orientation or swapped-orientation matches were retained, with sign-flipping of the effect estimate as appropriate. Strand-ambiguous (A/T or C/G) variants were excluded to prevent alignment errors. The EBI GWAS Catalog harmonization pipeline had already removed such variants from the iPSYCH-PGC input, so this exclusion affected only SNPs contributed by the FinnGen dataset.

For each retained SNP, the meta-analyzed effect size was computed as the precision-weighted mean of the two study-specific effect sizes, with weights *w_i_* = 1/SE_i_^2^. Cochran’s Q and I^2^ statistics were computed to quantify between-study heterogeneity (Willer et al., 2010). Genomic control inflation (λ; Yang et al., 2011) was estimated as the ratio of the median observed χ^2^ statistic to the expected median under the null, 0.4549 for one degree of freedom.

### Genetic overlap between population ASD risk and MZ-twin symptom discordance

To test whether the meta-analyzed population ASD case-control signal shares common-variant architecture with the MZ-twin symptom discordance GWAS, we did not use LD Score Regression (LDSC; Bulik-Sullivan et al., 2015). Instead, we implemented an LD-aware proxy analysis consisting of two complementary tests.

First, we performed genome-wide distance-based pruning, retaining one variant site per one Mb window per chromosome. Variant sites were selected independently of the significance of either trait to avoid selection-induced bias (Zollner & Pritchard, 2007). Within this approximately LD-independent variant site set, we computed the Pearson correlation and sign-concordance rate of *z*-scores between the meta-analyzed ASD signal and each twin-discordance GWAS. Because physical distance does not guarantee LD independence, this analysis was intended to reduce, rather than completely eliminate, redundancy arising from local LD. Second, because genome-wide pruned correlation can be underpowered to detect concordance concentrated at a small number of strongly associated loci (Dudbridge, 2013; Shi et al., 2016), we additionally identified approximately independent lead loci among variant sites with *p* < 10⁻^5^ in the ASD meta-analysis. Independence was defined operationally by physical genomic distance rather than by direct estimation of LD (e.g., *r*^2^ or *D*′; Hill & Robertson, 1968; Devlin & Risch, 1995) from a matched reference panel. We partitioned the genome into non-overlapping one Mb windows and retained the variant site with the lowest ASD meta-analysis *p*-value within each window. Because local recombination rate and LD decay vary substantially across the genome, this distance-based procedure approximates linkage equilibrium and may have residual correlation between loci in regions of long-range LD or unnecessarily split a single LD block that spans a window boundary. We therefore interpret this analysis as a distance-pruned lead-locus analysis rather than a formal LD-based conditional analysis. We explicitly contrasted these distance-pruned analyses with a naïve analysis using all nominally associated variant sites without pruning. The latter can be dominated by genomic loci containing many highly correlated variant sites in strong LD, producing inflated apparent concordance.

### Transcriptome imputation with S-PrediXcan

Tissue-specific genetically regulated gene expression was imputed for all eight traits across 49 GTEx v8 tissues using S-PrediXcan (Barbeira et al., 2018) with MASHR-based prediction models (GTEx Consortium, 2020). We used all 49 GTEx tissues rather than restricting the analysis to brain tissues for two main reasons. First, GTEx brain tissues generally have relatively small sample sizes, with most brain regions including only approximately 100-200 donors, which limits the statistical power of brain-specific gene-expression prediction models. Because many regulatory variants (eQTLs) have shared effects across tissues, integrating signals across all 49 tissues using ACAT can improve the power of gene-level association testing without restricting discovery to relatively underpowered brain tissues. Second, our FMRP signal was derived from plasma protein pQTLs and therefore represents a peripheral molecular phenotype. We instead adopted a tissue-agnostic approach, allowing the data to identify the most relevant tissue for each gene.

For each gene-trait pair, we defined the best_tissue as the GTEx tissue yielding the smallest S-PrediXcan association *p*-value across all 49 tissues tested. This approach avoids assuming in advance that brain tissue is necessarily the sole biologically relevant context. The GTEx v8 MASHR models and their associated SNP covariance matrices use GRCh38 variant IDs (chr_pos_ref_alt_b38), whereas the input GWAS summary statistics used rsID and GRCh37 coordinates. We therefore constructed an rsID-to-variant-ID translation table directly from the union of the weight tables for all 49 tissue models, yielding 217,195 unique mappings (https://github.com/zwerhu/ASD_FXS). This approach avoided the need for explicit coordinate liftover. The summary statistics for each trait were re-keyed to GTEx variant IDs through this mapping before S-PrediXcan execution.

To ensure compatibility with NumPy ≥ v2.0, we applied two targeted patches to the S-PrediXcan reference implementation (MetaXcan, commit current as of 2024) before execution. The first patch coerces legacy ‘numpy.matrix’ covariance objects to standard arrays via ‘numpy.asarray’ before linear algebra operations, ensuring that matrix-vector products used in variance calculations return scalars as expected under current NumPy behavior. The second patch adds a dimensionality guard to the diagnostic eigenvalue check for near-singular covariance matrices, accommodating genes with a single-SNP prediction model for which the covariance matrix is represented as a zero-dimensional array. The patched codebase was validated against the reference implementation and produced numerically identical results in all cases. Using this patched pipeline, we executed 343 tissue-by-trait S-PrediXcan runs (49 tissues × 7 traits from the initial analysis), followed by an additional 49 runs for the corrected ASD case-control trait, yielding 3,822,889 valid gene-tissue-trait association statistics after quality filtering.

We performed a simplified SNP-level consistency check for selected top genes as a lightweight complement for formal statistical localization. *XRN2* was the top gene for ASD_caseControl_iPSYCH2017, whereas *PPP3CA* and *TNRC18* were the top two genes for FMRP_protein. These three genes are located on chromosomes 20, 4, and 7, respectively. *PPP3CA* and *TNRC18* had the strongest association signals after integration across tissues for the FMRP_protein trait. For each gene, we identified the single SNP comprising its MASHR expression-prediction model in the relevant tissue and examined its association statistics in the corresponding trait GWAS, together with the regional GWAS association pattern within a window spanning 250 kb upstream and downstream of the SNP. Regional association was summarized as both the absolute number and proportion of tested SNPs reaching a suggestive threshold (*p* < 10^-5^). The proportion accounts for differences in local SNP density and genotyping imputation coverage across loci, allowing the three loci to be compared on a common scale, whereas the absolute count reflects the overall size of the associated regional signal. Considering both metrics together allowed us to assess whether the model SNP falls within a multi-SNP association peak rather than representing an isolated single-SNP signal, which would provide weaker support for a shared causal variant between the expression-prediction model and the GWAS signal.

This approach does not yield a formal statistical estimate of the probability that the GWAS and expression signals share a single causal variant, such as that produced by dedicated colocalization methods including COLOC (Gamazon et al., 2015; Giambartolomei et al., 2014) and cannot distinguish true colocalization from association driven by LD with an untested causal variant. This analysis was therefore intended as a directional exploratory check without requiring the full eQTL summary statistics.

### Cross-tissue integration and pathway enrichment

For each trait, gene-level association signals were integrated across all tested GTEx tissues using the aggregated Cauchy association test (ACAT), a *p*-value combination method that can accommodate dependence among component tests without requiring the correlation structure to be explicitly modeled (Liu et al., 2019). This property is useful given the substantial cross-tissue correlation inherent to GTEx expression-prediction models. Genes located within the extended major histocompatibility complex (MHC) region were excluded from integrative and pathway analyses. The chr6p21 cytogenetic band gene set was used as an exclusion proxy because genomic coordinates were not available in the S-PrediXcan gene-level output. GSEA was performed for each trait against the 50 MSigDB Hallmark gene sets (Subramanian et al., 2005) using the ‘gseapy’ pre-ranked implementation with 1,000 permutations. Genes were ranked by signed ACAT-combined significance, calculated as -log_10_(*p*) multiplied by the sign of the mean cross-tissue *z*- score.

### Connectivity mapping and drug repurpose

Level 5 moderated *z*-score, replicate-collapsed CMap LINCS L1000 data from the Broad Institute (GSE92742) were filtered to small-molecule compound treatment signatures (pert_type = "*trt_cp*") flagged by the Broad Institute as exemplar signatures selected through its quality-control pipeline for each unique combination of perturbagen, cell line, dose, duration, and time point (112,495 of 473,648 total signatures). Given computational constraints on available memory and storage, signatures were further aggregated to one representative signature per unique compound by selecting the signature with the highest transcriptional activity score (TAS), yielding 19,730 unique compound signatures. This aggregation collapses transcriptional responses across cell lines into a single representative profile per compound and therefore prioritizes the cellular context in which each compound produced its strongest transcriptional response. The resulting under-representation of neural-lineage cell lines in this aggregated profile is a limitation discussed further in Discussion section. All analyses were restricted to the 11,141 genes present in both the CMap L1000 best-inferred gene space and the union of genes tested across all eight S-PrediXcan trait analyses.

For each of the eight traits, we computed five complementary connectivity metrics against each of the 19,730 compound signatures, following the classical CMap methodology (Lamb et al., 2006) and its extensions: (1) Pearson correlation across all 11,141 genes; (2) Spearman correlation across all genes; (3) a signed KS connectivity score computed from the positions of the top and bottom *N* disease-associated genes within the rank-ordered compound signature, following the original CMap enrichment-score formulation; (4) extreme Pearson correlation restricted to the top and bottom *N* disease-associated genes; and (5) extreme Spearman correlation over the same restricted gene set. Metrics 3-5 were computed at four gene-set size thresholds (*N* = 50, 100, 250, and 500) and averaged. All five metrics and their associated per-trait rank statistics were combined into a single mean rank for each compound-trait pair, with more negative reversal- direction scores corresponding to a lower, more favorable combined rank. The analysis involved 19,730 compounds across eight traits, yielding approximately 158,000 compound-trait combinations. Because each combination required genome-wide rank computations, all connectivity metrics were implemented as vectorized NumPy operations and processed in compound batches of 1,500 to limit peak memory usage.

Connectivity scores were standardized for prioritization by converting the KS-signed score of each compound to a *z*-score relative to the empirical distribution of KS-signed scores across all compounds evaluated against the same disease signature (Lamb et al., 2006; Subramanian et al., 2017). This empirical normalization evaluates the relative strength of each compound within the tested library rather than assigning gene-level *p*-values. We considered a conventional per-gene correlation significance framework during method development. However, because genome-wide compound-disease expression correlations are typically modest in magnitude and gene expression measurements exhibit substantial correlation structure, this approach provided limited discrimination after multiple-testing correction. We therefore adopted the empirical *z*- score framework for prioritization.

Compounds were considered candidate reversal signatures if their empirical *z*-score was less than -2 and all five connectivity metrics were concordantly negative. We emphasize that this is a nominal, exploratory threshold rather than a formal multiple-testing correction criterion. The five connectivity metrics are themselves correlated, so requiring concordance across all five reduces the candidate set only modestly relative to the *z* < −2 threshold alone, from 3,927 to 2,788 compound-trait pairs (29% reduction). MOA annotations were obtained for compounds with resolvable common names by matching to the Broad Institute Drug Repurposing Hub annotation table (data version 2020-03-24; 4,382 unique annotated compounds), providing MOA information for 1,773 of 19,730 (9.0%) compounds in the tested library. MOA enrichment among significant reversal candidates was assessed by Fisher’s exact test for each elementary MOA term. Combination MOA annotations were decomposed into constituent terms. For each term, the proportion of significant candidates annotated with that MOA was compared with the corresponding proportion among all tested compounds with any available MOA annotation, followed by Benjamini-Hochberg FDR correction across all tested MOA terms (Benjamini & Hochberg, 1995).

To characterize the mechanistic organization of the significant candidate pool, we performed hierarchical clustering using Ward linkage on Euclidean distance for the top 30 MOA-annotated candidates. Candidates were selected by cross-trait convergence and best within-trait rank among the top 100 reversal candidates per trait. Clustering was based on 1 minus the Spearman correlation between the eight-trait KS *z*-score connectivity profiles of each pair of compounds. For visualization purposes only, each candidate was additionally classified into a coarse MOA class (e.g., serotonergic, dopaminergic, adenosine/purinergic, or kinase inhibitor) by keyword matching against its Broad Repurposing Hub MOA annotation.

### Software and computational environment

All analyses were performed in Python 3.12 using ‘pandas’, ‘numpy’, ‘scipy’, ‘statsmodels’, ‘cyvcf2’, ‘cmapPy’, ‘gseapy’, and the MetaXcan/S-PrediXcan reference implementation, which was locally patched as described above. Network and layered-flow visualizations were produced with ‘networkx’, ‘matplotlib’, and ‘plotly’. Given the scale of several intermediate datasets, including individual GWAS files up to 16 million variants sites and CMap Level 5 data exceeding 20 GB uncompressed, memory-bounded chunked-processing implementations were used throughout in preference to whole-dataset in-memory operations.

## Data Availability

GWAS summary statistics are publicly available via the IEU OpenGWAS project (https://gwas.mrcieu.ac.uk) and the NHGRI-EBI GWAS Catalog (https://www.ebi.ac.uk/gwas/). CMap LINCS L1000 Level 5 data are available from the Broad Institute (GSE92742). GTEx v8 MASHR prediction models are available via predictdb.org. All the summary statistics derived, integrated gene-level association results, and connectivity-mapping outputs generated in this study are available upon requests to corresponding author. Pipeline of the code implementation along with processed data is available at GitHub repository https://github.com/zwerhu/ASD_FXS.

## Acknowledgements

We thank the iPSYCH-PGC Autism Working Group, FinnGen, the Within Families Consortium, the INTERVAL study investigators, and the Broad Institute Connectivity Map/LINCS team for making the summary-level data underlying this analysis publicly available.

