## supplementary materials for "Integrative genetic and transcriptomic analysis identifies candidate genes and compounds for Fragile X syndrome and autism spectrum disorder"

This document provides 23 supplementary figures and eight supplementary tables extending the analyses reported in the main text, including additional quality-control diagnostics, extended drug repurposing and pathway enrichment results, a brain-tissue-restricted drug-disease-gene-tissue analysis, and a literature-based validation check of top drug repurposing candidates against the published FXS and ASD pharmacology literature.

### Supplementary Figures

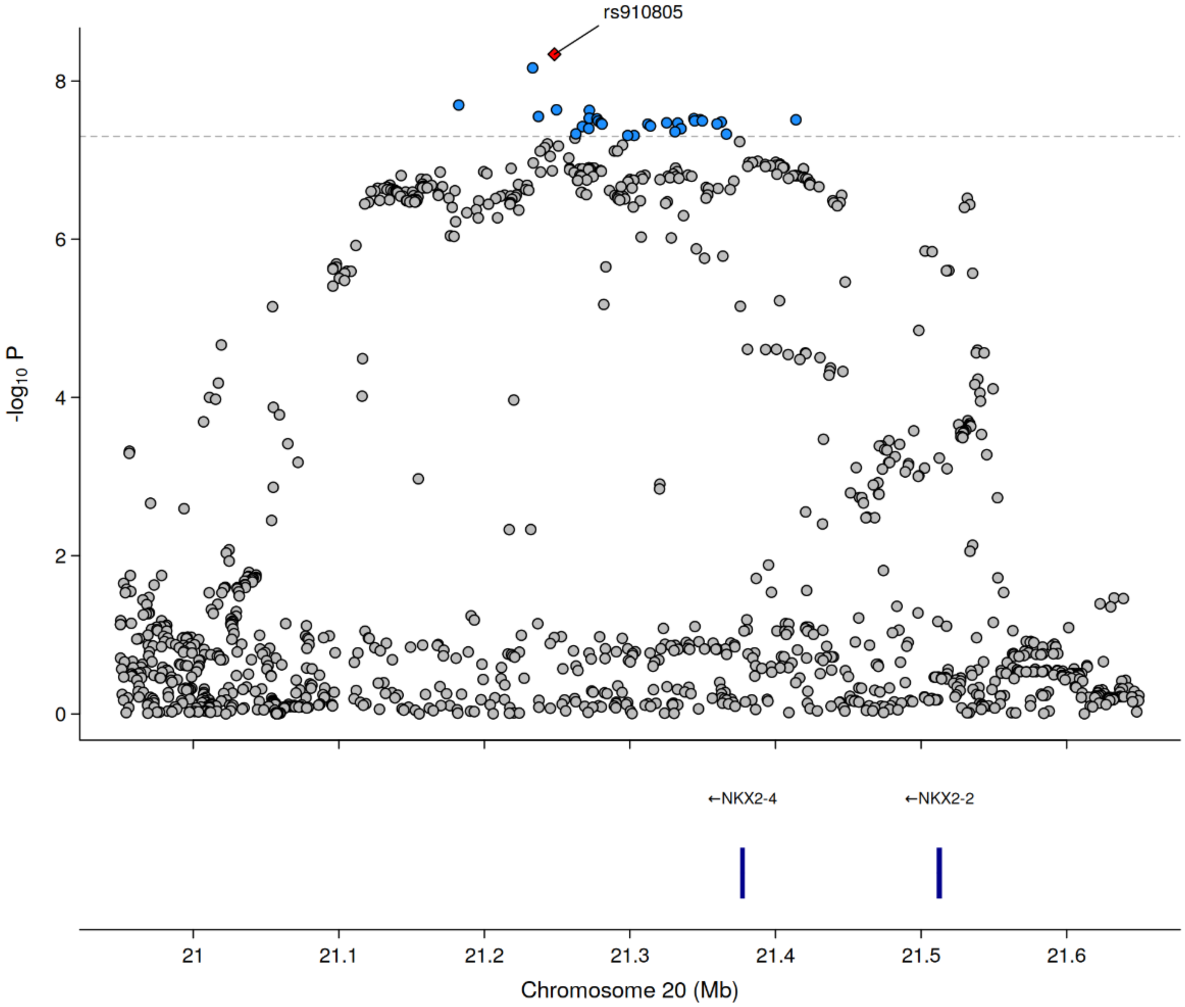


**Figure S1. Regional association plot of the chromosome 20 locus.** Regional association plot of the corrected ASD case-control meta-analysis (iPSYCH-PGC 2017 + FinnGen) spanning chromosome 20:20.95–21.65 Mb (GRCh37). The top panel shows each tested SNP plotted by genomic position (*x*-axis) against $-\log_{10}(p)$ from the fixed-effects inverse-variance-weighted meta-analysis (*y*-axis). The index SNP, rs910805 ($p = 4.6\times10^{-9}$), is highlighted with a red diamond. SNPs reaching genome-wide significance ($p < 5\times10^{-8}$) are shown in blue, and all other SNPs are shown in gray.

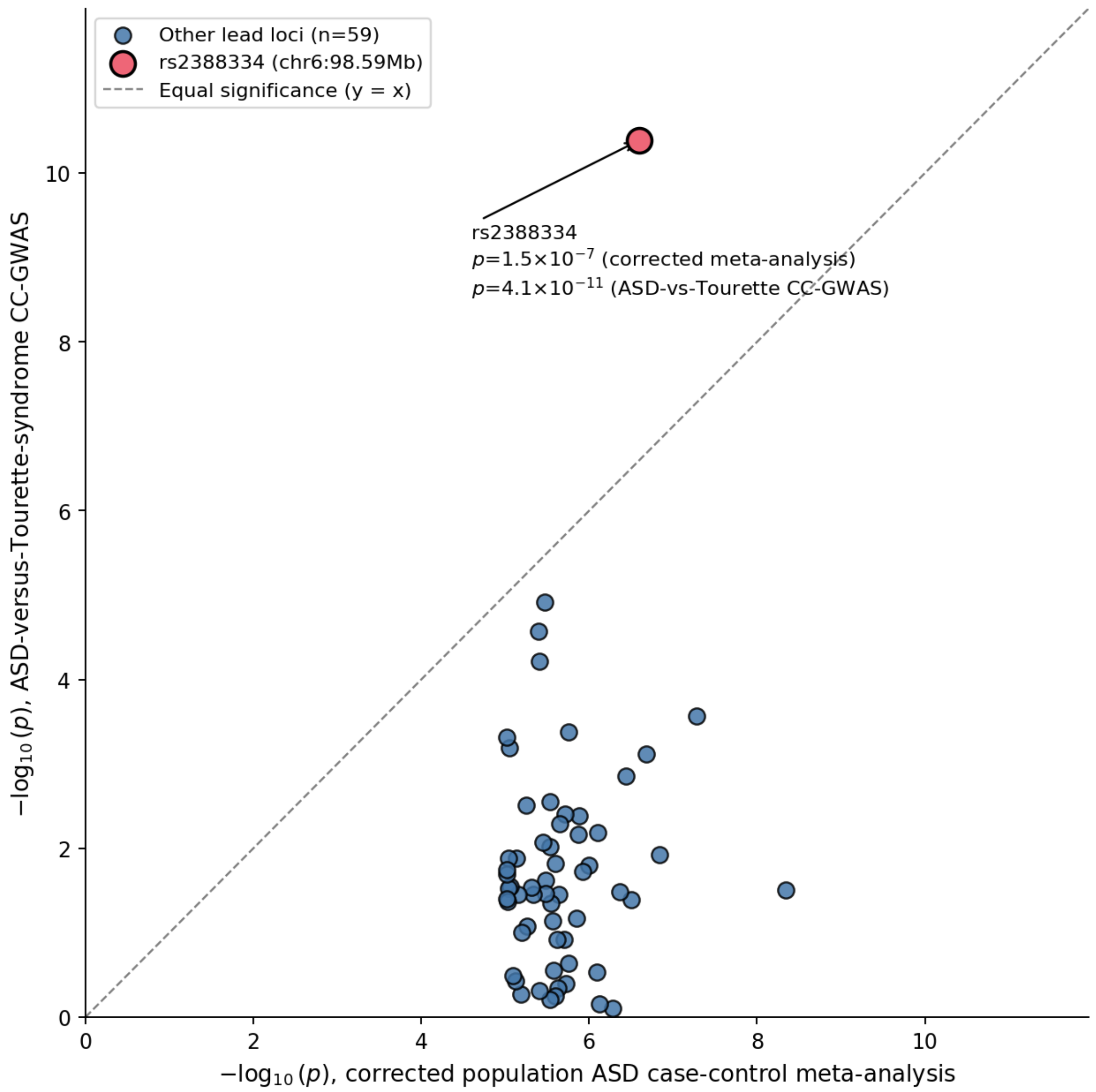


**Figure S2. Interpretation of the chromosome 6 signal under the correct case-case GWAS design.** For each of the 60 lead loci from the corrected ASD meta-analysis (Table S2) with available association statistics in the ASD-versus-Tourette-syndrome CC-GWAS, the significance of the population case-control association ($x$-axis) is plotted against the significance of the case-case association ($y$-axis). The dashed diagonal indicates equal significance in both analyses. Fifty-nine of the 60 loci fall below the diagonal, showing the expected pattern of a weaker signal in the case-case contrast than in the population case-control meta-analysis. The single exception, rs2388334 (chr6:98.59 Mb, highlighted in red), shows stronger significance in the case-case contrast ($p = 4.1\times10^{-11}$) than in the corrected population case-control meta-analysis ($p = 1.5\times10^{-7}$, below genome-wide significance).

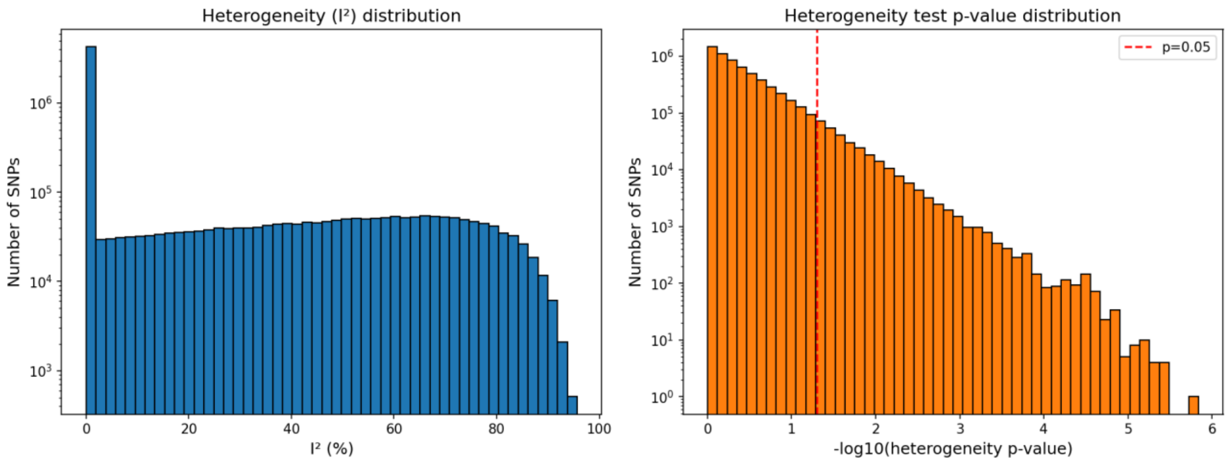


**Figure S3. Heterogeneity across the corrected ASD case-control meta-analysis.** Distribution of the $I^2$ heterogeneity statistic (left) and the corresponding Cochran's Q heterogeneity test *p*-value (right) across all 6,159,907 SNPs tested in the corrected fixed-effects meta-analysis (iPSYCH-PGC 2017 + FinnGen). $I^2$ quantifies the proportion of total variance in SNP effect-size estimates attributable to heterogeneity between the two contributing studies rather than sampling error and ranges from 0% to 100%. The right panel shows the corresponding heterogeneity test *p*-value on the $-\log_{10}$ scale, with the dashed red line marking the nominal $p = 0.05$ threshold.

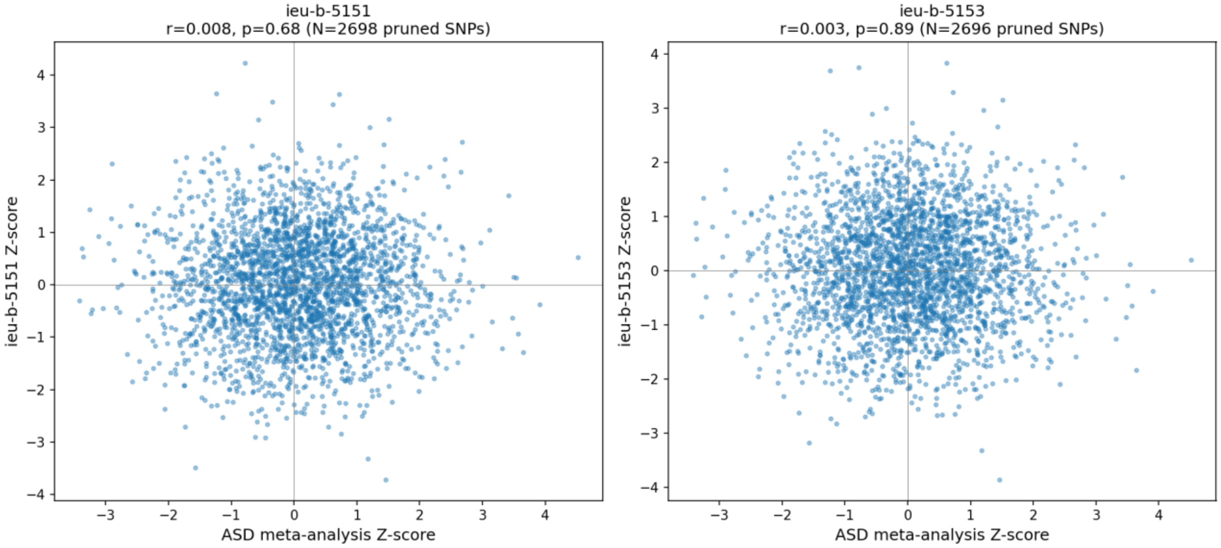


**Figure S4. Genome-wide *z*-score concordance after LD pruning between the ASD case-control meta-analysis and each MZ-twin symptom-discordance GWAS.** *z*-scores from the corrected population ASD meta-analysis (x-axis) are plotted against z-scores from each MZ-twin symptom-discordance GWAS (y-axis) for MZtwin_ASD_all (ieu-b-5151, left; N=2,698 pruned SNPs) and MZtwin_ASD_children (ieu-b-5153, right; N=2,696 pruned SNPs), following the LD-pruning procedure described in Methods section.

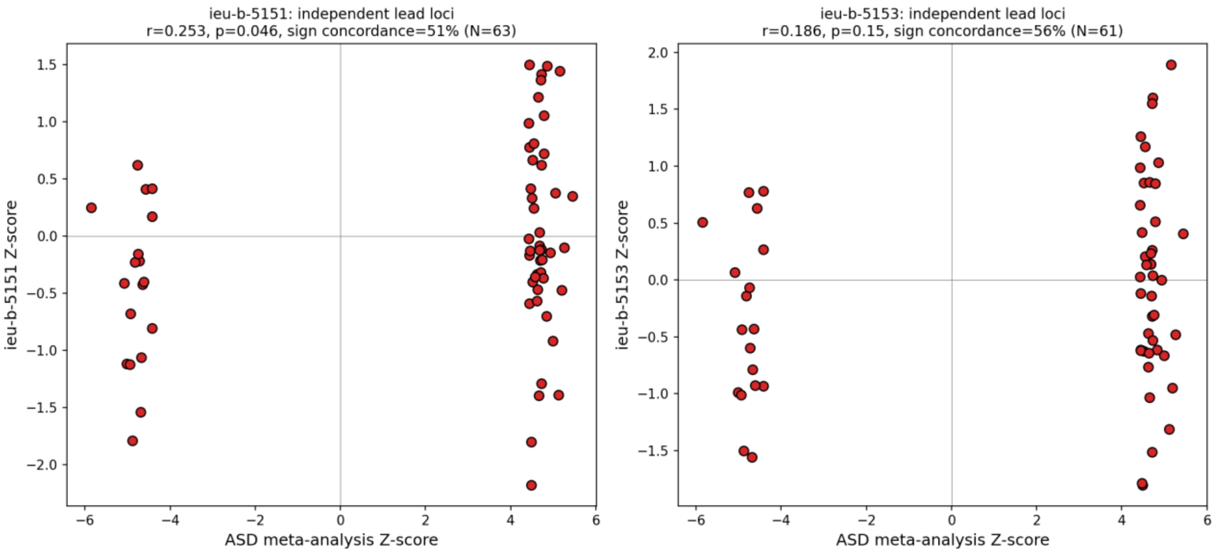


**Figure S5. *Z*-score concordance at independent lead variant sites between the ASD meta-analysis and each MZ-twin symptom-discordance GWAS.** As in Figure S4, but restricted to lead variants from the corrected ASD case-control meta-analysis reaching $p < 10^{-5}$ and selected using the distance-based pruning procedure described in the Methods. Results are shown for MZtwin_ASD_all (ieu-b-5151, left; $N = 63$ loci) and MZtwin_ASD_children (ieu-b-5153, right; $N = 61$ loci). Pearson correlation was $r = 0.253$ ($p = 0.046$; sign concordance = 51%) for MZtwin_ASD_all and $r = 0.186$ ($p = 0.15$; sign concordance = 56%) for MZtwin_ASD_children.

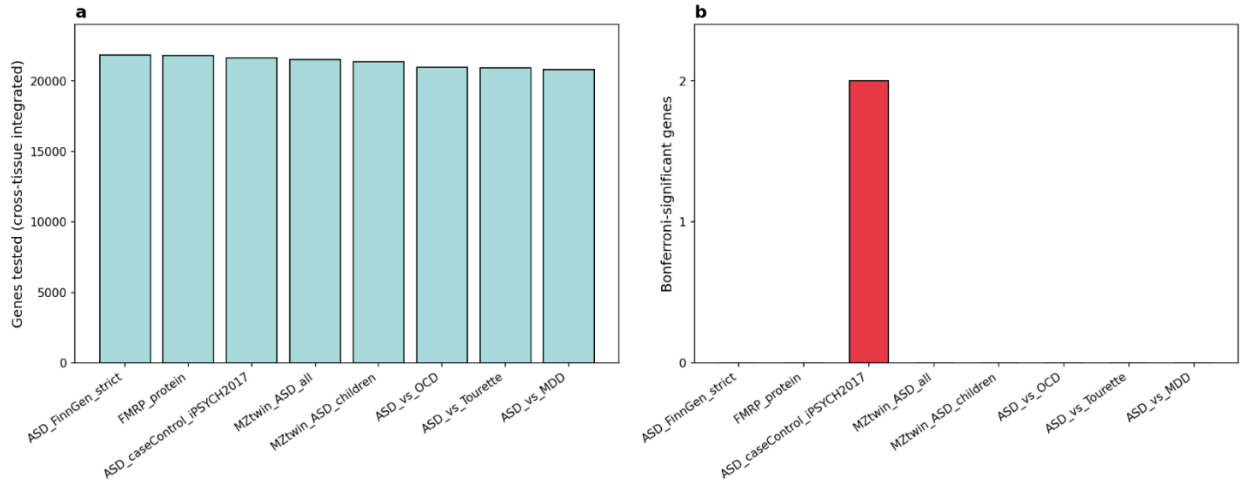


**Figure S6. Number of genes entering cross-tissue ACAT integration and number reaching Bonferroni significance, per trait.** (a) Number of genes with ACAT *p*-values after cross-tissue integration and exclusion of the MHC region, ranging from 20,807 (ASD_vs_MDD) to 21,846 (ASD_FinnGen_strict). (b) Number of genes reaching trait-specific Bonferroni significance ($\alpha$ = 0.05 divided by the number of genes tested for that trait). Only ASD_caseControl_iPSYCH2017 yielded Bonferroni-significant genes ($n$ = 2).

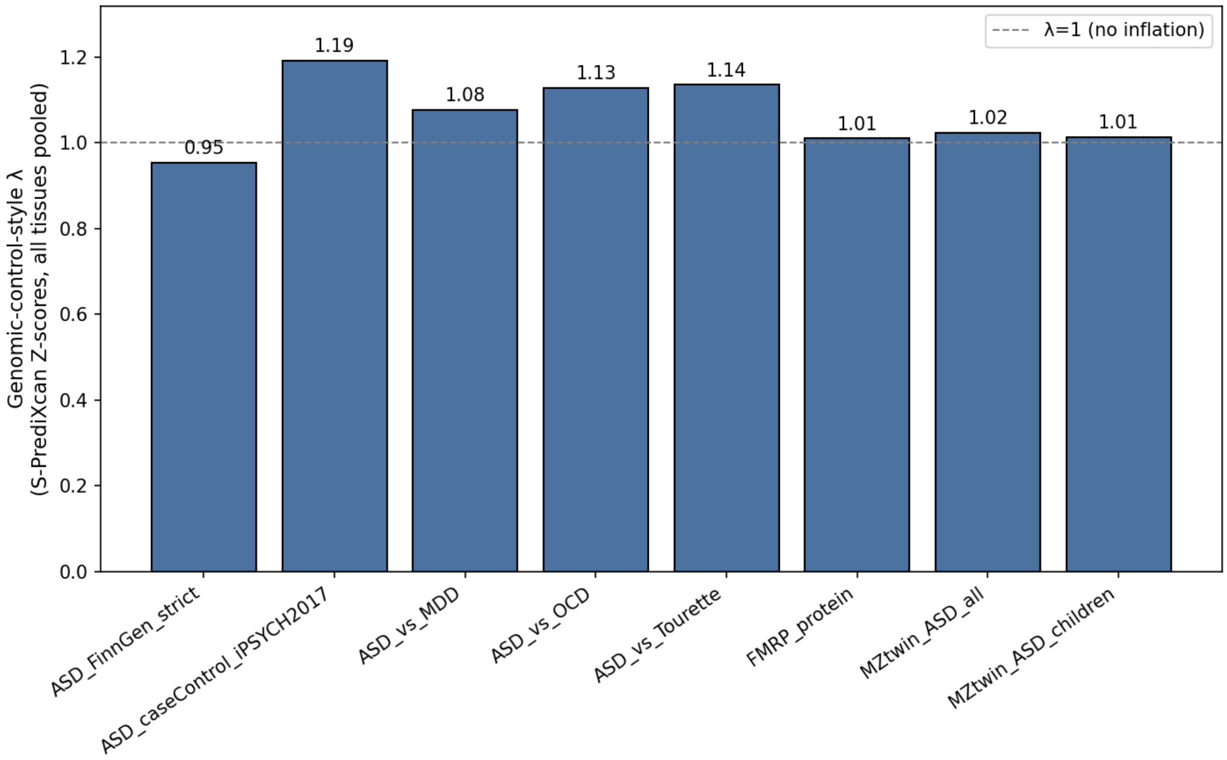


**Figure S7. Genomic control-style inflation statistic for pooled S-PrediXcan *z*-scores across tissues.** The inflation statistic ($\lambda$ = median($z^2$)/0.4549) is shown for each trait. Values cluster near $\lambda = 1$ (dashed line) for most traits, ranging from 0.95 (ASD_FinnGen_strict) to 1.19 (ASD_caseControl_iPSYCH2017), indicating minimal to modest inflation in the pooled cross-tissue S-PrediXcan test statistics. This pattern is consistent with the modest test-statistic inflation but does not by itself establish a source of inflation.

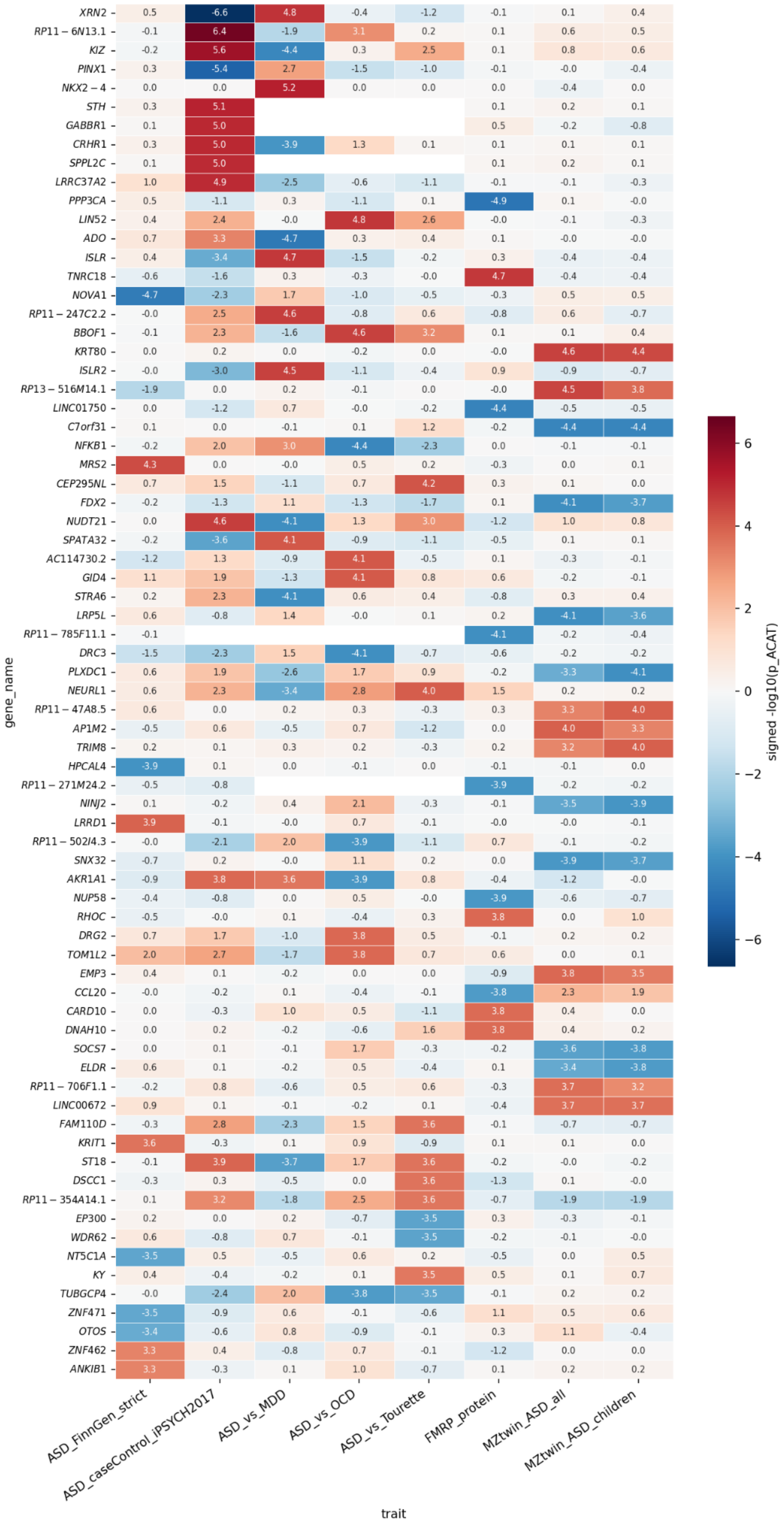

XRN2
RP11 – 6N13.1
KIZ
PINX1
NKX2 – 4
STH
GABBR1
CRHR1
SPPL2C
LRRC37A2
PPP3CA
LIN52
ADO
ISLR
TNRC18
NOVA1
RP11 – 247C2.2
BBOF1
KRT80
ISLR2
RP13 – 516M14.1
LINC01750
C7orf31
NFKB1
MRS2
CEP295NL
FDX2
NUDT21
SPATA32
AC114730.2
GID4
STRA6
LRP5L
RP11 – 785F11.1
DRC3
PLXDC1
NEURL1
RP11 – 47A8.5
AP1M2
TRIM8
HPCAL4
RP11 – 271M24.2
NINJ2
LRRD1
RP11 – 502I4.3
SNX32
AKR1A1
NUP58
RHOC
DRG2
TOM1L2
EMP3
CCL20
CARD10
DNAH10
SOCS7
ELDR
RP11 – 706F1.1
LINC00672
FAM110D
KRIT1
ST18
DSCC1
RP11 – 354A14.1
EP300
WDR62
NT5C1A
KY
TUBGCP4
ZNF471
OTOS
ZNF462
ANKIB1
gene_name
ASD_FinnGen_strict
ASD_caseControl_iPSYCH2017
ASD_vs_MDD
ASD_vs_OCD
ASD_vs_Tourette
FMRP_protein
MZtwin_ASD_all
MZtwin_ASD_children
trait
signed -log10(p_ACAT)
6
4
2
0
−2
−4
−6

**Figure S8. Extended gene-by-trait heatmap showing the top 10 cross-tissue genes for each trait.** As in **Figure 2**, each cell shows the signed $-\log_{10}(p\_ACAT)$, magnitude representing cross-tissue association significance and sign representing the direction of the mean cross-tissue S-PrediXcan $z$-score. White cells indicate genes that were not among the top 10 for that trait. The expanded view includes 73 unique genes and captures genes highlighted in **Figure 2**, including *XRN2*, *PPP3CA*, and *TNRC18*, alongside additional trait-specific top associations. Several genes, including *KRT80*, *AP1M2*, *TRIM8*, and *RP11-47A8.5*, show concordant signals specifically across MZtwin_ASD_all and MZtwin_ASD_children.

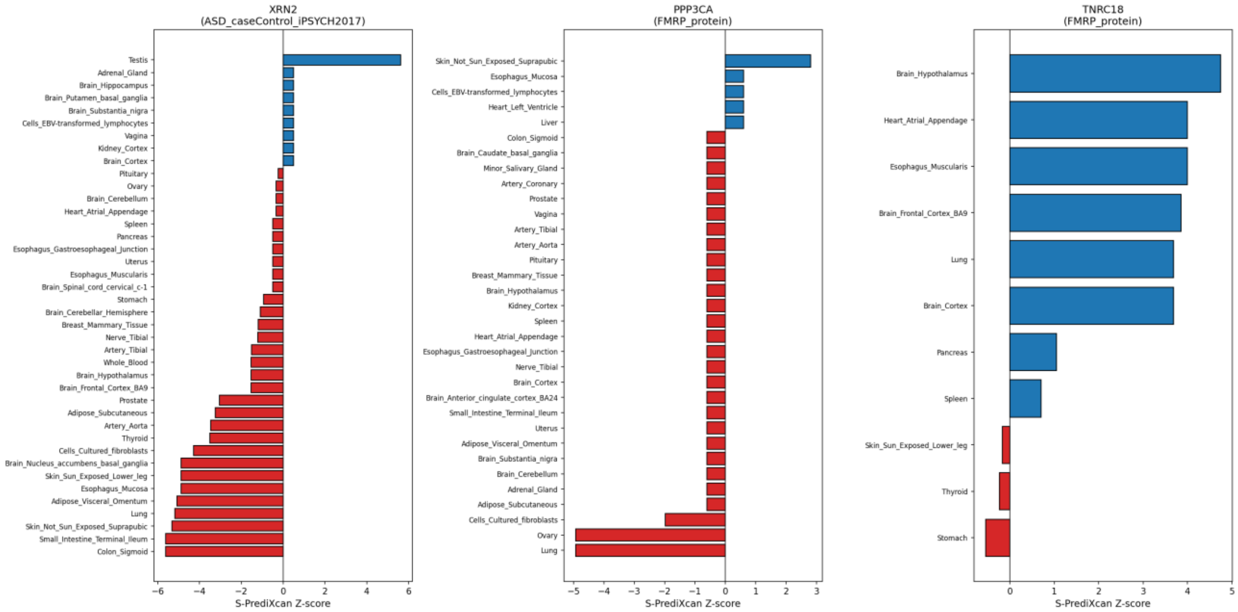


**Figure S9. Per-tissue S-PrediXcan *z*-scores for the three top cross-tissue genes.** Per-tissue S-PrediXcan *z*-sores are shown for *XRN2* for ASD_caseControl_iPSYCH2017 and *PPP3CA* and *TNRC18* for FMRP_protein across all GTEx tissues in which each gene was tested. *XRN2* and *PPP3CA* show a bidirectional patterns, with some tissues showing positive *z*-scores and the majority showing negative *z*-scores. *TNRC18* shows positive *z*-scores across nearly all 11 tissues in which it was tested. For all three genes, the tissue with the strongest absolute *z*-score (Colon_Sigmoid for *XRN2*, Lung for *PPP3CA*, and Brain_Hypothalamus for *TNRC18*) matches the best_tissue reported in the main text.

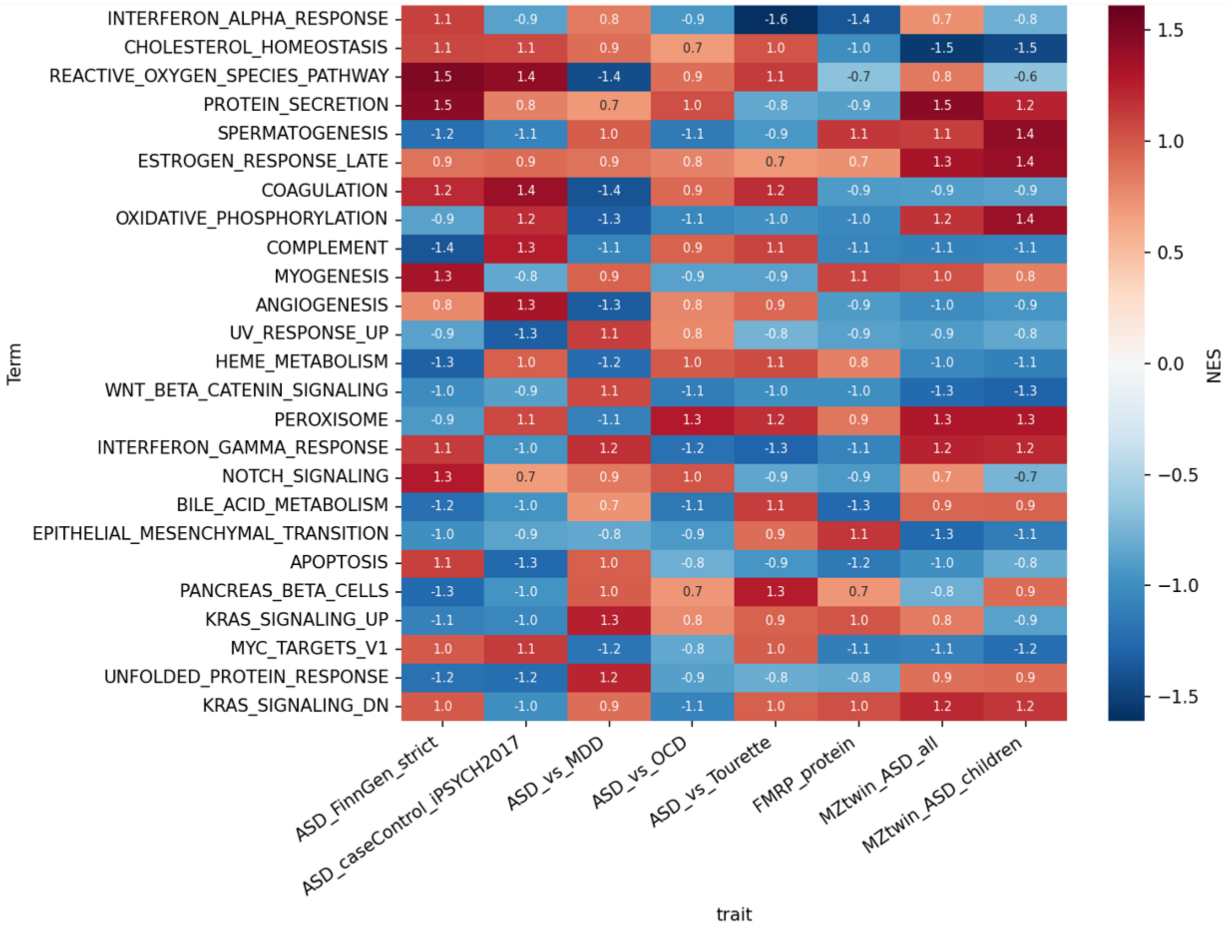


**Figure S10. Normalized enrichment scores for the top 25 Hallmark pathways.** Normalized enrichment scores (NES) are shown for the 25 hallmark pathways with the largest maximum absolute NES across the eight traits. This proves an extended view beyond the FDR < 0.25 threshold used to identify the pathways highlighted in **Figure 3**. Most trait-pathway pairs shown here do not reach FDR < 0.25 and are included solely because they rank among the top 25 by NES magnitude in at least one trait. The five pathways reaching FDR < 0.25 in **Figure 3** therefore reflect statistical support under the permutation-based testing framework rather than simply the largest NES values. Sign patterns vary across traits and pathways, with no single trait or pathway showing uniformly positive or negative enrichment.

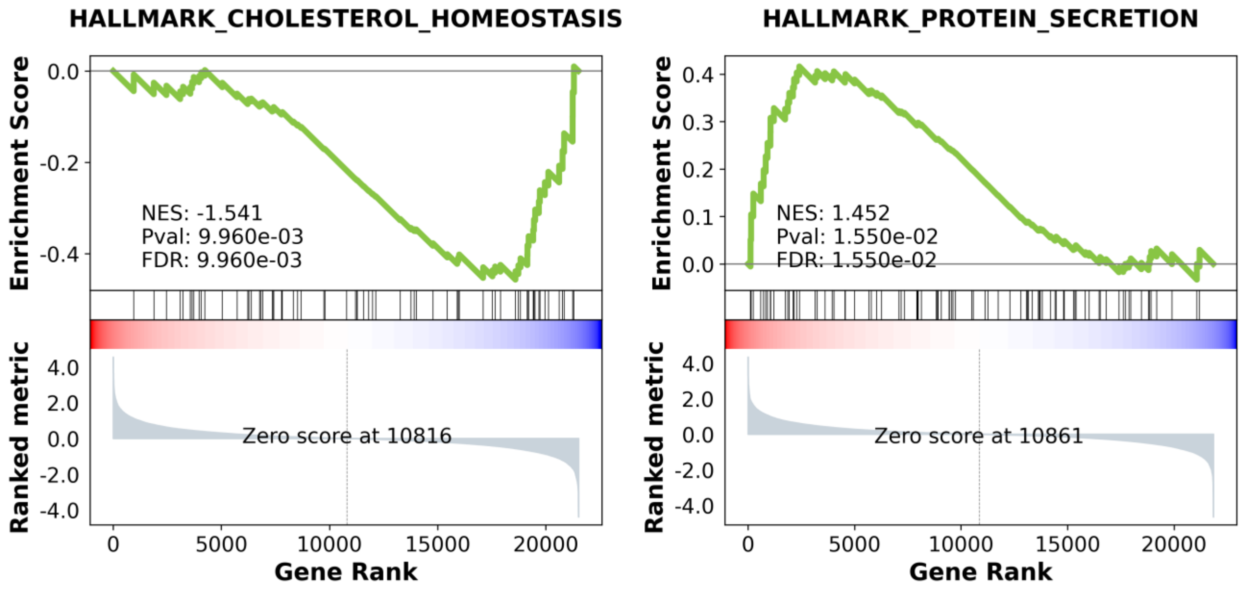


**Figure S11. Classical GSEA running enrichment score plots for two representative trait-pathway pairs.** MZtwin_ASD_all / HALLMARK_CHOLESTEROL_HOMEOSTASIS (left) and ASD_FinnGen_strict / HALLMARK_PROTEIN_SECRETION (right). The green curve shows the running enrichment score as the ranked gene list is traversed. Vertical black tick marks indicate the rank positions of pathway member genes, and the bottom panel shows the signed gene-level association statistic used to rank the genes. The dashed vertical line marks the point at which the ranked metric crosses zero. Cholesterol homeostasis shows negative enrichment with an NES of −1.541 and FDR = $9.96\times10^{-3}$, whereas protein secretion shows positive enrichment with an NES of 1.452 and FDR = $1.55\times10^{-2}$, consistent with the opposite-signed enrichment patterns for these two trait-pathway pairs.

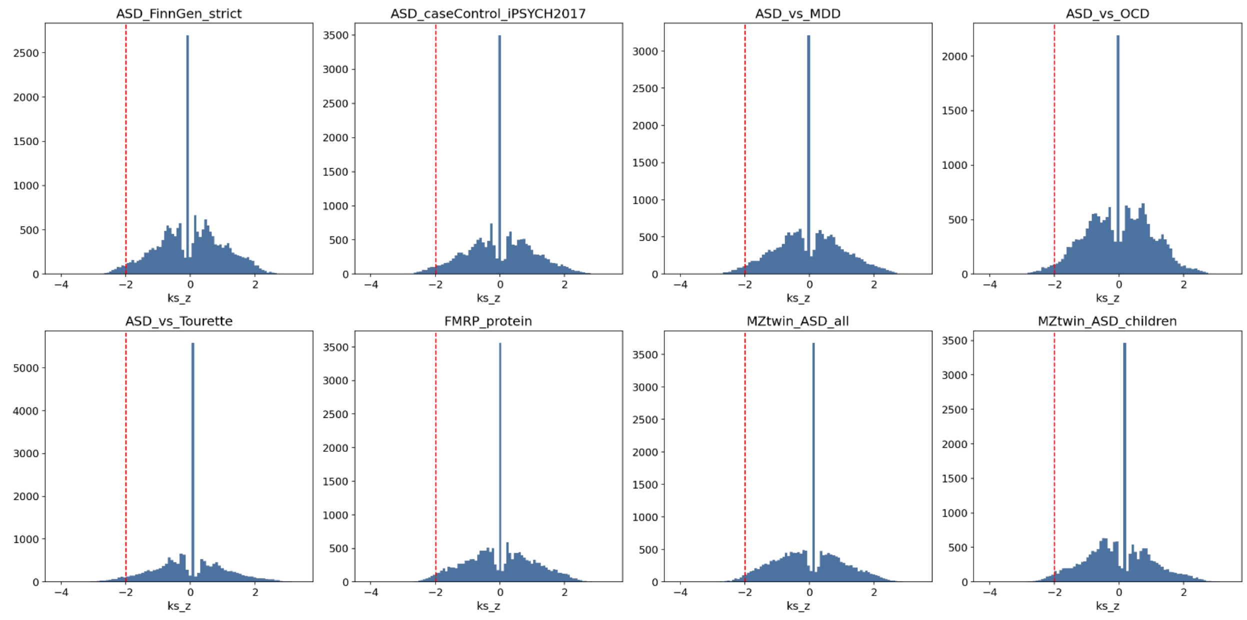


**Figure S12. Empirical distribution of the ks_z connectivity statistic across tested compounds.** The distribution is shown separately for each of the eight traits. The dashed red line indicates the reversal candidate threshold (ks_z < -2). All eight distributions show a similar overall shape, with a broad distribution around zero and a pronounced narrow peak near ks_z = 0. The ks_z < −2 threshold falls within the left tail of each distribution, consistent with the exploratory nature of the candidate-selection criterion described discussed in the main text. The broadly similar distributions across traits indicate that the threshold applied to comparable empirical score distributions, while differences in the candidate counts reflect variation in the underlying trait-specific connectivity signals.

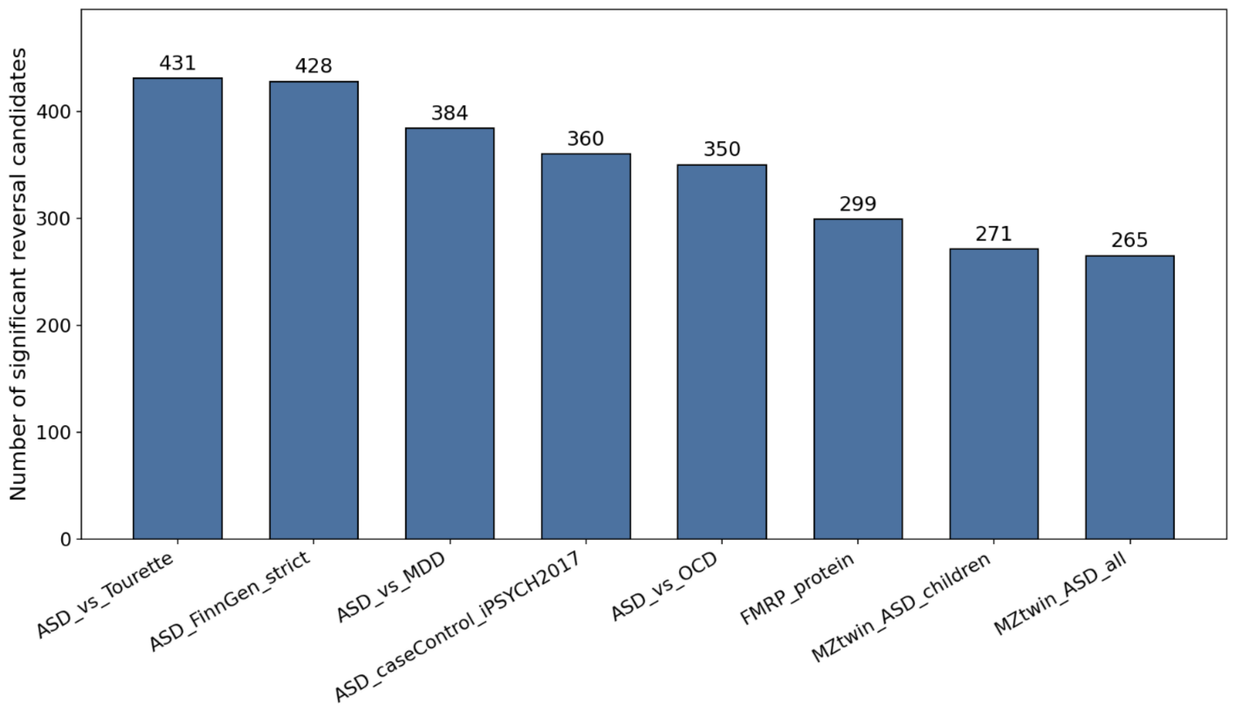


**Figure S13. Number of significant reversal candidate compounds identified per trait.** Candidate counts range from 265 (MZtwin_ASD_all) to 431 (ASD_vs_Tourette), summing to the 2,788 significant compound-trait pairs reported in the main text. The two MZ-twin symptom-discordance traits yield the fewest candidates. These differences may reflect differences in signal strength, sample size, or other properties underlying trait signatures. Candidate count does not scale simply with GWAS sample size across all eight traits. For example, ASD_caseControl_iPSYCH2017 (*N* = 46,350) yields fewer candidates than ASD_vs_OCD (*N* = 21,069), indicating that candidate counts depend on more than GWAS sample size alone.

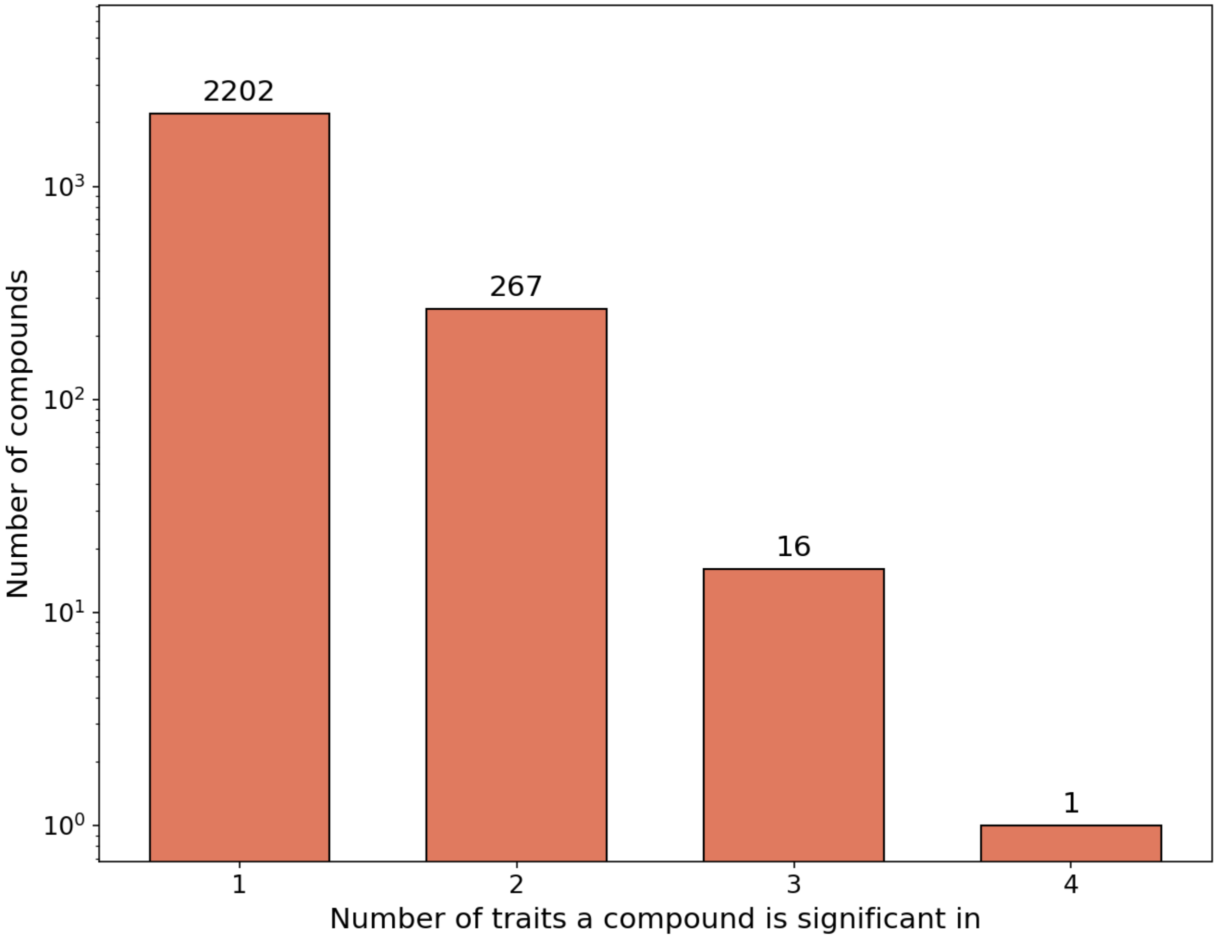


**Figure S14. Distribution of the number of traits in which each significant compound is identified as a reversal candidate.** The distribution shows that the 17 cross-trait-shared compounds represent a small subset of the full candidate pool. Most significant compounds are associated with a single trait, with progressively fewer compounds recurring across multiple traits. The majority of significant compounds (2,202 of 2,486 unique compounds, 88.6%) are significant in only one trait, while 267 are significant in two traits, 16 in three traits, and one in four traits. The 17 compounds significant in three or more traits therefore represent the small tail of the recurrence distribution rather than the typical pattern of compound identification.

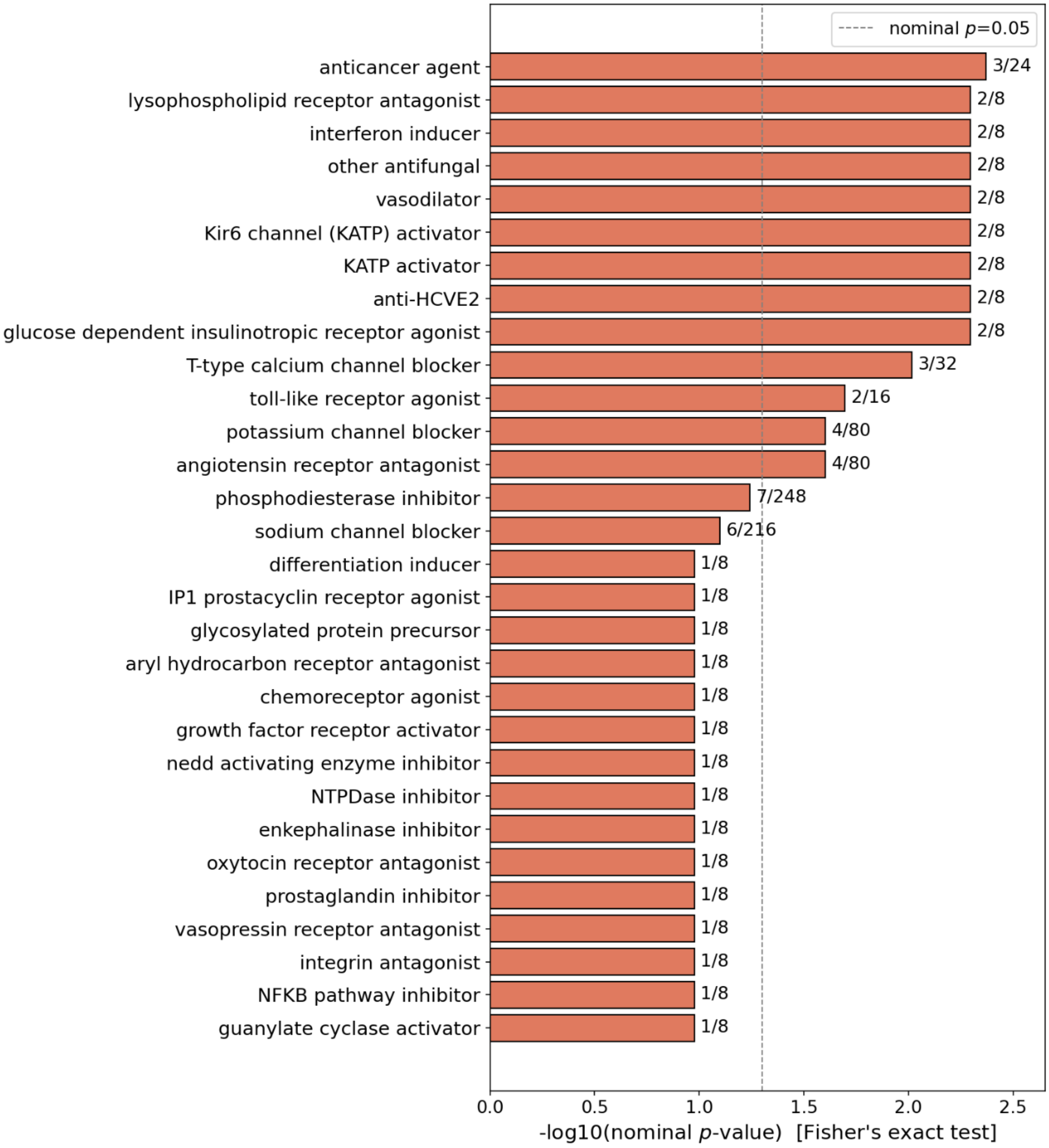


**Figure S15. Full mechanism-of-action enrichment results.** The top 30 elementary MOA terms are shown according to nominal Fisher's exact test $p$-value. Consistent with the main text, none of these MOA terms survive Benjamini-Hochberg FDR correction. The results are therefore presented as nominal, exploratory rankings rather than statistically confirmed enrichments. The lower portion of the ranking contains several terms with identical nominal $p$-values, arising from small and identical annotated-compound denominators. The two mechanistic classes highlighted in the main text as the strongest nominal enrichments not driven by such ties, T-type calcium channel blockade and angiotensin receptor antagonism, are among the higher-ranked terms. Glutamate receptor antagonism is not an enriched MOA class in this analysis because it is represented by a single significant compound, spaglumic acid (Table 2).

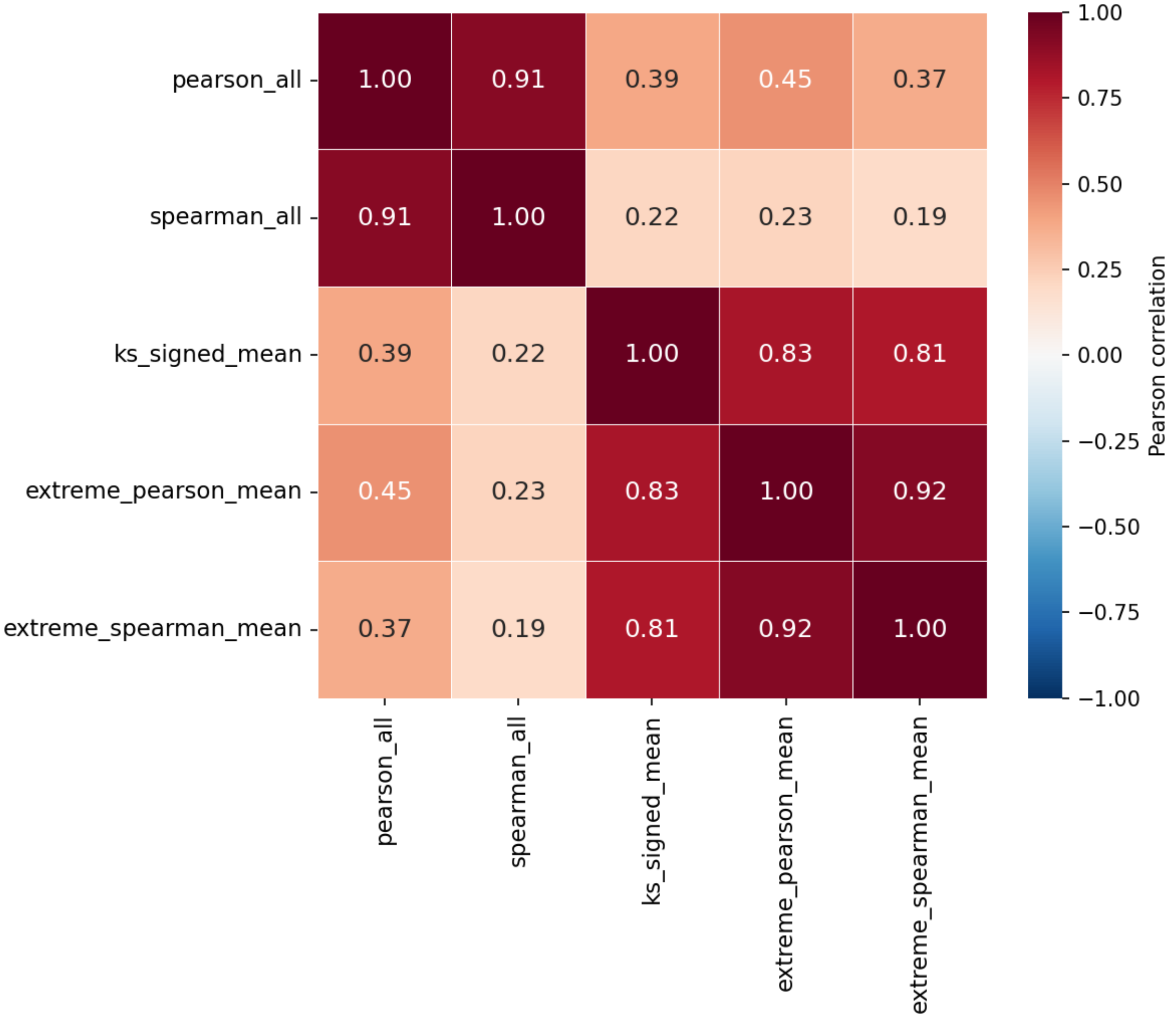


**Figure S16. Pearson correlation among the five connectivity metrics.** Pearson correlation among the five connectivity metrics (Pearson, Spearman, KS-signed, extreme Pearson, and extreme Spearman) is shown across compound-trait pairs. The five metrics form two broad clusters. The two full-signature correlation metrics (pearson_all and spearman_all; $r = 0.91$) form one cluster, while the KS-signed statistic and the two extreme-gene correlation metrics form a second, more tightly correlated cluster ($r = 0.81$–$0.92$). Correlations between the two clusters are weaker ($r = 0.19$–$0.45$). These patterns confirm that the five metrics are correlated rather than independent tests, supporting the interpretation in the main text that require concordance across all five metrics provides additional but limited stringency beyond the ks_z < −2 criterion. All pairwise correlations are positive.

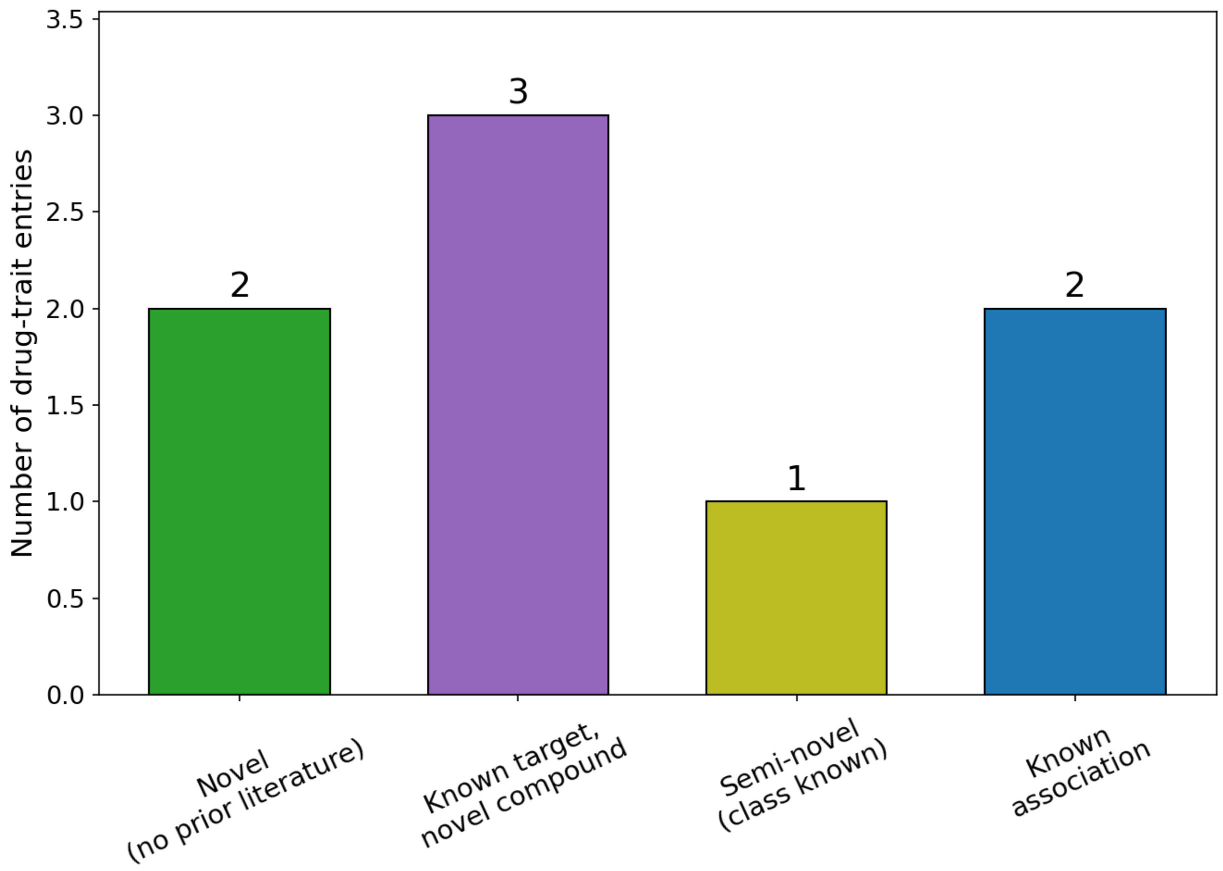


**Figure S17. Literature validation status of manually evaluated drug-repurposing candidates.** The subset of top drug-repurposing candidates that were manually evaluated against the published FXS and ASD pharmacology literature is summarized in **Table S7**. The validation set comprises eight compound-trait entries spanning several categories of prior evidence, including known target classes with prior evidence for the specific compound, novel compounds and mechanisms, and partially supported associations. Three entries were classified as "known target, novel compound". Two were classified as "novel" or "known association", and one was classified as "semi-novel". Because this literature check covers only a small, non-random subset of the full candidate set, these classifications provide contextual validation rather than an estimate of the overall novelty or success rate of the pipeline.

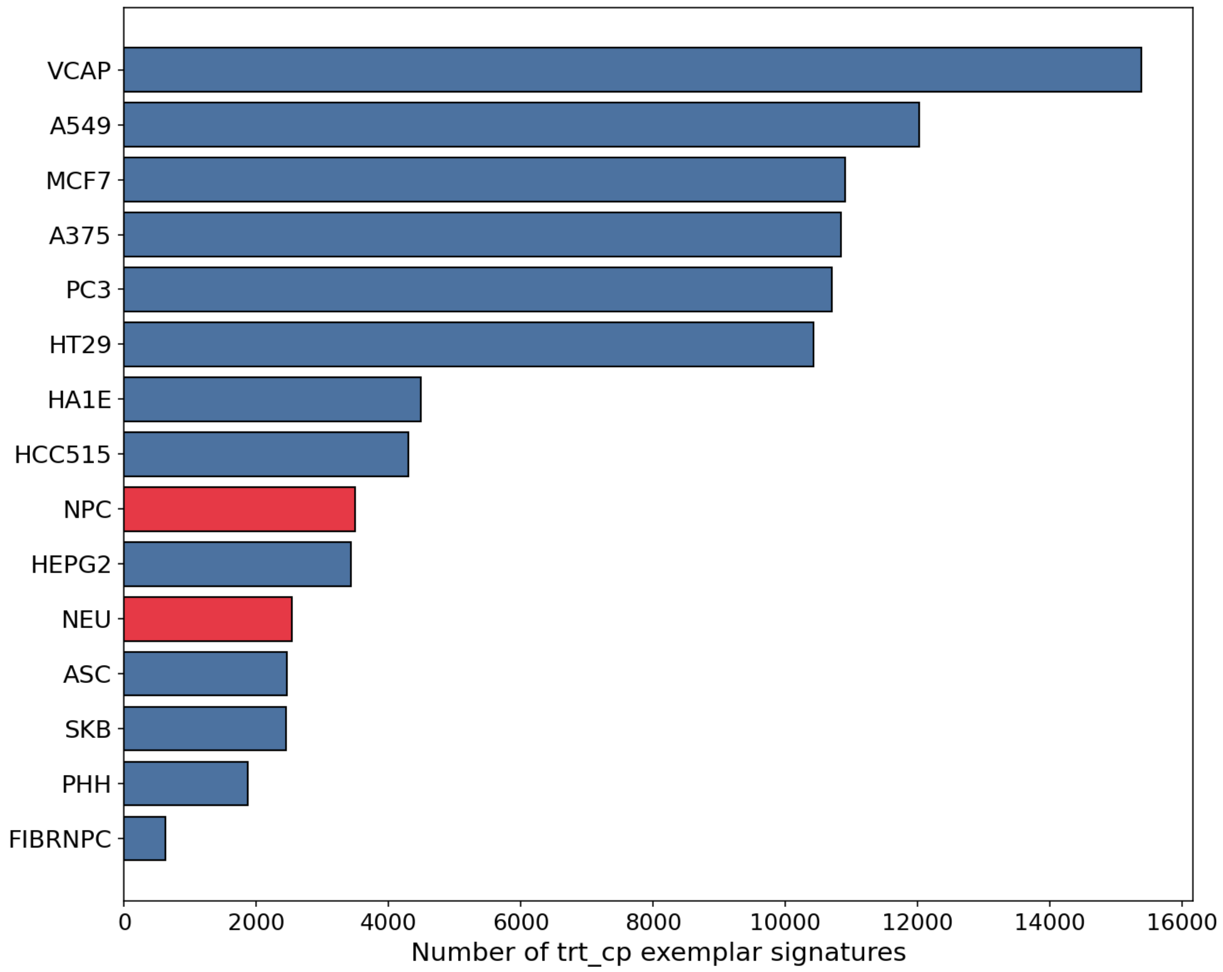


**Figure S18. Cell-line composition of the CMap LINCS L1000 exemplar signature library.** The top 15 cell lines are shown to illustrate the under-representation of neural-lineage cell lines (NPC and NEU) relative to the most represented cell lines. NPC (3,496 signatures) and NEU (2,538 signatures) rank ninth and eleventh among the top 15, respectively. The six most represented cell lines each contribute more than 10,000 exemplar signatures. VCAP, the most represented cell line, contributes 15,390 signatures, more than four times the combined number contributed by NPC and NEU. This composition illustrates that the compound screen was optimized for broad chemical-genomic coverage across predominantly non-neural, largely cancer-derived cell lines rather than for depth in neurally relevant cell types. This imbalance motivates the neural-lineage sensitivity analysis discussed in the main text.

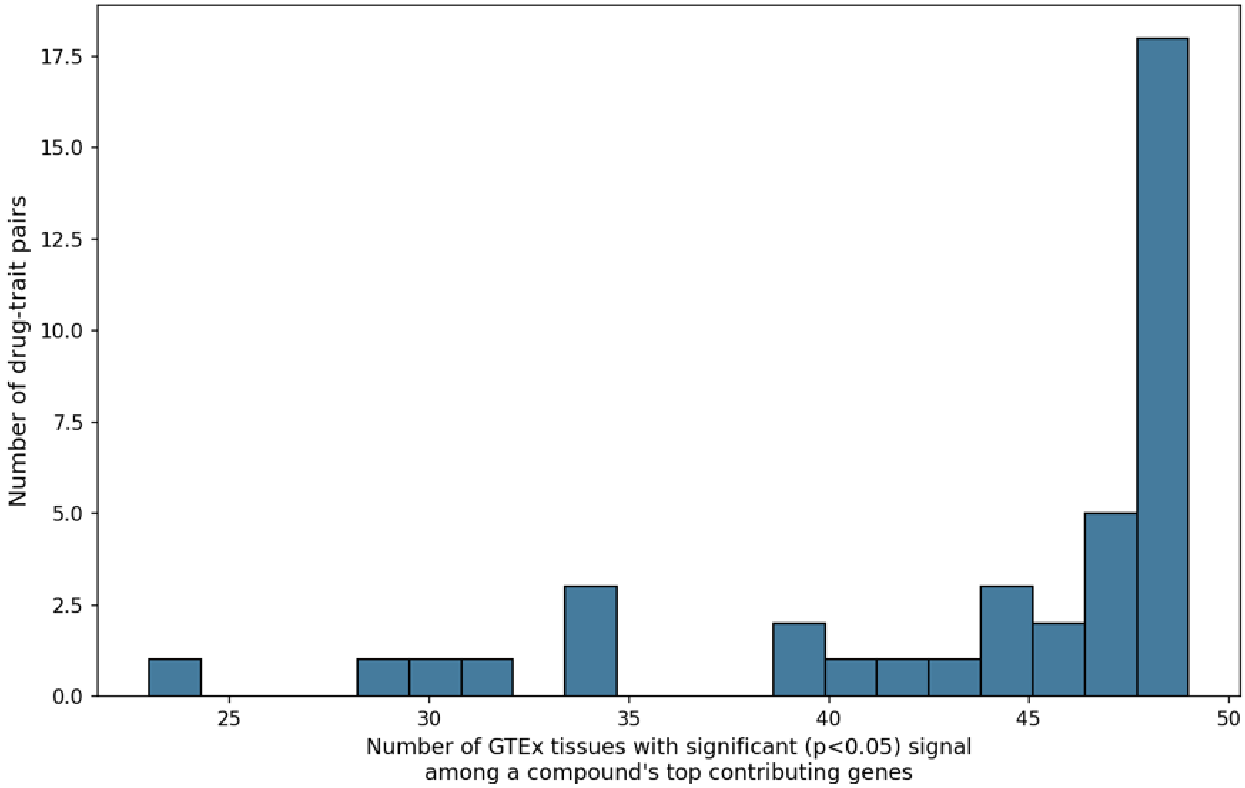


**Figure S19. Distribution of the number of GTEx tissues showing significant S-PrediXcan associations among top contributing genes.** The distribution is shown for the 40 compound-trait pairs included in the compound-trait-tissue heatmap analysis. Twenty-six of the 40 pairs (65%) show significant signal in at least 45 of the 49 tested GTEx tissues, with 18 pairs showing significance in 48 tissues. The number of tissues showing significant signal ranges down to 23 for idazoxan in ASD_vs_MDD. Thus, the breadth of tissue-tissue association varies substantially even among candidate compound-trait pairs meeting the same trait-level selection criteria. This prevalence of broad cross-tissue signal is consistent with genetically regulated expression reflecting regulatory effects shared across tissues for many of the genes contributing to these associations, rather than tissue-restricted regulation alone.

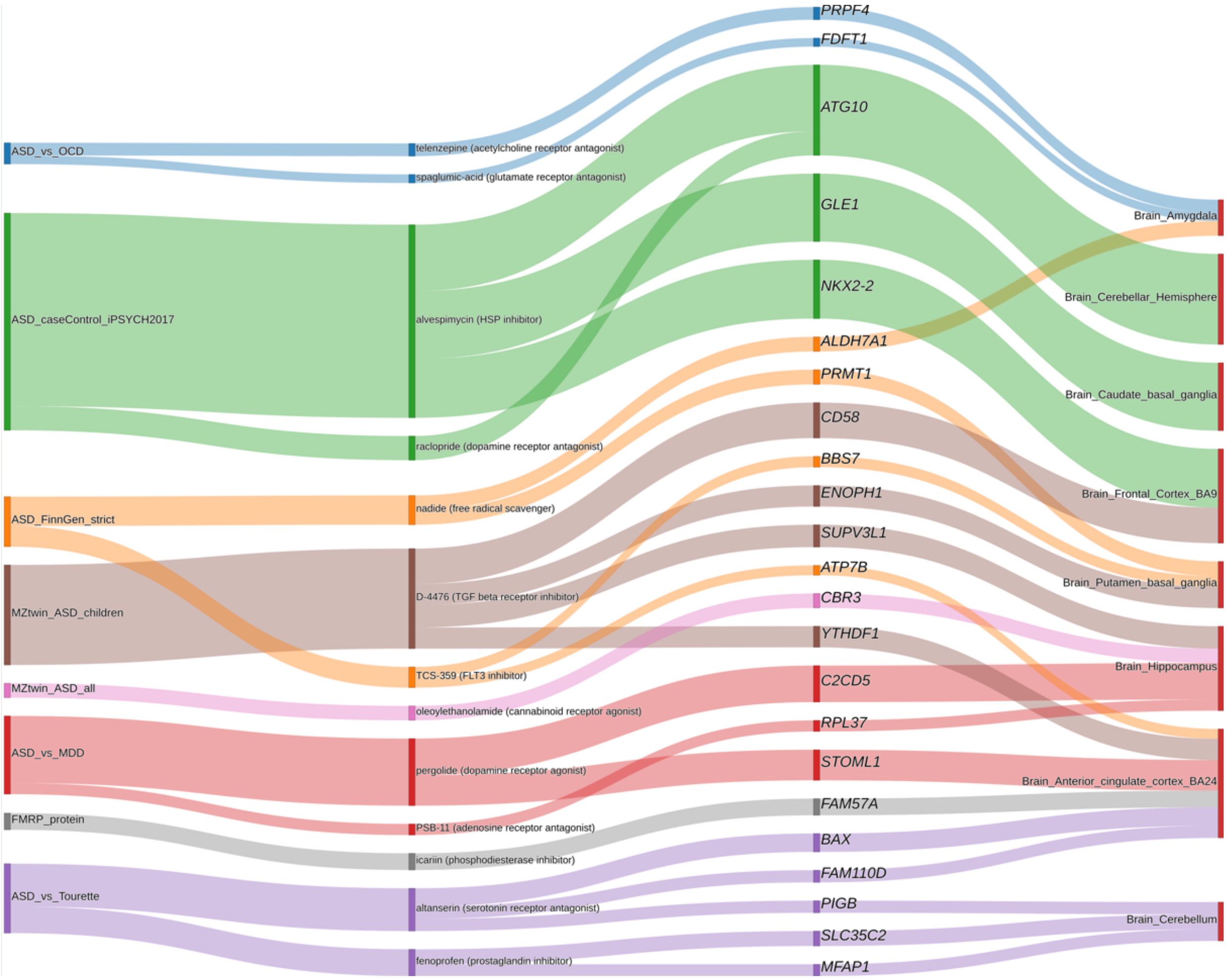


**Figure S20. Brain-tissue-restricted drug-trait-gene-tissue Sankey diagram.** The diagram shows the subset of links (24 of 80) in which a contributing gene's best-supporting GTEx tissue is a brain region, extending the full Sankey diagram shown in the main-text (**Figure 7**). All eight traits retain at least one brain-tissue link, and 13 of the 15 compounds shown in the full diagram remain represented. The eight brain regions represented include cortical, subcortical, and cerebellar structures, including Brain_Frontal_Cortex_BA9, Brain_Anterior_cingulate_cortex_BA24, Brain_Amygdala, Brain_Cerebellum, and Brain_Caudate_basal_ganglia. This brain-restricted view provides a complementary, hypothesis-generating perspective on the broader candidate set and does not imply that brain tissue is the sole or primary biologically relevant context.

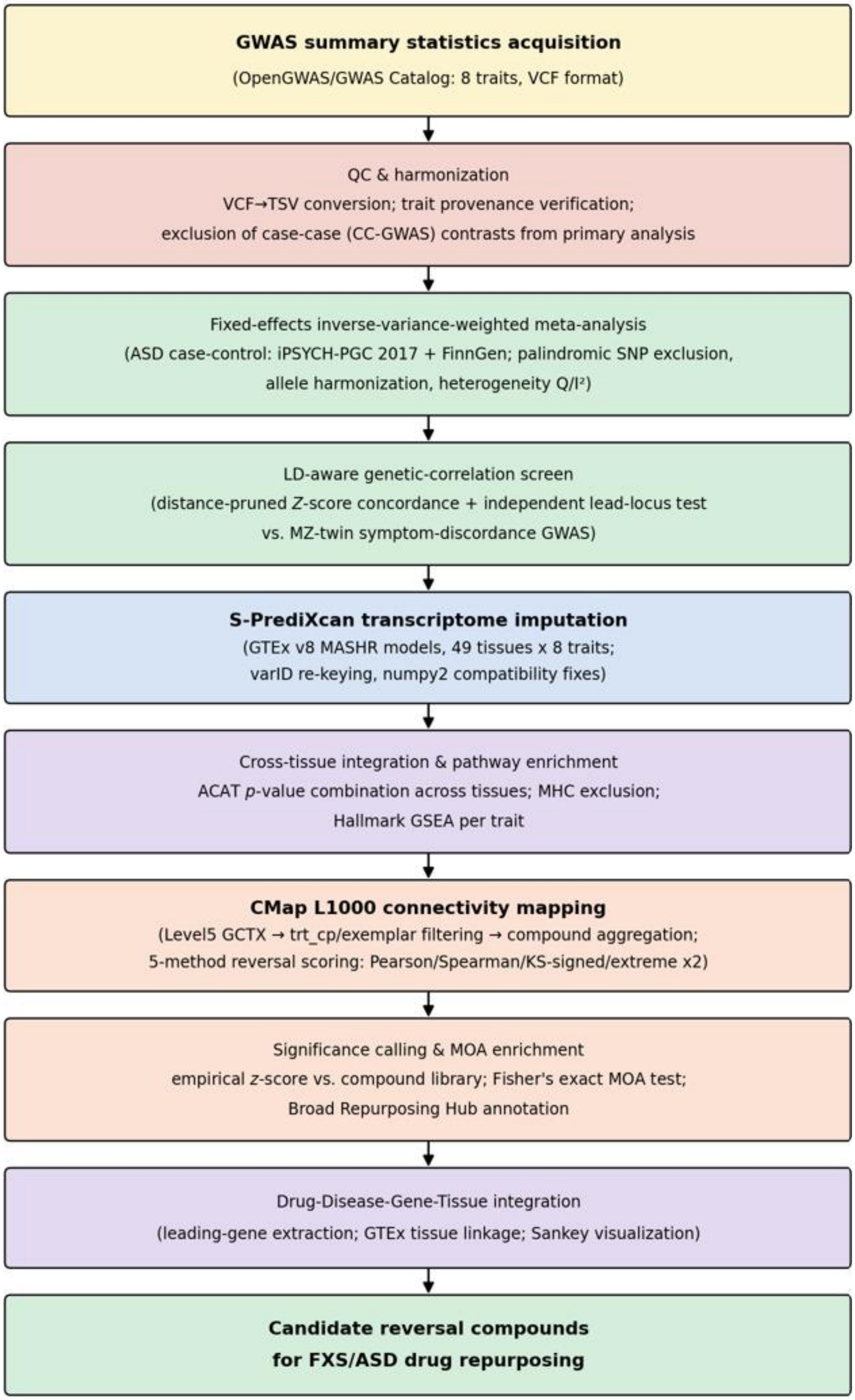
GWAS summary statistics acquisition
(OpenGWAS/GWAS Catalog: 8 traits, VCF format)
QC & harmonization
VCF→TSV conversion; trait provenance verification;
exclusion of case-case (CC-GWAS) contrasts from primary analysis
Fixed-effects inverse-variance-weighted meta-analysis
(ASD case-control: iPSYCH-PGC 2017 + FinnGen; palindromic SNP exclusion,
allele harmonization, heterogeneity $Q/I^2$)
LD-aware genetic-correlation screen
(distance-pruned $Z$-score concordance + independent lead-locus test
vs. MZ-twin symptom-discordance GWAS)
S-PrediXcan transcriptome imputation
(GTEx v8 MASHR models, 49 tissues x 8 traits;
varID re-keying, numpy2 compatibility fixes)
Cross-tissue integration & pathway enrichment
ACAT $p$-value combination across tissues; MHC exclusion;
Hallmark GSEA per trait
CMap L1000 connectivity mapping
(Level5 GCTX → trt_cp/exemplar filtering → compound aggregation;
5-method reversal scoring: Pearson/Spearman/KS-signed/extreme x2)
Significance calling & MOA enrichment
empirical $z$-score vs. compound library; Fisher's exact MOA test;
Broad Repurposing Hub annotation
Drug-Disease-Gene-Tissue integration
(leading-gene extraction; GTEx tissue linkage; Sankey visualization)
Candidate reversal compounds
for FXS/ASD drug repurposing

**Figure S21. Overview of the integrative analytical pipeline.** The pipeline integrates genetic datasets from GWAS summary statistics acquisition through ASD case-control meta-analysis, transcriptome imputation, cross-tissue integration, pathway analysis, and connectivity-based drug repurposing. The eight GWAS-scale datasets pass through a common quality-control and harmonization stage. The population case-control ASD datasets are then used for the dedicated meta-analysis, while all eight traits proceed through transcriptome imputation, cross-tissue integration, and pathway analysis. The resulting trait-level gene signatures are queried against the CMap L1000 compound library using five complementary connectivity metrics. Candidate compounds are filtered using the empirical reversal criteria and annotated by mechanism of action before integration into a unified compound, gene, tissue, and trait resource. The explicit exclusion of CC-GWAS contrasts from the primary population ASD meta-analysis and the use of an LD-aware analysis for genetic overlap with MZ-twin symptom-discordance data represent methodological safeguards against trait misclassification and inflation cause by non-independent variants.

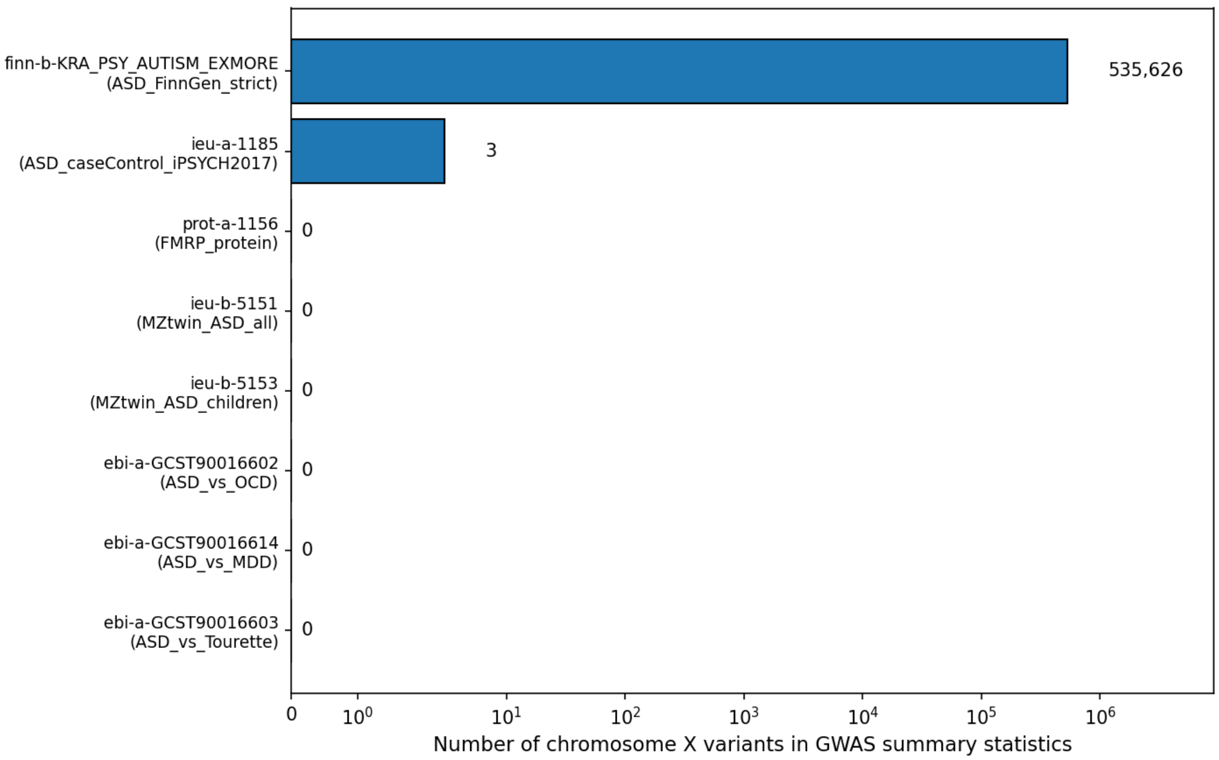


**Figure S22. Chromosome X variant and gene coverage across the eight input GWAS datasets and GTEx prediction models.** Chromosome X variant coverage varies substantially across the eight GWAS datasets. The GTEx v8 MASHR models used for S-PrediXcan contained no chromosome X genes, preventing transcriptome imputation at X-linked genes in the analysis. Six of the eight datasets contained no chromosome X variants, ASD_caseControl_iPSYCH2017 contained three, and ASD_FinnGen_strict provided substantial chromosome X coverage (535,626 variants). *FMR1*, the gene whose loss of function causes FXS, is located on chromosome X. Because the GTEx v8 prediction models contained no chromosome X genes, this pipeline could not evaluate a genetically regulated expression association for *FMR1* itself, regardless of chromosome X variant coverage in the GWAS datasets.

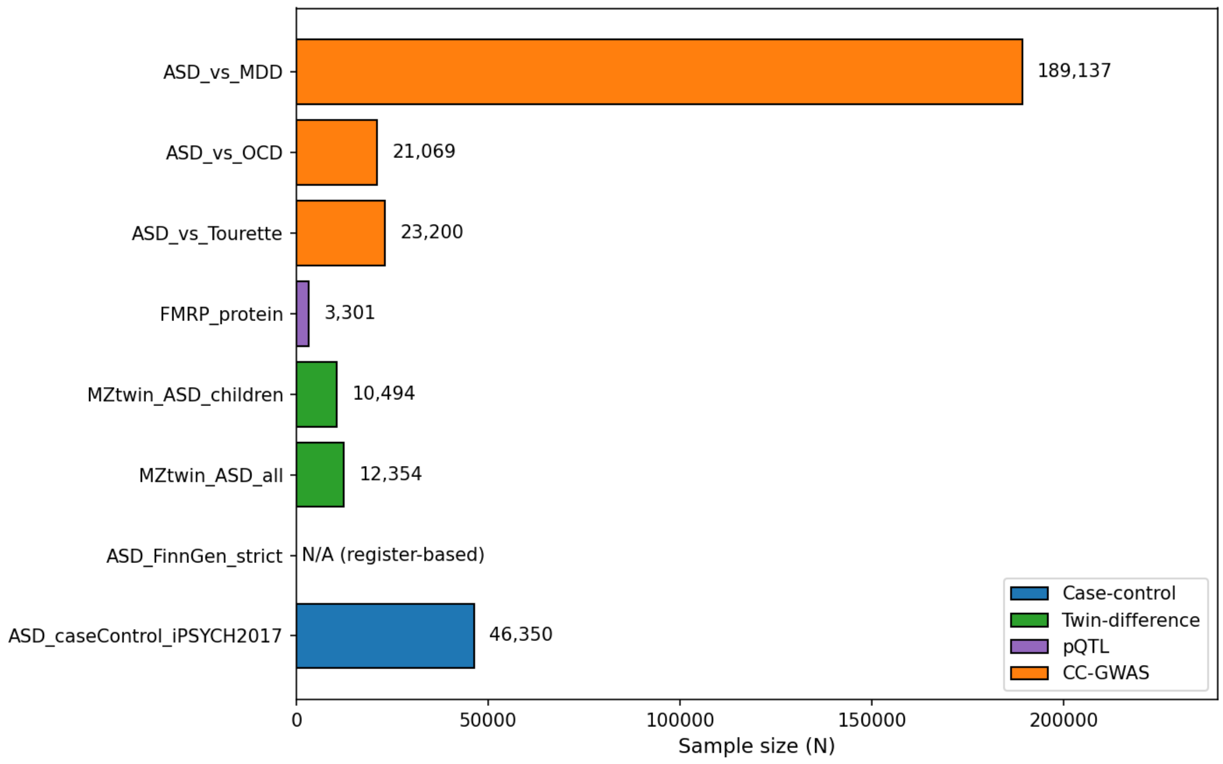


**Figure S23. Sample size and study design summary across the eight input GWAS datasets.** Sample sizes range from 3,301 (FMRP_protein, a plasma proteomic QTL study) to 189,137 (ASD_vs_MDD, the largest of the three CC-GWAS case-case contrast datasets), spanning four distinct study designs: case-control, within-twin-pair difference, plasma protein QTL, and case-case contrast. The three CC-GWAS datasets vary substantially in size despite sharing the same underlying ASD case set. Each compares the ASD case group with different psychiatric comparison group of different size. The implications of this sample overlap and study-design heterogeneity are considered in **Figure S6**.

### Supplementary Tables

**Table S1. Genome-wide significant SNPs from the corrected ASD case-control meta-analysis.** All 30 SNPs reaching genome-wide significance ($p < 5\times10^{-8}$) are located on chromosome 20 within a single 232 kb interval (21,182,388–21,414,037 bp, GRCh37). The lead SNP is rs910805 ($p = 4.6\times10^{-9}$). No genome-wide significant SNPs were identified on any other chromosome. The genome-wide significant associations therefore cluster within a single chromosome 20 region containing *NKX2-2* and *NKX2-4*.

| variant_id | chr | position | effect_allele | non_effect_allele | beta_meta | se_meta | *p*_meta | $I^2$ |
|---|---|---|---|---|---|---|---|---|
| rs910805 | 20 | 21248116 | A | G | -0.0928870008179416 | 0.015848137 | 4.5990139746409485e-09 | 38.716741781815 |
| rs1000177 | 20 | 21233198 | C | T | -0.0935520791355536 | 0.016144082 | 6.839373492296788e-09 | 45.557777355326 |
| rs12625304 | 20 | 21182388 | C | T | 0.1005839045922723 | 0.017927337 | 2.0157529458343093e-08 | 59.5406356000828 |
| rs6047339 | 20 | 21249538 | T | C | -0.0841665821538128 | 0.015065034 | 2.3118133859097927e-08 | 0.0 |
| rs112777342 | 20 | 21272136 | T | C | 0.0961536596941556 | 0.017219622 | 2.3510429508562232e-08 | 0.0 |
| rs73128919 | 20 | 21237133 | G | A | 0.1027517755121634 | 0.01850508 | 2.8141069417929973e-08 | 45.69170475030375 |
| rs73128955 | 20 | 21272052 | A | C | 0.101972520139832 | 0.018392904 | 2.9543115887647845e-08 | 10.607866546084209 |
| rs3848794 | 20 | 21343944 | A | G | -0.0954365044901669 | 0.017220506 | 2.9899090153664546e-08 | 0.0 |
| rs73128960 | 20 | 21277361 | A | G | 0.0954237920611322 | 0.017221093 | 3.0057823163324684e-08 | 29.899543431912424 |
| rs6075797 | 20 | 21348281 | A | G | -0.095339345176549 | 0.017220506 | 3.087792793939447e-08 | 0.0 |
| rs809220 | 20 | 21414037 | C | T | 0.0877037825646869 | 0.015842535 | 3.095120750230327e-08 | 5.009988752022576 |
| rs4815040 | 20 | 21344454 | A | G | -0.0952420772645924 | 0.017220506 | 3.188897035389002e-08 | 0.0 |
| rs6047401 | 20 | 21349729 | A | G | -0.0952420772645924 | 0.017220506 | 3.188897035389002e-08 | 0.0 |
| rs17736848 | 20 | 21278148 | A | G | 0.0952284736924628 | 0.017221093 | 3.2067328732904996e-08 | 29.706739481965656 |
| rs6047417 | 20 | 21362758 | G | T | -0.0951416871504005 | 0.017220506 | 3.2966121536665174e-08 | 0.0 |
| rs202847 | 20 | 21325222 | G | T | 0.0950682424209202 | 0.017220506 | 3.3776428304944244e-08 | 0.0 |
| rs73128966 | 20 | 21279983 | T | C | 0.0950542566683603 | 0.0172208 | 3.395112650679972e-08 | 32.99818396034861 |
| rs73130930 | 20 | 21332866 | G | A | 0.0950477495874949 | 0.017220506 | 3.40059491774385e-08 | 0.0 |
| rs6047413 | 20 | 21359787 | G | A | -0.094974764478656 | 0.0172208 | 3.4854414271283016e-08 | 0.0 |
| rs73128969 | 20 | 21280821 | G | A | 0.0949578723561425 | 0.0172208 | 3.504934443431782e-08 | 32.90929978590963 |
| rs202837 | 20 | 21312288 | G | A | 0.094950590273877 | 0.017220506 | 3.511488818133592e-08 | 0.0 |
| rs202840 | 20 | 21314079 | C | T | 0.094784837607651 | 0.0172208 | 3.7107990273165803e-08 | 0.0 |
| rs726025 | 20 | 21267360 | T | C | 0.094756527875047 | 0.017223699 | 3.76530487563875e-08 | 38.13497989392935 |
| rs73128953 | 20 | 21271737 | A | G | 0.0945779538572032 | 0.017221386 | 3.9765525850400735e-08 | 27.510981299753624 |
| rs2025811 | 20 | 21335113 | C | T | -0.1004788971180572 | 0.018300029 | 4.0047636279495374e-08 | 56.73285002060518 |
| rs202851 | 20 | 21330991 | T | C | 0.0942940503100829 | 0.017223125 | 4.3787911546609256e-08 | 36.473874084218686 |
| rs150548349 | 20 | 21262868 | G | A | 0.0940965093601702 | 0.017221678 | 4.6594110317498695e-08 | 0.0 |
| rs6035864 | 20 | 21366260 | C | T | -0.0940806834958038 | 0.017222548 | 4.6908928240696135e-08 | 40.68632008189071 |
| rs202835 | 20 | 21302885 | A | G | 0.0939544936739892 | 0.017222548 | 4.8884990678750775e-08 | 41.64700432576248 |
| rs202829 | 20 | 21298446 | T | C | 0.0939544936739892 | 0.017222548 | 4.8884990678750775e-08 | 41.64700432576248 |

**Table S2. Independent lead loci from the corrected ASD case-control meta-analysis.** This table contains 67 lead loci reaching $p < 10^{-5}$, with one variant retained per one-Mb physical window according to the pruning described in the Methods. The displayed table is truncated to the first 30 of 67 rows and the complete table is provided in the accompanying data file.

| variant_id | chromosome | position | *p*_meta | *z*_meta |
|---|---|---|---|---|
| rs910805 | 20 | 21248116 | 4.5990139746409485e-09 | -5.861067344953658 |
| rs1452075 | 3 | 62481063 | 5.2185846469576185e-08 | 5.443697385900878 |
| rs7000772 | 8 | 10795613 | 6.807587379595925e-08 | 5.396176785715179 |
| rs112635299 | 14 | 94838142 | 1.46301375413545e-07 | 5.257154529543762 |
| rs2391769 | 1 | 96978961 | 2.094911610376588e-07 | 5.190712252976642 |
| rs2388334 | 6 | 98591622 | 2.5357489644725053e-07 | 5.155041402303637 |
| rs2224274 | 20 | 14760747 | 3.150616430911402e-07 | 5.114207040596956 |
| rs9366877 | 6 | 11730878 | 3.6172339547800093e-07 | -5.08807129369419 |
| rs45595836 | 10 | 16691399 | 4.341365972683121e-07 | 5.053345906775606 |
| rs325485 | 5 | 103995368 | 5.191781632317432e-07 | -5.019086328114423 |
| rs147118670 | 19 | 37531025 | 5.931234088366388e-07 | 4.993442855141835 |
| rs325481 | 5 | 104000752 | 7.473111012897213e-07 | -4.948647075654624 |
| rs16879023 | 6 | 16753147 | 7.947926301993984e-07 | -4.936641780227362 |
| rs35404050 | 12 | 73196902 | 8.14124807364153e-07 | 4.931950666999971 |
| rs71375338 | 17 | 44332793 | 8.346431685991858e-07 | 4.927087657060896 |
| rs11249905 | 8 | 9042454 | 1.0179738404884748e-06 | -4.888131778141578 |
| rs28729902 | 9 | 76179384 | 1.1955542430895527e-06 | 4.856372958986429 |
| rs292441 | 18 | 55872558 | 1.3265029899563818e-06 | -4.835741375291667 |
| rs17517971 | 13 | 110828119 | 1.3443911526580117e-06 | 4.8330764708899725 |
| rs13188074 | 5 | 113801423 | 1.410421515244518e-06 | -4.823526311527852 |
| rs76397219 | 8 | 60390318 | 1.7403076064098214e-06 | 4.781455234099825 |
| rs2635182 | 5 | 92255166 | 1.7705480518371243e-06 | 4.777991566432729 |
| rs12759704 | 1 | 99090942 | 1.9000623615904497e-06 | -4.763773011782564 |
| rs79940520 | 3 | 191838169 | 1.9579853894235207e-06 | 4.757713072235641 |
| rs7783557 | 7 | 71646872 | 1.966958386995947e-06 | -4.756789752481917 |
| rs11185408 | 1 | 104792257 | 2.2470871230514475e-06 | -4.729829284454049 |
| rs29227 | 6 | 29580588 | 2.27871649729728e-06 | 4.726990742551177 |
| rs118058985 | 15 | 93941693 | 2.323670005231707e-06 | 4.723020863970449 |
| rs35923829 | 7 | 114972493 | 2.35674553811934e-06 | 4.720146750948451 |
| rs77691144 | 13 | 66970212 | 2.429600326915437e-06 | 4.7139502281929335 |

**Table S3. Top 20 cross-tissue ACAT gene associations per trait.** The table contains 160 gene-trait entires. The displayed table is truncated to the first rows, and the complete table is provided in the accompanying data file. Twenty-three genes, including *XRN2*, appear among the top 20 for two traits. These repeated appearances provide a descriptive measure of cross-trait overlap among highly ranked genes and do not represent formal evidence of shared genetic architecture. The table ranks the strongest nominal gene associations separately for each trait, with each ACAT *p*- value interpreted in relation to the trait-specific number of genes tested.

| trait | gene_name | *p*_acat | mean_*z*score | *n*_tissues_tested | best_tissue | best_*p* |
|---|---|---|---|---|---|---|
| ASD_caseControl_iPSYCH2017 | XRN2 | 2.2563970003552927e-07 | -1.605639525424553 | 40 | Colon_Sigmoid | 1.9017438380008466e-08 |
| ASD_caseControl_iPSYCH2017 | RP11-6N13.1 | 3.9647994815039677e-07 | 5.0706400871276855 | 1 | Testis | 3.964799481356577e-07 |
| ASD_caseControl_iPSYCH2017 | KIZ | 2.4516638821858727e-06 | 3.564230154513513 | 6 | Esophagus_Mucosa | 8.227164767574704e-07 |
| ASD_caseControl_iPSYCH2017 | PINX1 | 3.981406839104196e-06 | -2.2132956050716834 | 43 | Artery_Coronary | 1.870978021388433e-07 |
| ASD_caseControl_iPSYCH2017 | NKX2-4 | 4.618515130949685e-06 | 0.0 | 2 | Pituitary | 4.618515130883317e-06 |
| ASD_vs_MDD | NKX2-4 | 6.795346249477419e-06 | 4.440892098500626e-16 | 2 | Testis | 6.7953462494600815e-06 |
| ASD_caseControl_iPSYCH2017 | STH | 7.381171698683797e-06 | 4.4823899269104 | 4 | Brain_Cortex | 7.381171698711373e-06 |
| ASD_caseControl_iPSYCH2017 | GABBR1 | 9.303084686940188e-06 | 2.051864442101838 | 33 | Brain_Nucleus_accumbens_basal_ganglia | 1.869025038189912e-06 |
| ASD_caseControl_iPSYCH2017 | CRHR1 | 9.765888787760524e-06 | 1.3091859864917668 | 22 | Artery_Coronary | 7.994538430255471e-06 |
| ASD_caseControl_iPSYCH2017 | SPPL2C | 1.0598202894451791e-05 | 3.981057602538274 | 10 | Brain_Putamen_basal_ganglia | 6.617630474256149e-06 |
| ASD_caseControl_iPSYCH2017 | LRRC37A2 | 1.1475793212922802e-05 | 3.809166829160898 | 36 | Cells_Cultured_fibroblasts | 5.7906491849070696e-06 |
| FMRP_protein | PPP3CA | 1.3630767760453022e-05 | -0.6576128873373551 | 33 | Lung | 8.261773732055864e-07 |
| ASD_vs_MDD | XRN2 | 1.586938576270125e-05 | 1.2097858496213345 | 37 | Skin_Not_Sun_Exposed_Suprapubic | 7.452665054980452e-07 |
| ASD_vs_OCD | LIN52 | 1.6972089791789102e-05 | 2.5445473370077565 | 34 | Testis | 9.85415706114323e-07 |
| ASD_caseControl_iPSYCH2017 | PLEKHM1 | 1.7556020720399523e-05 | 0.0475492635306592 | 14 | Brain_Cerebellum | 7.579505244710511e-06 |
| ASD_vs_MDD | ADO | 1.82961872056131e-05 | -2.449304414647868 | 29 | Artery_Aorta | 6.7953462494600815e-06 |
| ASD_vs_MDD | ISLR | 2.049556204786196e-05 | 2.307500110673053 | 28 | Heart_Left_Ventricle | 1.7079796241978608e-05 |
| FMRP_protein | TNRC18 | 2.0990746345483924e-05 | 2.2531660957769915 | 11 | Brain_Hypothalamus | 2.106074792209358e-06 |
| ASD_caseControl_iPSYCH2017 | MAPT | 2.2009294754543607e-05 | -0.0778281650339536 | 44 | Whole_Blood | 1.2390992599842568e-06 |
| ASD_FinnGen_strict | NOVA1 | 2.2210891064688987e-05 | -3.36652671653492 | 15 | Spleen | 5.344236800292955e-06 |
| ASD_caseControl_iPSYCH2017 | RP1L1 | 2.434657724537104e-05 | -2.59496783890116 | 9 | Uterus | 3.324986340735485e-06 |
| ASD_caseControl_iPSYCH2017 | NSF | 2.5036545560019885e-05 | -1.6111650109291076 | 10 | Ovary | 8.345242717221576e-06 |
| ASD_vs_MDD | RP11-247C2.2 | 2.5765988058368272e-05 | 3.4008364321942826 | 11 | Lung | 1.7079796241978642e-05 |
| ASD_caseControl_iPSYCH2017 | NUDT21 | 2.6768816688838012e-05 | 3.466887966445919 | 46 | Muscle_Skeletal | 5.88610601022317e-06 |
| ASD_caseControl_iPSYCH2017 | ARL17A | 2.6861509633269165e-05 | 2.5352756121635265 | 45 | Brain_Hippocampus | 7.985271169733048e-06 |
| ASD_vs_OCD | BBOF1 | 2.732287157364288e-05 | 2.6990971444695395 | 45 | Colon_Transverse | 5.481795478511283e-06 |
| ASD_caseControl_iPSYCH2017 | ADTRP | 2.752632442099001e-05 | -0.3663666683621352 | 25 | Spleen | 1.1063658130641912e-06 |
| MZtwin_ASD_all | KRT80 | 2.8181069206489884e-05 | 0.4040378870814455 | 39 | Minor_Salivary_Gland | 7.23896613202871e-07 |
| ASD_vs_MDD | ISLR2 | 2.829476763743788e-05 | 3.7637746749648926 | 16 | Lung | 1.7079796241978642e-05 |

| trait | gene_name | *p*_acat | mean_*z*score | *n*_tissues_tested | best_tissue | best_*p* |
|---|---|---|---|---|---|---|
| MZtwin_ASD_all | RP13-516M14.1 | 3.1608689076367824e-05 | 3.850459572864719 | 47 | Uterus | 1.7653851104319043e-05 |

**Table S4. Full Hallmark GSEA results for all 50 pathways across eight traits.** The table contains 400 trait-pathway combinations. The displayed table is truncated to the first 30 rows and the complete table is provided in the accompanying data file. Seven trait-pathway combinations reach FDR < 0.25, spanning five distinct Hallmark pathways: interferon alpha response, cholesterol homeostasis, protein secretion, reactive oxygen species pathway, and oxidative phosphorylation. FDR-adjusted *q*-values for these seven combinations range from 0.039 to 0.226.

| Trait | Term | NES | NOM *p*-val | FDR *q*-val | Lead_genes |
|---|---|---|---|---|---|
| ASD_FinnGen_strict | HALLMARK_COMPLEMENT | -1.37580883 | 0.00994036 | 0.59050445 | HPCAL4;USP8;USP15;LAP3;GZMK;DYRK2;RCE1;KYNU;GCA;MMP12;CD55;GNB4;GATA3;PRCP;RAF1;HSPA5;PIK3CG;MT3;PPP2CB;ADAM9;SCG3;ADRA2B;PIK3CA;SERPINE1;GZMA;PPP4C;CCL5;APOBEC3F;GNAI3;CR1;GZMB;C1QC;PIK3R5;SIRT6;KLKB1;C1S;PCSK9;CASP10;C4BPB;LYN;PDGFB;DUSP5 |
| ASD_FinnGen_strict | HALLMARK_PROTEIN_SECRETION | 1.45197940 | 0.01550388 | 0.15911564 | RAB2A;ADAM10;SGMS1;ATP6V1B1;SSPN;MON2;TMED10;TSPAN8;SOD1;AP3S1;COPE;SNAP23;ARFIP1;CLCN3;RER1;RAB14;SEC22B |
| ASD_FinnGen_strict | HALLMARK_REACTIVE_OXYGEN_SPECIES_PATHWAY | 1.50889843 | 0.01879699 | 0.16770283 | GPX3;SCAF4;TXN;NQO1;MSRA;LAMTOR5;SOD1;STK25;SOD2;PRDX6;GLRX2 |
| ASD_FinnGen_strict | HALLMARK_MYOGENESIS | 1.33800031 | 0.01890359 | 0.30307740 | GPX3;REEP1;ATP2A1;ACTA1;NQO1;KCNH2;CDH13;SMTN;TEAD4;RYR1;PICK1;SSPN;COL6A2;COL1A1;NOS1;MYL6B;PKIA;MYL1;MRAS;PLXNB2;BHLHE40;EPHB3;MYH3;SLN;KIFC3;COL6A3;SPTAN1;SPARC;MYLK;MYO1C;FKBP1B;LAMA2;MEF2A;FDPS;PVALB;PYGM;ACTC1;AK1;TAGLN;KLF5;CLU;IGF1;TNNI1;ITGB4;TNNT3;TNNI2;CD36;LARGE1;PTP4A3;CHRNB1;CNN3 |
| ASD_FinnGen_strict | HALLMARK_HEME_METABOLISM | -1.30946234 | 0.02 | 0.53808111 | ELL2;BMP2K;USP15;OSBP2;BPGM;MKRN1;PICALM;RNF19A;KAT2B;IGSF3;SLC2A1;DCAF11;XPO7;LRP10;SLC4A1;ABCG2;HEBP1;FOXO3;GLRX5;HBZ;NARF;HAGH;CDC27;ATP6V0A1;LPIN2;RNF123;NR3C1;CDR2;ACSL6;HTRA2;SPTA1;TCEA1;PGLS;DCUN1D1;MFHAS1;EPOR;AHSP;GMPS |
| ASD_FinnGen_strict | HALLMARK_SPERMATOGENESIS | -1.21549865 | 0.09297521 | 0.60731949 | MAST2;THEG;NPHP1;ART3;PEBP1;ODF1;MLF1;EZH2;PSMG1;TNNI3;IDE;MAP7;HSPA4L;PRKAR2A;CLVS1;YBX2;SCG3;AGFG1;HBZ;VDAC3;IP6K1;DNAJB8;TUBA3C;DPEP3;ZC2HC1C;CNIH2;RFC4;TEKT2;PACRG;ACE;SLC12A2;BUB1;PHKG2;ALOX15 |
| ASD_FinnGen_strict | HALLMARK_UNFOLDED_PROTEIN_RESPONSE | -1.20188182 | 0.10472279 | 0.45252226 | WFS1;SPCS1;EXOSC9;DNAJC3;EDEM1;POP4;HSPA5;SEC11A;BAG3;EIF2S1;EIF4G1;ERO1A;HSPA9;HERPUD1;EIF4A2;NFYB;DDIT4;SHC1;ATF3;BANF1;HSP90B1;SEC31A;CEBPB;KDELR3;STC2;EIF4A3 |
| ASD_FinnGen_strict | HALLMARK_COAGULATION | 1.20513728 | 0.10825688 | 0.57524091 | ANXA1;ITIH1;MMP3;BMP1;HNF4A;OLR1;MMP10;DPP4;GP1BA;TFPI2;CPN1;KLF7;MMP8;TMPRSS6;PF4;LTA4H;C1QA;SPARC;F13B;PREP;CD9;MST1;F10;PROS1;CLU;APOC1;F12;MASP2 |
| ASD_FinnGen_strict | HALLMARK_NOTCH_SIGNALING | 1.28052811 | 0.11776860 | 0.37556341 | ST3GAL6;FZD1;PSENEN;FBXW11;HEYL;ARRB1 |
| ASD_FinnGen_strict | HALLMARK_BILE_ACID_METABOLISM | -1.21292265 | 0.12007874 | 0.49535114 | DIO2;CYP8B1;PEX13;NR0B2;ALDH8A1;SLC27A5;CYP7B1;TFCP2L1;ABCA3;RXRG;NR3C2;PEX26;ABCA1;ACSL5;SLC27A2;ABCG4;SULT2B1;PEX11G;ALDH1A1;PNPLA8;PEX19 |
| ASD_FinnGen_strict | HALLMARK_PANCREAS_BETA_CELLS | -1.25069906 | 0.13663366 | 0.60138477 | SPCS1;SCGN;IAPP;SEC11A;PAX6;CHGA;ISL1;INSM1;G6PC2;FOXO1;NEUROG3;SRP14;NEUROD1 |
| ASD_FinnGen_strict | HALLMARK_GLYCOLYSIS | -1.14768502 | 0.14845361 | 0.58061325 | ALDH7A1;GOT1;PMM2;HS2ST1;MERTK;SLC25A10;PAXIP1;HSPA5;GPC1;PPP2CB;DPYSL4;HDLBP;SDC1;GFPT1;LHX9;KIF20A;AKR1A1;SLC25A13;MIOX;GLRX;GMPPB;ERO1A;NASP;ELF3;P4HA1;ADORA2B;CD44;B4GALT4;UGP2 |
| ASD_FinnGen_strict | HALLMARK_HYPOXIA | -1.13357478 | 0.15975104 | 0.44975272 | ETS1;PFKFB3;BTG1;PPARGC1A;SLC2A1;ENO3;HSPA5;GPC1;DPYSL4;HDLBP;RBPJ;FOXO3;GLRX;SERPINE1;ERO1A;PLAC8;P4HA1;ADORA2B;PPP1R3C;DTNA;GAPDH;RORA;HS3ST1;UGP2;ACKR3;NR3C1;SULT2B1;ERRFI1;BCL2;CDKN1B;PDGFB;DDIT4;TIPARP;ATF3;HK2;HMOX1;PPP1R15A;ANXA2;CA12;KDELR3;B4GALNT2;IRS2;STC2;PRDX5;AKAP12 |

| Trait | Term | NES | NOM *p*-val | FDR *q*-val | Lead_genes |
| --- | --- | --- | --- | --- | --- |
| ASD_FinnGen_strict | HALLMARK_INTERFERON_GAMMA_RESPONSE | 1.12476327 | 0.16195859 | 0.88178686 | FCGR1A;NAMPT;RBCK1;SAMD9L;IL2RB;CXCL10;PSME1;RTP4;SSPN;IFI44;MVP;IFIH1;PSME2;SPPL2A;SOD2;NLRC5;CXCL11;IFIT2;SAMHD1;CASP1;NUP93;LATS2;VCAM1;MX1;SOCS1;USP18;ZNFX1;IL6;ADAR;LGALS3BP;CD38;IFI27;MTHFD2;IFI30;IL7;STAT4;CD69;TRIM21;ST8SIA4;CASP4;IRF7;IL4R;SERPING1;VAMP5;PDE4B |
| ASD_FinnGen_strict | HALLMARK_ANDROGEN_RESPONSE | -1.14399323 | 0.18473896 | 0.51916419 | PTPN21;ELL2;ITGAV;ANKH;PTK2B;MERTK;MAP7;ACTN1;UAP1;SCD;ARID5B;TMEM50A;GNAI3;ELOVL5;INPP4B;HERC3;VAPA;ZMIZ1;PIAS1 |
| ASD_FinnGen_strict | HALLMARK_PI3K_AKT_MTOR_SIGNALING | -1.13572535 | 0.20654397 | 0.49126278 | PIK3R3;GRK2;SLC2A1;MAPK8;DAPP1;CAB39;RAF1;E2F1;PRKAR2A;UBE2N;ECSIT;CFL1;ACTR2;PAK4;RPS6KA1;ACTR3;NFKBIB;TIAM1;TSC2;CDKN1B;AP2M1;PIN1;ARHGDIA;ARF1;HSP90B1;GNA14;THEM4;CDK2;CLTC |
| ASD_FinnGen_strict | HALLMARK_APOPTOSIS | 1.09551625 | 0.23204420 | 0.80340768 | FASLG;GPX3;ANXA1;CDC25B;SMAD7;BTG3;PMAIP1;SOD1;SOD2;PSEN1;SATB1;CASP1;BNIP3L;CREBBP;SPTAN1;BIK;PLCB2;AIFM3;NEFH;IL6;PDCD4;CCND2;SLC20A1;CASP2;CD38;BCL2L11;CLU;CD2;GSR;BMP2;CD69 |
| ASD_FinnGen_strict | HALLMARK_MTORC1_SIGNALING | -1.08817704 | 0.23886640 | 0.56253934 | PIK3R3;NUPR1;IMMT;POLR3G;MCM4;SLC2A1;DAPP1;EDEM1;GOT1;TUBG1;PSMG1;TUBA4A;ETF1;HSPA5;TMEM97;SEC11A;PSMD13;PSMC2;BCAT1;GLRX;ERO1A;ARPC5L;RDH11;PSMC6;P4HA1;GAPDH;SCD;HSPA9;ACTR2;ACTR3;ELOVL5;NFKBIB |
| ASD_FinnGen_strict | HALLMARK_INTERFERON_ALPHA_RESPONSE | 1.10457391 | 0.24091778 | 0.86362627 | SAMD9L;TMEM140;CXCL10;PSME1;RTP4;IFI44;IFIH1;PSME2;PARP9;CXCL11;IFIT2;CASP1;MX1;USP18;ADAR;LGALS3BP;UBA7;IFI27;IFI30;IL7;GBP2;TRIM21 |
| ASD_FinnGen_strict | HALLMARK_KRAS_SIGNALING_UP | -1.08515133 | 0.25053079 | 0.52761292 | ETS1;EVI5;TSPAN13;BPGM;ITGBL1;CIDEA;ST6GAL1;ANKH;NR0B2;TMEM158;ETV1;CMKLR1;MYCN;MAP7;USP12;PTPRR;BTBD3;SCG3;GLRX;ERO1A;F13A1;SPON1;ZNF277;PECAM1;ITGA2;GADD45G;F2RL1;IRF8;DNMBP;PRKG2;CD37;MMD;ID2;CPE;APOD;ACE;RETN;LY96;TPH1;PPP1R15A;KLF4;GPNMB;CBX8;PLAU;AKAP12 |
| ASD_FinnGen_strict | HALLMARK_CHOLESTEROL_HOMEOSTASIS | 1.07370626 | 0.30101010 | 0.82055400 | NFIL3;ACAT2;FBXO6;CYP51A1;MVK;DHCR7;PNRC1;TRIB3;SEMA3B;CD9;FDPS;CLU;LDLR |
| ASD_FinnGen_strict | HALLMARK_XENOBIOTIC_METABOLISM | 1.03949562 | 0.34086629 | 0.91509170 | NQO1;ANGPTL3;DDC;ITIH1;PAPSS2;CNDP2;HNF4A;ATP2A2;CDO1;PDLIM5;DCXR;CROT;ABCC3;GABARAPL1;IL1R1;PINK1;UGDH;TMBIM6;SLC6A6;ARG2;GAD1;AKR1C2;PDK4;SAR1B;F10;PROS1;MCCC2;SERTAD1;SHMT2;CYP26A1;IGF1;APOE;CD36;NPC1;ALDH3A1;GSR;IGFBP1;ABCD2;BLVRB;SERPINA6;GCNT2;ALAS1;ALDH2 |
| ASD_FinnGen_strict | HALLMARK_MYC_TARGETS_V2 | 1.03905062 | 0.37298387 | 0.83373838 | GNL3;MRTO4;DUSP2;PA2G4;SORD;TCOF1;RABEPK;NOLC1 |
| ASD_FinnGen_strict | HALLMARK_WNT_BETA_CATENIN_SIGNALING | -1.00924708 | 0.42181070 | 0.77988283 | AXIN1;DKK1;TP53;CTNNB1;RBPJ |
| ASD_FinnGen_strict | HALLMARK_EPITHELIAL_MESENCHYMAL_TRANSITION | -0.98781905 | 0.45030426 | 0.81743677 | CADM1;EDIL3;ITGAV;TNFRSF11B;DKK1;NOTCH2;BASP1;GJA1;SGCB;FMOD;GPC1;TIMP3;SDC1;MMP1;TPM4;SERPINE1;GREM1;SGCG;NTM;CD44;SCG2;ITGA2;P3H1;GADD45B;WNT5A;COL7A1;CAP2;MFAP5;PTX3;COLGALT1;TNFRSF12A;COL16A1;FGF2;ID2;FERMT2;TPM2;HTRA1;PCOLCE;BDNF;RGS4;IL15;SFRP4;CDH2;PLOD2;FOXC2;FBLN5;NT5E |
| ASD_FinnGen_strict | HALLMARK_TNFA_SIGNALING_VIA_NFKB | 0.99509770 | 0.46572581 | 0.98980028 | NAMPT;NFIL3;CCL4;BTG3;CXCL10;OLR1;TNFSF9;NR4A3;PDLIM5;DUSP2;IER5;IFIH1;SOD2;ICOSLG;CXCL11;IFIT2;SLC16A6;BHLHE40;PNRC1;MSC;GPR183;CD83;NFAT5;REL;TRIP10;STAT5A;IL6;BCL2A1;AREG;TNIP1;EDN1;CXCL2;YRDC;INHBA;LDLR;IL7R;BCL6;BMP2;CD69;FOS;GFPT2;SERPINB8;MAP2K3;SDC4;PDE4B;DNAJB4;PMEPA1;TNFAIP8;CCL2;JUNB;SQSTM1;HES1;FJX1 |
| ASD_FinnGen_strict | HALLMARK_KRAS_SIGNALING_DN | 0.98638198 | 0.47159091 | 0.96036680 | ITIH3;KRT15;CD207;VPS50;ATP6V1B1;RYR1;TLX1;SHOX2;NOS1;GP1BA;THRB;SLC12A3;ZBTB16;PRODH;MX1;MYOT;SLC5A5;KMT2D;GTF3C5;TENM2;TSHB;GPR19;ACTC1;EDN1 |
| ASD_FinnGen_strict | HALLMARK_MYC_TARGETS_V1 | 0.98157370 | 0.49632353 | 0.91601536 | GNL3;ODC1;RUVBL2;USP1;EIF4H;CANX;CCT4;CDC45;SRSF1;HNRNPA3;PA2G4;RANBP1;AP3S1;SF3B3;POLD2;EIF4E;PPM1G;SYNCRIP;NCBP1;NME1;PSMD8;PSMA6;ERH;COPS5;VDAC1;NOLC1;HDAC2;ILF2;C1QBP;RPLP0 |

| Trait | Term | NES | NOM *p*-val | FDR *q*-val | Lead_genes |
|---|---|---|---|---|---|
| ASD_FinnGen_strict | HALLMARK_PEROXISOME | -0.93055726 | 0.60599572 | 0.94943126 | ACOT8;CADM1;UGT2B17;PEX13;ESR2;IDE;ALB;FIS1;RDH11;RXRG;NUDT19;CNBP;ACSL5;SLC27A2;ELOVL5;SULT2B1;ALDH1A1;VPS4B;PEX14;CRAT |
| ASD_FinnGen_strict | HALLMARK_ADIPOGENESIS | -0.94058997 | 0.60652174 | 0.96557864 | SULT1A1;PFKFB3;CYC1;IMMT;CIDEA;BCKDHA;FABP4;GPX4;SDHB;SLC25A10;MAP4K3;TST;PEMT;UQCRC1;POR;CPT2;GPAT4;TKT;ABCA1;PPM1B;COQ5;UBQLN1;LAMA4;DLAT;DECR1;PTCD3;COX6A1;TOB1;NKIRAS1;PEX14;CRAT;REEP5;IFNGR1;MGLL;CMBL;PHYH;APLP2;ORM1;RETN;ACAA2 |

**Table S5. Compounds identified as significant reversal candidates in three or more traits.** The table contains 52 compound-trait entries corresponding to 17 compounds. The displayed table is truncated to first 30 rows and the complete table is provided in the accompanying data file. Sixteen compounds are shared across exactly three traits, and one compound, BRD-K12148366, is shared across four traits. Empirical $z$-scores for individual compound-trait pairs range from −2.12 to −3.02, indicating that cross-trait recurrence is observed across a range of reversal scores rather than being confined to the most extreme individual signals.

| compound | trait | ks_z | pearson_all | spearman_all | ks_signed_mean |
|---|---|---|---|---|---|
| BRD-K12148366 | ASD_caseControl_iPSYCH2017 | -2.689942405582322 | -0.009649313 | -0.00948407 | -0.1918027411997318 |
| BRD-K12148366 | ASD_vs_OCD | -2.0735935309683464 | -0.010425302 | -0.010754102 | -0.1413289447203278 |
| BRD-K12148366 | MZtwin_ASD_all | -2.284424811040516 | -0.02054371 | -0.012927385 | -0.1864799491167068 |
| BRD-K12148366 | MZtwin_ASD_children | -2.17552642847807 | -0.022409761 | -0.016826665 | -0.1798557260632515 |
| BRD-K25631986 | ASD_vs_MDD | -2.560763892362006 | -0.00547558 | -0.0040797936 | -0.1845236855149269 |
| BRD-K25631986 | MZtwin_ASD_all | -2.3546713183488834 | -0.014202386 | -0.008921119 | -0.1919097747504711 |
| BRD-K25631986 | MZtwin_ASD_children | -2.133339681719484 | -0.01468834 | -0.010404108 | -0.176603305786848 |
| BRD-K26087938 | ASD_vs_OCD | -2.393590657651109 | -0.012291295 | -0.008997972 | -0.1633061710596084 |
| BRD-K26087938 | ASD_vs_Tourette | -2.305962750354221 | -0.016941333 | -0.020395797 | -0.1511272360086441 |
| BRD-K26087938 | FMRP_protein | -2.3306351572220207 | -0.03143937 | -0.02357056 | -0.1902303495444357 |
| BRD-K35743871 | ASD_FinnGen_strict | -2.68111753842652 | -0.009818496 | -0.007763019 | -0.2035447673201561 |
| BRD-K35743871 | ASD_vs_OCD | -2.20556145755513 | -0.01768058 | -0.018407466 | -0.150392429202795 |
| BRD-K35743871 | ASD_vs_Tourette | -2.1988187346881958 | -0.01151744 | -0.008747068 | -0.1443509483933448 |
| BRD-K40910787 | ASD_vs_MDD | -2.119084876593129 | -0.01505868 | -0.010002257 | -0.1528957766294479 |
| BRD-K40910787 | FMRP_protein | -2.110281828710461 | -0.013487552 | -0.0058705094 | -0.1722865231633186 |
| BRD-K40910787 | MZtwin_ASD_all | -2.0256827938803927 | -0.0067708422 | -0.0030149438 | -0.1664800360798835 |
| BRD-K43939579 | ASD_vs_Tourette | -2.4144179198372417 | -0.0076359473 | -0.008338593 | -0.1579864471107721 |
| BRD-K43939579 | MZtwin_ASD_all | -2.018562529366372 | -0.013581354 | -0.008741685 | -0.1659296628832817 |
| BRD-K43939579 | MZtwin_ASD_children | -2.186966849982852 | -0.013547964 | -0.006780372 | -0.1807377343177795 |
| BRD-K46284961 | ASD_caseControl_iPSYCH2017 | -2.5644498059897005 | -0.008722365 | -0.0040275 | -0.1829030321836471 |
| BRD-K46284961 | MZtwin_ASD_all | -2.2258326773014487 | -0.0051549785 | -0.0021080475 | -0.181950968503952 |
| BRD-K46284961 | MZtwin_ASD_children | -2.235801419371659 | -0.0060663573 | -0.002146476 | -0.1845026736855506 |
| BRD-K60807183 | ASD_caseControl_iPSYCH2017 | -2.3146419032537966 | -0.0022197065 | -0.0023208775 | -0.1651871058940887 |
| BRD-K60807183 | ASD_vs_OCD | -2.1875463708620955 | -0.012577193 | -0.015740499 | -0.149155162975192 |
| BRD-K60807183 | ASD_vs_Tourette | -2.3531218871482094 | -0.006791968 | -0.008557839 | -0.1541097999215126 |
| BRD-K70243592 | ASD_vs_Tourette | -2.112366983462911 | -0.008323359 | -0.0040214066 | -0.1388833361566066 |
| BRD-K70243592 | MZtwin_ASD_all | -2.062901919621273 | -0.021485517 | -0.018995121 | -0.1693569529652595 |
| BRD-K70243592 | MZtwin_ASD_children | -2.536440794123608 | -0.021553345 | -0.01828978 | -0.2076807017028331 |
| BRD-K75961033 | ASD_caseControl_iPSYCH2017 | -2.0430436553065565 | -0.022493543 | -0.012679764 | -0.1459258475899696 |
| BRD-K75961033 | MZtwin_ASD_all | -2.5346990585644 | -0.021172583 | -0.021584498 | -0.2058253312110901 |

**Table S6. Literature validation status of manually evaluated drug-repurposing candidates.** The table summarizes six manually evaluated top drug-repurposing candidates and their published FXS and ASD pharmacology evidence. The six compounds span four literature-validation categories and five mechanistic classes. Two compounds, icariin and GS-39783, have target classes that have reached at least Phase 2 clinical evaluation in FXS specifically. providing an external benchmark against which the remaining, less-studied candidates can be compared. Two compounds, spaglumic-acid and valsartan, had no prior compound-specific FXS or ASD literature identified. The manual evaluation was limited to a small, selected subset of candidates and therefore provides contextual validation rather than evidence for a broader candidate set.

| Compound | Trait(s) | Mechanism of action | Literature status | Supporting notes |
|---|---|---|---|---|
| oleoylethanolamide | MZtwin_ASD_all / MZtwin_ASD_children | cannabinoid receptor agonist | Known association | Aran et al. 2019 (Molecular Autism) reported significantly reduced circulating OEA in children with ASD; endocannabinoid system dysregulation is an established ASD mechanism. |
| spaglumic-acid | ASD_vs_OCD | glutamate receptor antagonist | Semi-novel | Compound itself not previously studied in ASD; its target class (NAAG/mGluR3/NMDA modulation) falls within the well-established glutamate hypothesis of autism. |
| valsartan | ASD_FinnGen_strict / MZtwin_ASD_all | angiotensin receptor antagonist | Novel | No ASD/FXS-specific literature identified; known mechanisms are neuroprotection, BDNF modulation, and anti-oxidative-stress effects in other neurological/cardiovascular contexts. |
| L-368899 | ASD_vs_OCD | oxytocin receptor antagonist | Known target, novel compound | Oxytocin receptor signalling is among the most extensively studied ASD drug targets (multiple agonist trials); this specific antagonist compound has not been studied in ASD. |
| icariin | FMRP_protein | phosphodiesterase inhibitor | Known target, novel compound | PDE4 inhibition is the most clinically advanced FXS drug target (BPN14770/zatolmilast, Phase 2/3 trials, FRAXA-funded); icariin itself has not been studied for FXS/ASD. |
| GS-39783 | ASD_vs_OCD | GABA receptor modulator | Known target, novel compound | GABA-B receptor agonism (arbaclofen) reached Phase 2/3 FXS clinical trials; GS-39783 itself has not been studied for FXS/ASD. |

**Table S7. Full mechanism-of-action enrichment results for all 473 elementary MOA terms tested.** The displayed table is truncated to the first 30 rows, and the complete table is provided in the accompanying data file. Of the 473 tested MOA terms, 135 (28.5%) contain at least one significant reversal candidate, while the remainder contain none. The minimum FDR-corrected *q*-value across all 473 terms is 0.265. Approximately 45% of terms (215 of 473) have fewer than 10 annotated compounds.

| MOA | sig_count | total_count | odds_ratio | *p*-value | FDR |
|---|---|---|---|---|---|
| anticancer agent | 3 | 24 | 10.309734513274336 | 0.0042625743983055 | 0.2652126102672896 |
| lysophospholipid receptor antagonist | 2 | 8 | 23.97209985315712 | 0.0050463287365869 | 0.2652126102672896 |
| interferon inducer | 2 | 8 | 23.97209985315712 | 0.0050463287365869 | 0.2652126102672896 |
| other antifungal | 2 | 8 | 23.97209985315712 | 0.0050463287365869 | 0.2652126102672896 |
| vasodilator | 2 | 8 | 23.97209985315712 | 0.0050463287365869 | 0.2652126102672896 |
| Kir6 channel (KATP) activator | 2 | 8 | 23.97209985315712 | 0.0050463287365869 | 0.2652126102672896 |
| KATP activator | 2 | 8 | 23.97209985315712 | 0.0050463287365869 | 0.2652126102672896 |
| anti-HCVE2 | 2 | 8 | 23.97209985315712 | 0.0050463287365869 | 0.2652126102672896 |
| glucose dependent insulinotropic receptor agonist | 2 | 8 | 23.97209985315712 | 0.0050463287365869 | 0.2652126102672896 |
| T-type calcium channel blocker | 3 | 32 | 7.462007934086055 | 0.0096332859091997 | 0.4556544235051497 |
| toll-like receptor agonist | 2 | 16 | 10.268722466960352 | 0.0201112494479989 | 0.8647837262639562 |
| potassium channel blocker | 4 | 80 | 3.8023391812865497 | 0.0248915947441631 | 0.90567110107609 |
| angiotensin receptor antagonist | 4 | 80 | 3.8023391812865497 | 0.0248915947441631 | 0.90567110107609 |
| phosphodiesterase inhibitor | 7 | 248 | 2.105154947478599 | 0.0572686102909404 | 1.0 |
| sodium channel blocker | 6 | 216 | 2.065470852017937 | 0.0798842877749285 | 1.0 |
| differentiation inducer | 1 | 8 | 10.228070175438596 | 0.1054404202630559 | 1.0 |
| IP1 prostacyclin receptor agonist | 1 | 8 | 10.228070175438596 | 0.1054404202630559 | 1.0 |
| glycosylated protein precursor | 1 | 8 | 10.228070175438596 | 0.1054404202630559 | 1.0 |
| aryl hydrocarbon receptor antagonist | 1 | 8 | 10.228070175438596 | 0.1054404202630559 | 1.0 |
| chemoreceptor agonist | 1 | 8 | 10.228070175438596 | 0.1054404202630559 | 1.0 |
| growth factor receptor activator | 1 | 8 | 10.228070175438596 | 0.1054404202630559 | 1.0 |
| nedd activating enzyme inhibitor | 1 | 8 | 10.228070175438596 | 0.1054404202630559 | 1.0 |
| NTPDase inhibitor | 1 | 8 | 10.228070175438596 | 0.1054404202630559 | 1.0 |
| enkephalinase inhibitor | 1 | 8 | 10.228070175438596 | 0.1054404202630559 | 1.0 |
| oxytocin receptor antagonist | 1 | 8 | 10.228070175438596 | 0.1054404202630559 | 1.0 |
| prostaglandin inhibitor | 1 | 8 | 10.228070175438596 | 0.1054404202630559 | 1.0 |
| vasopressin receptor antagonist | 1 | 8 | 10.228070175438596 | 0.1054404202630559 | 1.0 |
| integrin antagonist | 1 | 8 | 10.228070175438596 | 0.1054404202630559 | 1.0 |
| NFKB pathway inhibitor | 1 | 8 | 10.228070175438596 | 0.1054404202630559 | 1.0 |
| guanylate cyclase activator | 1 | 8 | 10.228070175438596 | 0.1054404202630559 | 1.0 |

**Table S8. Summary of the eight GWAS-scale datasets used in this study.** All eight datasets are in GRCh37/hg19 coordinates and are predominantly of European ancestry. They span four study designs: case-control, within-twin-pair difference, plasma protein QTL, and case-case contrast. Sample sizes range from 3,301 (FMRP_protein) to 189,137 (ASD_vs_MDD).

| Trait short name | GWAS ID | Sample size | First author/cohort | Source journal/ repository | Study design | Sample size | Genome build | Ancestry |
|---|---|---|---|---|---|---|---|---|
| ASD_caseControl_iPSYCH2017 | ieu-a-1185 | 18,381 / 27,969 | Grove et al. 2019 | Nature Genetics | Case-control | 18,381 cases / 27,969 controls | GRCh37 | European |
| ASD_FinnGen_strict | finn-b-KRA_PSY_AUTISM_EXMORE | N/A (register-based) | FinnGen R5 | FinnGen | Case-control | Register-based (N not disclosed) | GRCh37 | European (Finnish) |
| MZtwin_ASD_all | ieu-b-5151 | 12,354 pairs | Assary et al. 2025 | Within Families Consortium | MZ-twin within-pair difference | 12,354 pairs | GRCh37 | European |
| MZtwin_ASD_children | ieu-b-5153 | 10,494 pairs | Assary et al. 2025 | Within Families Consortium | MZ-twin within-pair difference | 10,494 pairs | GRCh37 | European |
| FMRP_protein | prot-a-1156 | 3,301 | Sun et al. 2018 | Nature (INTERVAL) | Plasma protein QTL | 3,301 | GRCh37 | European |
| ASD_vs_Tourette | ebi-a-GCST90016603 | 23,200 | Peyrot & Robinson 2021 | Nature Genetics (CC-GWAS) | Case-case contrast | 23,200 | GRCh37 | Mixed |
| ASD_vs_OCD | ebi-a-GCST90016602 | 21,069 | Peyrot & Robinson 2021 | Nature Genetics (CC-GWAS) | Case-case contrast | 21,069 | GRCh37 | Mixed |
| ASD_vs_MDD | ebi-a-GCST90016614 | 189,137 | Peyrot & Robinson 2021 | Nature Genetics (CC-GWAS) | Case-case contrast | 189,137 | GRCh37 | Mixed |
